# Meta-analytic evidence for a novel hierarchical model of conceptual processing

**DOI:** 10.1101/2022.11.05.515278

**Authors:** Philipp Kuhnke, Marie C. Beaupain, Johannes Arola, Markus Kiefer, Gesa Hartwigsen

## Abstract

Conceptual knowledge plays a pivotal role in human cognition. Grounded cognition theories propose that concepts consist of perceptual-motor features represented in modality-specific perceptual-motor cortices. However, it is unclear whether conceptual processing consistently engages modality-specific areas. Here, we performed an activation likelihood estimation (ALE) meta-analysis across 212 neuroimaging experiments on conceptual processing related to 7 perceptual-motor modalities (action, sound, visual shape, motion, color, olfaction-gustation, and emotion). We found that conceptual processing consistently engages brain regions also activated during real perceptual-motor experience of the same modalities. In addition, we identified multimodal convergence zones that are recruited for multiple modalities. In particular, the left inferior parietal lobe (IPL) and posterior middle temporal gyrus (pMTG) are engaged for three modalities: action, motion, and sound. These “trimodal” regions are surrounded by “bimodal” regions engaged for two modalities. Our findings support a novel model of the conceptual system, according to which conceptual processing relies on a hierarchical neural architecture from modality-specific to multimodal areas up to an amodal hub.

## Introduction

Conceptual knowledge about objects, people, and events in the world is crucial for core human cognitive abilities, such as object recognition and use, problem solving, as well as language comprehension (Lambon Ralph, 2013; van Elk et al., 2014). Therefore, a central question in cognitive neuroscience has been how concepts are represented and processed in the human brain.

According to the traditional view in cognitive science—amodal theories—concepts can be considered as entirely abstract, amodal symbols (Fodor, 1975; Pylyshyn, 1984). Thus, under this view, the conceptual system is completely separated from the modality-specific systems for perception and action. In contrast, grounded theories of cognition propose that concepts consist of perceptual and motor features represented in modality-specific perceptual-motor brain regions (Barsalou, 2008; Kiefer and Barsalou, 2013). For example, concepts like ‘dog’ are assumed to comprise visual shape, color, and motion features represented in visual brain areas, sound features represented in auditory areas, action features in somatomotor regions, as well as olfaction and emotion features in olfactory-gustatory and emotion-related brain regions, respectively (Binder and Desai, 2011; Fernandino et al., 2016b, 2016a). While a common terminology is still lacking in the field, we refer to “perceptual-motor modalities” as the brain’s major input and output channels of perception and action (cf. Kuhnke et al., 2021). Note that these modalities do not simply correspond to the senses (hence the term “perceptual-motor” and not “sensory”) as they include channels of internal perception (e.g. emotion) as well as motor action (Kiefer and Harpaintner, 2020). Moreover, some senses (e.g. vision) may contain several sub-modalities (e.g. color, shape, and motion). We call brain regions “modality-specific” if they represent information related to a single perceptual-motor modality (Kiefer and Pulvermüller, 2012). Grounded cognition theories are mainly supported by neuroimaging studies demonstrating that conceptual processing related to a certain perceptual-motor modality activates the corresponding modality-specific areas (for reviews, see Hauk and Tschentscher, 2013; Kiefer and Harpaintner, 2020; Kiefer and Pulvermüller, 2012; Meteyard et al., 2012).

In addition to modality-specific regions, previous evidence suggests an involvement of “cross-modal convergence zones” which integrate multiple different modality-specific features into more abstract conceptual representations (Binder et al., 2009; Binder and Desai, 2011; Lambon Ralph et al., 2016). We recently proposed a distinction among cross-modal convergence zones between “multimodal” regions which retain modality-specific information, and “amodal” regions which completely abstract away from modality-specific input (Kuhnke et al., 2021, 2020b). Multimodal regions seem to include the left inferior parietal lobe (IPL), posterior middle temporal gyrus (pMTG), and medial prefrontal cortex (mPFC) (Fernandino et al., 2016b, 2016a; Kuhnke et al., 2021, 2020b). In contrast, the anterior temporal lobe (ATL) appears to act as an amodal hub of the conceptual system (Jefferies, 2013; Lambon Ralph et al., 2016; Patterson et al., 2007).

However, several issues remain open. First, it is unknown whether conceptual processing consistently engages modality-specific perceptual-motor regions across neuroimaging studies. Several studies failed to find modality-specific perceptual-motor activity during conceptual tasks (e.g. Bedny et al., 2008; Postle et al., 2008; Raposo et al., 2009) and the involvement of modality-specific regions in conceptual processing remains controversial (Kompa, 2021; Kompa and Mueller, 2020; Mahon, 2015; Mahon and Caramazza, 2008). Second, it is unknown which modalities robustly overlap in multimodal convergence zones. In particular, it is unclear which cross-modal regions are multimodal (rather than amodal), as well as *how many* and *which* modalities are integrated in each multimodal area. A better understanding of the overlap and dissociation within and between areas would advance the current knowledge of the neural basis of conceptual processing.

Coordinate-based meta-analyses allow us to formally assess the consistency of functional activations across numerous neuroimaging experiments, capitalizing on the common reporting of activation coordinates in standard space (Fox et al., 2014; Yarkoni et al., 2010). However, previous meta-analyses have mainly investigated conceptual processing in general (i.e. regardless of perceptual-motor content; Binder et al., 2009; Visser et al., 2010) or executive control during concept retrieval (Jackson, 2021; Noonan et al., 2013). No meta-analysis has systematically tested the grounded cognition hypothesis that conceptual processing related to a certain perceptual-motor modality recruits the respective modality-specific perceptual-motor regions. Two meta-analyses focused on action-related conceptual processing (Binder et al., 2009; Watson et al., 2013) and identified consistent engagement of left pMTG and anterior supramarginal gyrus (aSMG). However, neither study tested for overlap with real action execution (also see Giacobbe et al., 2022). Likewise, a previous meta-analysis on emotion-related concepts (Desai et al., 2018) did not test for overlap with real emotion perception. In addition, no meta-analysis has simultaneously investigated conceptual processing related to multiple different modalities.

To address these issues, we performed an activation likelihood estimation (ALE) meta-analysis across a total of 212 neuroimaging experiments (3893 participants, 3101 activation foci) on conceptual processing related to 7 perceptual-motor modalities (action, sound, visual shape, motion, color, olfaction-gustation, and emotion). We also performed ALE meta-analyses on real perception or action in each modality (studies identified using the BrainMap database; total of 1582 experiments, 21,349 participants, 39,221 foci) and tested for overlap with conceptual processing. This design allowed us to investigate several modalities simultaneously, thereby dissociating modality-specific brain areas and multimodal convergence zones. To test modality specificity, we directly contrasted the meta-analytic maps between conceptual modalities. Finally, to identify multimodal convergence zones, we investigated the conjunctions between all modalities, and analyzed how many and which modalities overlap in each multimodal area.

Following grounded cognition theories, we hypothesized that conceptual processing should engage brain regions that are also activated during real perceptual-motor experience of the same modalities. These perceptual-motor regions should exhibit a high modality specificity. Regarding multimodal convergence zones, we predicted that several modalities should overlap in the “multimodal” left IPL, pMTG and mPFC, but not in the “amodal” ATL.

## Materials and Methods

### Literature search

We performed a systematic literature search using the PubMed, Web of Science, and BrainMap databases, as well as manual reference tracing (through reviews and original research articles). Table S1 shows a checklist following the guidelines for neuroimaging meta-analyses by Müller et al. (2018), which includes the literature search terms. Figure 1 presents a flowchart following the PRISMA guidelines (Page et al., 2021).

**Figure 1.**
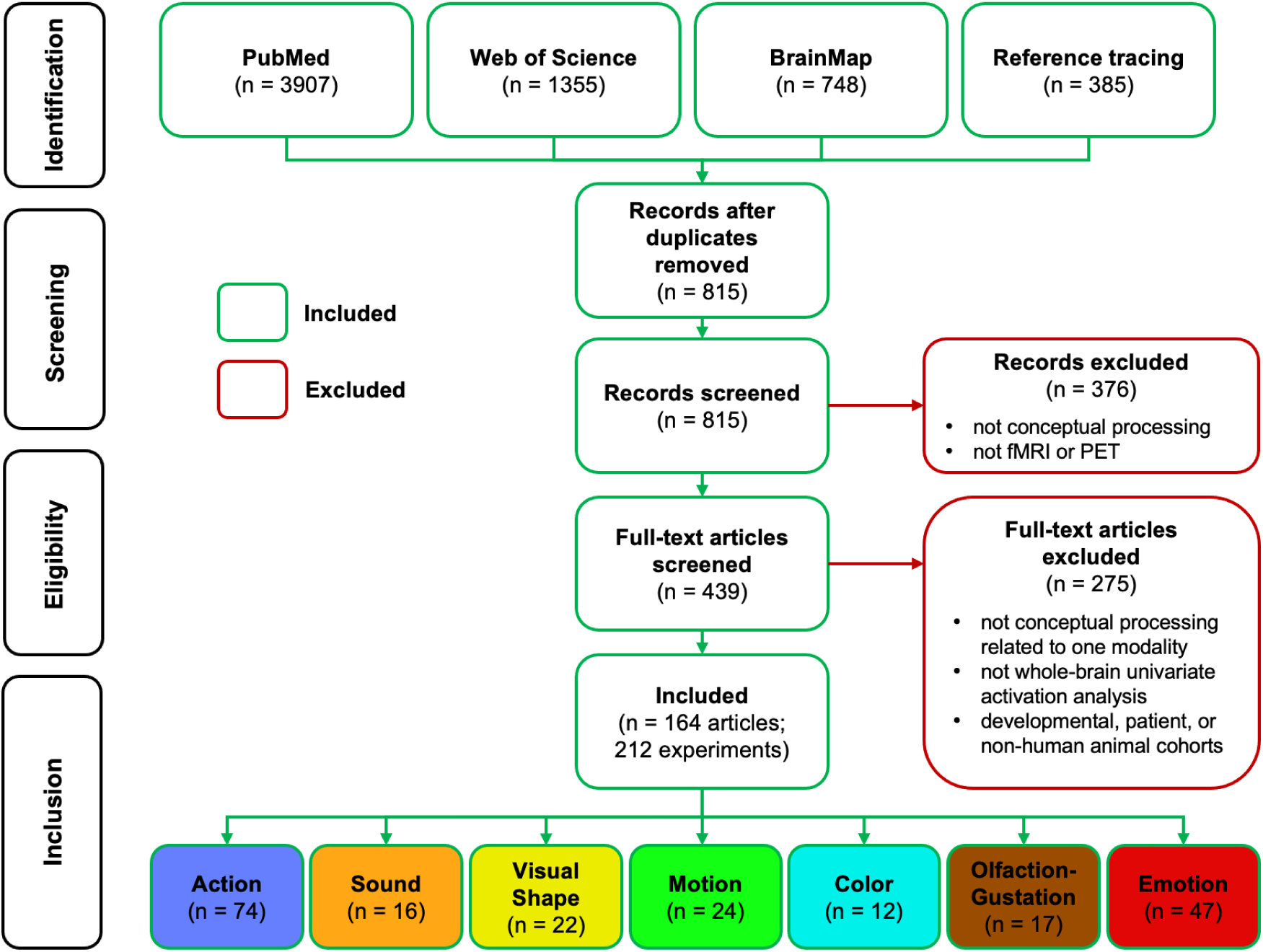
Flowchart illustrating the inclusion-exclusion process leading to the datasets included in the meta-analyses (following the PRISMA guidelines; Page et al., 2021). Before the final “inclusion” stage, *n* represents the number of articles. At the “inclusion” stage, *n* represents the number of experiments per modality (where one article could contain multiple experiments).

We included functional magnetic resonance imaging (fMRI) and positron emission tomography (PET) studies that reported peak coordinates from voxel-wise, whole-brain activation analyses in Montreal Neurological Institute (MNI) or Talairach (TAL) space. Therefore, we excluded non-whole-brain (e.g. region-of-interest), non-voxel-wise (e.g. multivariate), and non-activation (e.g. connectivity) analyses. Study participants had to be healthy, right-handed human adults; developmental, patient, and non-human animal cohorts were excluded. In studies using language stimuli, only native-speaker samples were included.

Appropriate experiments asked participants to perform implicit or explicit conceptual tasks on pictures, sounds, words, sentences, or stories. We selectively included activation contrasts that targeted conceptual processing related to one of the 7 perceptual-motor modalities (action, sound, visual shape, color, motion, olfaction-gustation, or emotion). To this end, we carefully ensured that the relationship between stimuli and perceptual-motor modality was purely conceptual-semantic (not perceptual). Therefore, we manually excluded all contrasts with a potential stimulus difference in the same modality as the targeted conceptual modality (e.g. visual-shape stimuli for visual-shape related conceptual processing, such as the contrast [animal pictures > tool pictures]; Chao et al., 1999). Conversely, we selectively included contrasts that targeted conceptual processing related to a different perceptual-motor modality than the stimulus modality (e.g. visual stimuli for sound-related conceptual processing, as in the contrast [sound-related > non-sound-related written words]; e.g. Kiefer et al., 2008; Kuhnke et al., 2020b). Experiments involving mental imagery were only included if they required conceptual knowledge retrieval (e.g. Zvyagintsev et al., 2013).

To enable tests for modality specificity and overlap across modalities, every contrast was assigned to only one modality. To completely distinguish the modalities action and motion, action was defined as object manipulation, whereas motion subsumed all other (non-object-directed) movements (cf. Fernandino et al., 2016; van Elk et al., 2014). Contrasts that could not be assigned to only one modality were excluded.

Our primary analysis included both high-level and low-level contrasts to maximize statistical power, which is considered crucial for the sensitivity and specificity of neuroimaging meta-analyses (Eickhoff et al., 2016; Müller et al., 2018). However, only high-level contrasts between different experimental conditions (e.g. action words > abstract words) can isolate brain activity specific to conceptual processing. Contrasts against a low-level baseline (e.g. action words > fixation) may yield concomitant activation for non-conceptual processes (e.g. orthographic or phonological processing). Therefore, to test the robustness of our meta-analytic results to baseline differences, we complemented our primary analysis with a supplementary analysis that excluded low-level contrasts (cf. Diveica et al., 2021).

Table 1 provides an exemplary overview of the included experiments for each modality. Tables S31-S37 present the full datasets for action (74 experiments, 1378 subjects, 1118 foci), sound (16 experiments, 323 subjects, 275 foci), visual shape (22 experiments, 342 subjects, 256 foci), motion (24 experiments, 450 subjects, 220 foci), color (12 experiments, 207 subjects, 154 foci), olfaction-gustation (17 experiments, 330 subjects, 167 foci), and emotion (47 experiments, 863 subjects, 911 foci).

**Table 1.**
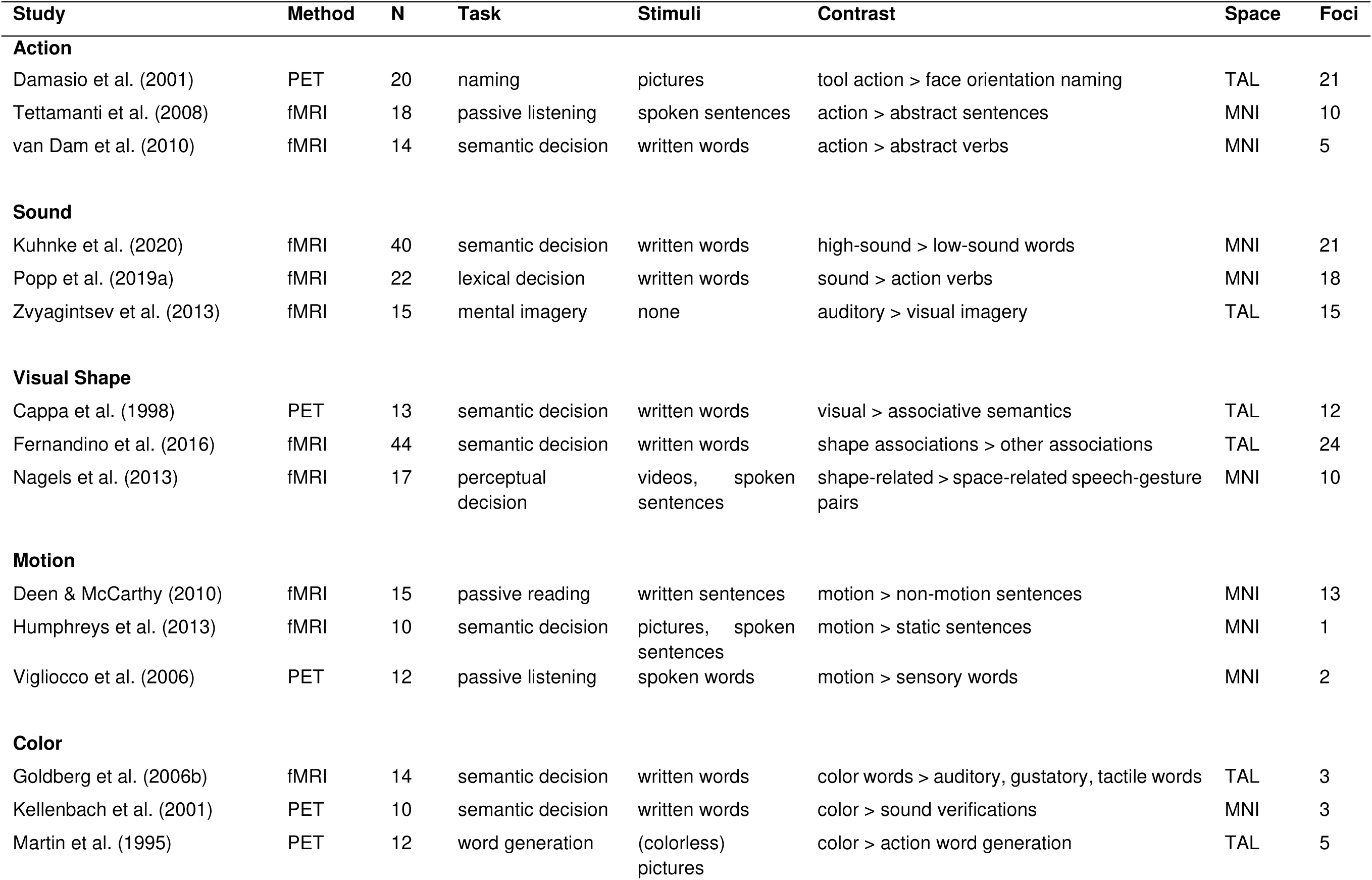
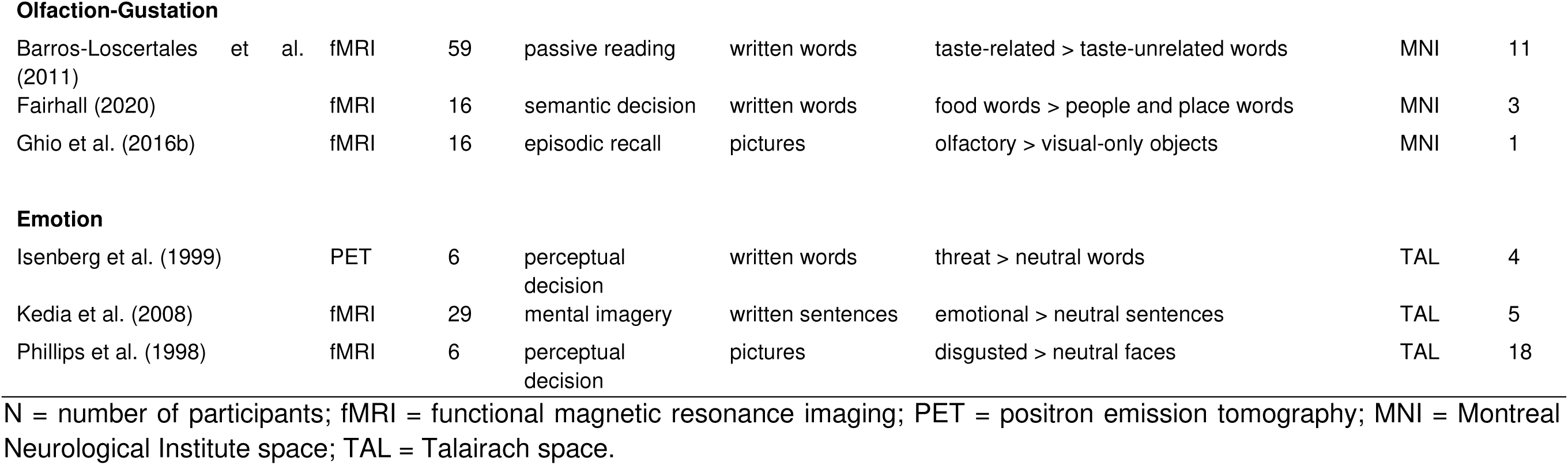
Examples of studies included in the meta-analysis.

| Study | Method | N | Task | Stimuli | Contrast | Space | Foci |
| --- | --- | --- | --- | --- | --- | --- | --- |
| **Action** |  |  |  |  |  |  |  |
| Damasio et al. (2001) | PET | 20 | naming | pictures | tool action > face orientation naming | TAL | 21 |
| Tettamanti et al. (2008) | fMRI | 18 | passive listening | spoken sentences | action > abstract sentences | MNI | 10 |
| van Dam et al. (2010) | fMRI | 14 | semantic decision | written words | action > abstract verbs | MNI | 5 |
| **Sound** |  |  |  |  |  |  |  |
| Kuhnke et al. (2020) | fMRI | 40 | semantic decision | written words | high-sound > low-sound words | MNI | 21 |
| Popp et al. (2019a) | fMRI | 22 | lexical decision | written words | sound > action verbs | MNI | 18 |
| Zvyagintsev et al. (2013) | fMRI | 15 | mental imagery | none | auditory > visual imagery | TAL | 15 |
| **Visual Shape** |  |  |  |  |  |  |  |
| Cappa et al. (1998) | PET | 13 | semantic decision | written words | visual > associative semantics | TAL | 12 |
| Fernandino et al. (2016) | fMRI | 44 | semantic decision | written words | shape associations > other associations | TAL | 24 |
| Nagels et al. (2013) | fMRI | 17 | perceptual decision | videos, spoken sentences | shape-related > space-related speech-gesture pairs | MNI | 10 |
| **Motion** |  |  |  |  |  |  |  |
| Deen & McCarthy (2010) | fMRI | 15 | passive reading | written sentences | motion > non-motion sentences | MNI | 13 |
| Humphreys et al. (2013) | fMRI | 10 | semantic decision | pictures, spoken sentences | motion > static sentences | MNI | 1 |
| Vigliocco et al. (2006) | PET | 12 | passive listening | spoken words | motion > sensory words | MNI | 2 |
| **Color** |  |  |  |  |  |  |  |
| Goldberg et al. (2006b) | fMRI | 14 | semantic decision | written words | color words > auditory, gustatory, tactile words | TAL | 3 |
| Kellenbach et al. (2001) | PET | 10 | semantic decision | written words | color > sound verifications | MNI | 3 |
| Martin et al. (1995) | PET | 12 | word generation | (colorless) pictures | color > action word generation | TAL | 5 |
| **Olfaction-Gustation** |  |  |  |  |  |  |  |
| Barros-Loscertales et al. (2011) | fMRI | 59 | passive reading | written words | taste-related > taste-unrelated words | MNI | 11 |
| Fairhall (2020) | fMRI | 16 | semantic decision | written words | food words > people and place words | MNI | 3 |
| Ghio et al. (2016b) | fMRI | 16 | episodic recall | pictures | olfactory > visual-only objects | MNI | 1 |
| **Emotion** |  |  |  |  |  |  |  |
| Isenberg et al. (1999) | PET | 6 | perceptual decision | written words | threat > neutral words | TAL | 4 |
| Kedia et al. (2008) | fMRI | 29 | mental imagery | written sentences | emotional > neutral sentences | TAL | 5 |
| Phillips et al. (1998) | fMRI | 6 | perceptual decision | pictures | disgusted > neutral faces | TAL | 18 |

N = number of participants; fMRI = functional magnetic resonance imaging; PET = positron emission tomography; MNI = Montreal Neurological Institute space; TAL = Talairach space.

### Activation Likelihood Estimation (ALE)

We performed a coordinate-based activation likelihood estimation (ALE) meta-analysis using *GingerALE* version 3.0 (https://brainmap.org/ale). ALE treats reported activation coordinates as centers of 3D Gaussian probability distributions, whose width depends on an empirical model of between-template and between-participant variance (Eickhoff et al., 2012, 2009; Turkeltaub et al., 2012). In practice, larger samples receive tighter distributions. For each experiment, these probability distributions are combined into a “modelled activation” (MA) map. Taking the voxel-wise union of all MA maps yields the ALE map, which quantifies the convergence of activations across experiments (Eickhoff et al., 2012). The ALE map is compared to a null distribution reflecting random spatial association between experiments. This results in a random-effects inference, allowing for generalization to the entire population of studies (Eickhoff et al., 2009).

### Main ALE analyses for each conceptual modality

Our primary goal was to identify brain regions that are consistently engaged across neuroimaging studies for conceptual processing related to one of the 7 perceptual-motor modalities (action, sound, visual shape, motion, color, olfaction-gustation, or emotion). Therefore, we first ran an ALE meta-analysis for each modality separately. To minimize within-sample effects, we used the conservative Turkeltaub ALE method (which eliminates effects of number and proximity of reported foci for each study), and pooled together multiple contrasts from the same study and sample (Turkeltaub et al., 2012). All meta-analyses were performed in MNI space; TAL coordinates were converted to MNI space using the *tal2icbm* algorithm in *GingerALE* (Lancaster et al., 2007). ALE maps were thresholded at a voxel-wise p < 0.001 and a cluster-wise p < 0.05 FWE-corrected for multiple comparisons using Monte Carlo simulation (10,000 permutations; as proposed by Müller et al., 2018).

### Overlap between conceptual and perceptual-motor regions

Second, we aimed to test the key prediction of grounded cognition theories that conceptual processing related to a certain perceptual-motor modality involves brain regions also engaged during real perception or action in that modality. To this end, we first searched the *BrainMap* database (using the *Sleuth* software; http://www.brainmap.org/sleuth/) for neuroimaging studies on actual perception or action in each modality. We queried the following behavioral domains restricted to activation-based experiments on healthy human participants: for action execution = Action.Execution (451 experiments, 5037 subjects, 12241 foci); auditory perception = Perception.Audition (152 experiments, 2068 subjects, 3670 foci); visual shape perception = Perception.Vision.Shape (147 experiments, 1869 subjects, 3074 foci); visual motion perception = Perception.Vision.Motion (97 experiments, 1019 subjects, 2703 foci); color perception = Perception.Vision.Color (40 experiments, 665 subjects, 835 foci); olfaction-gustation = Perception.Olfaction or Perception.Gustation (98 experiments, 1420 subjects, 1980 foci); and emotion = Emotion (597 experiments, 9271 subjects, 14718 foci). For each modality, we then performed an ALE meta-analysis using the same ALE methods and thresholding as our conceptual processing analyses. Finally, we computed the overlap between the meta-analytic maps for conceptual processing related to a certain modality (e.g. sound) and real perception/action in that modality (e.g. auditory perception) via minimum-statistic conjunction (testing the conjunction null; Nichols et al., 2005).

### Contrast analyses for modality specificity

Third, we investigated the modality specificity of the brain regions engaged for conceptual processing related to a certain modality. To this end, we performed direct contrasts between the ALE maps for one modality vs. every other modality (e.g. for action: action > sound; action > visual shape; action > motion; action > color; action > olfaction-gustation; and action > emotion). *GingerALE* employs the following procedure for contrast analysis (Eickhoff et al., 2011; Langner and Eickhoff, 2013): First, the voxel-wise difference between ALE maps is computed. To account for sample size differences, the two datasets are pooled and randomly divided into two new datasets of equal sizes. An ALE image is then created for each new dataset and subtracted from the other. We repeated this procedure 10,000 times to create a null distribution of ALE-value differences, which was used to threshold the observed difference map at a posterior probability of P > 0.95 for a true difference (Cieslik et al., 2016; Hardwick et al., 2018). To quantify modality specificity, we performed (minimum-statistic) conjunctions between all individual contrasts, and tested how many contrasts were significant in each voxel. All contrasts were inclusively masked by regions that were significantly engaged only for the respective modality (e.g. action) and no other modality (cf. Hardwick et al., 2018; Rottschy et al., 2012). Hence, we exclusively called regions “modality-specific” if they showed significant convergence for only a single modality *and* higher activation likelihood for that modality than for the other modalities. Brain regions with the strongest modality specificity were significant for the conjunction across all 6 contrasts (e.g. for action: [action > sound] ∩ [action > visual shape] ∩ [action > motion] ∩ [action > color] ∩ [action > olfaction-gustation] ∩ [action > emotion]).

### Conjunction analyses for multimodal convergence zones

Fourth, we aimed to identify “multimodal convergence zones”—brain regions engaged in conceptual processing related to multiple different perceptual-motor modalities. To this end, we computed the overlap between the ALE maps for all conceptual modalities. Again, we performed minimum-statistic conjunctions, testing the conjunction null (Nichols et al., 2005). That is, a voxel was only considered significant in the conjunction if it was significant for each involved modality. We then tested *how many* modalities overlap in which areas to determine “bimodal” (2 modalities), “trimodal” (3 modalities) areas, and so on. Finally, we analyzed *which* modalities overlap in the different multimodal areas.

## Results

### Action

Across neuroimaging studies, action-related conceptual processing consistently engaged the left inferior frontal gyrus (IFG) and premotor cortex (PMC), anterior supramarginal gyrus (aSMG) extending into primary somatosensory cortex (S1) and intraparietal sulcus (IPS), the lateral temporal-occipital junction (LTO) including parts of posterior middle and inferior temporal gyri (pMTG/ITG) and superior temporal sulcus (pSTS), as well as the bilateral (pre-)supplementary motor area (SMA) (Figure 2A; Table S2).

**Figure 2.**
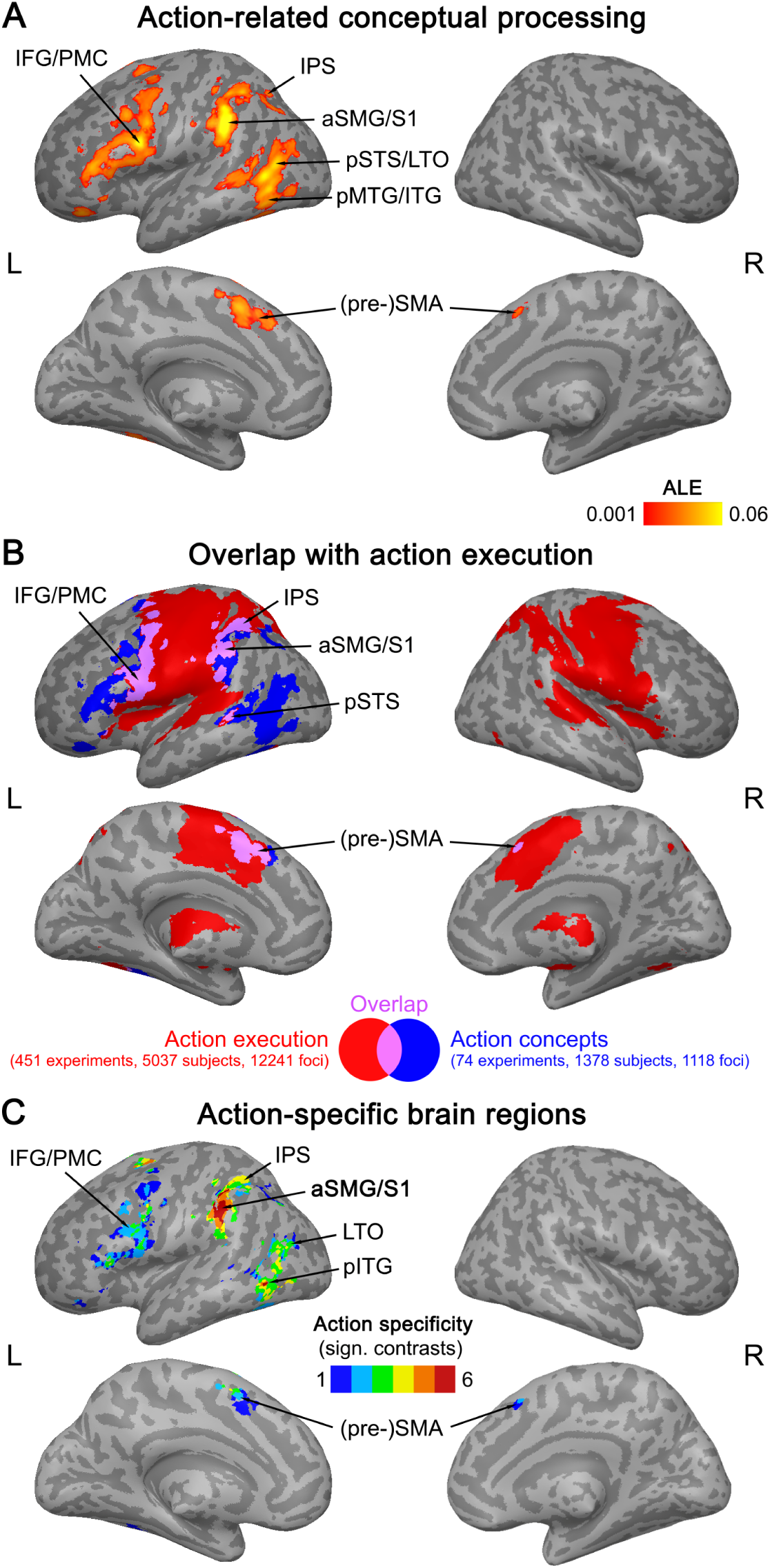
(A) ALE meta-analytic map for action-related conceptual processing (thresholded at a voxel-wise p < 0.001, cluster-wise p < 0.05 FWE-corrected). (B) Overlap (purple) between meta-analytic maps for action-related conceptual processing (blue) and real action execution (red). (C) Regions showing consistent engagement for conceptual processing related to action and no other modality, and higher activation likelihood for action than the other modalities (number of significant contrasts is displayed).

Real action execution robustly activated the bilateral primary motor cortex (M1), PMC, (pre-)SMA, aSMG extending into IPS and S1, pSTG/STS, thalamus, and cerebellum (Figure 2B red; Table S3). Action-related conceptual processing overlapped with real action execution in left IFG/PMC, aSMG/S1 and IPS, pSTS, and in bilateral (pre-)SMA (Figure 2B purple; Table S4).

Left aSMG/S1 and pITG were action-specific (Figure 2C; Table S5): These regions were significantly engaged only for action and no other modality, and contrast analyses revealed significantly higher activation likelihood for action than every other modality. Weaker evidence for action specificity was found in left IFG/PMC, IPS, LTO, and bilateral (pre-)SMA: These areas showed significant engagement selectively for action, and a higher activation likelihood for action than for several, but not all, other modalities.

A supplementary analysis that excluded low-level contrasts (e.g. action words > fixation) yielded qualitatively similar results (Figure S1), confirming their robustness to baseline differences. In particular, left aSMG/S1, IPS, IFG/PMC, LTO/pSTS, and pMTG/ITG (but not pre-SMA) were robustly engaged in action-related conceptual processing. Left aSMG/S1 and pITG showed high action specificity.

### Sound

Sound-related conceptual processing consistently activated the bilateral pSTS (extending into pMTG in the left hemisphere), as well as left pSMG and dorsomedial prefrontal cortex (dmPFC) (Figure 3A; Table S6).

**Figure 3.**
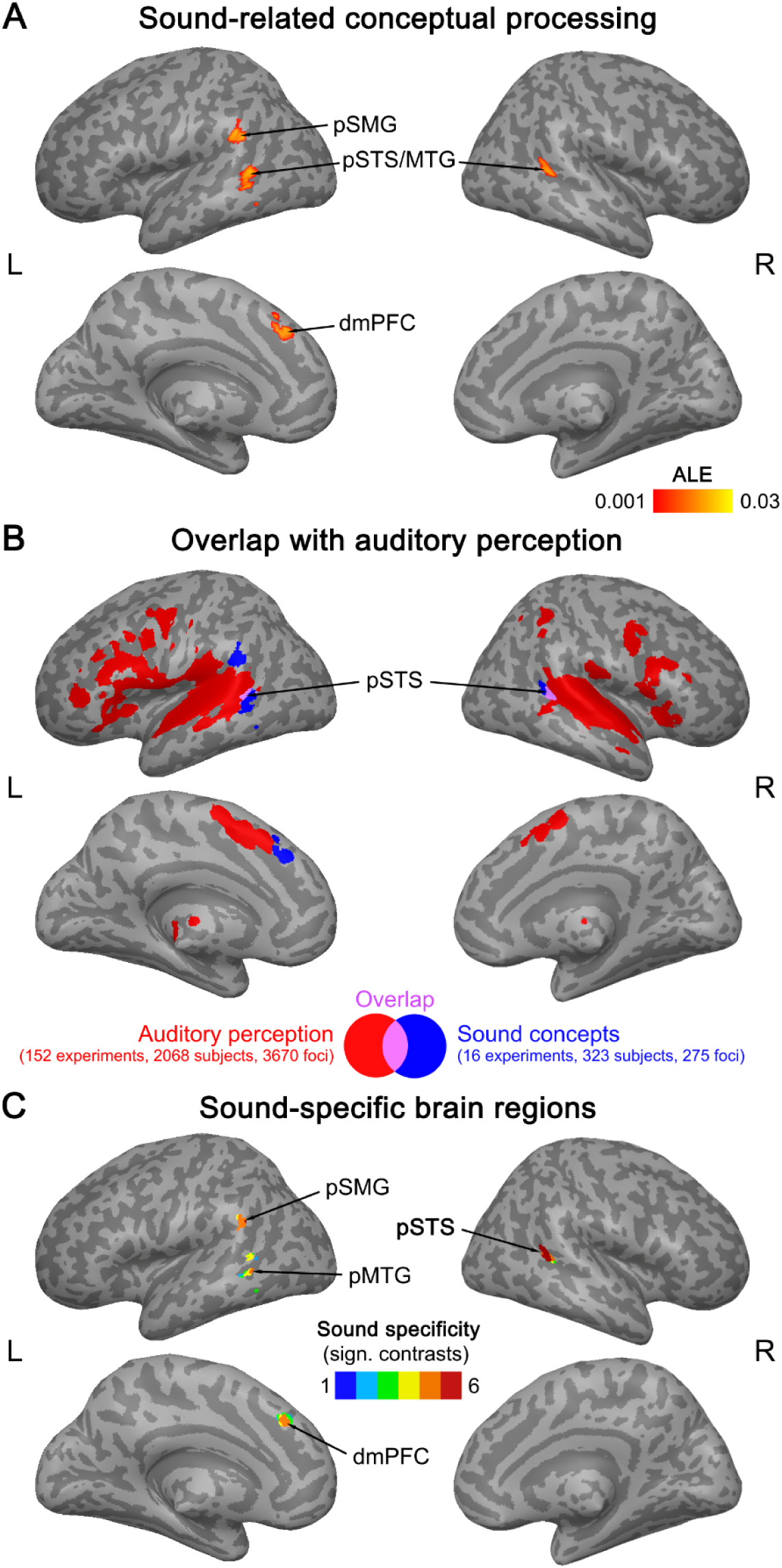
(A) ALE meta-analytic map for sound-related conceptual processing (thresholded at a voxel-wise p < 0.001, cluster-wise p < 0.05 FWE-corrected). (B) Overlap (purple) between meta-analytic maps for sound-related conceptual processing (blue) and real auditory perception (red). (C) Regions showing consistent engagement for conceptual processing related to sound and no other modality, and higher activation likelihood for sound than the other modalities (number of significant contrasts is displayed).

Real auditory perception robustly engaged the bilateral early auditory cortex and surrounding STG/MTG, IFG (extending into insula), middle frontal gyrus (MFG) and precentral sulcus (PreCS), dmPFC, thalamus, and right IPS (Figure 3B red; Table S7). Sound-related conceptual processing overlapped with real auditory perception in bilateral pSTS (Figure 2B purple; Table S8). Left dmPFC and pSMG areas engaged for sound-related conceptual processing were adjacent to, but non-overlapping with regions engaged for auditory perception.

The right pSTS was sound-specific (Figure 3C; Table S9), showing consistent engagement exclusively for sound (and no other modality) and higher activation likelihood for sound than every other modality. Left pMTG, pSMG, and dmPFC showed weaker evidence for sound specificity, with significant convergence selectively for sound and higher activation likelihood for sound than multiple, but not all, other modalities.

A supplementary analysis without low-level contrasts (e.g. sound-related words > fixation) yielded very similar results (Figure S2). Indeed, this analysis provided even stronger evidence for sound specificity in right pSTS, left pMTG, and pSMG. In contrast, left dmPFC was not robustly engaged.

### Visual shape

Conceptual processing related to visual shape consistently activated the left precentral sulcus (PreCS), lateral occipital cortex (LOC; area hOc4la), and anterior fusiform gyrus (FG) (Figure 4A; Table S10).

**Figure 4.**
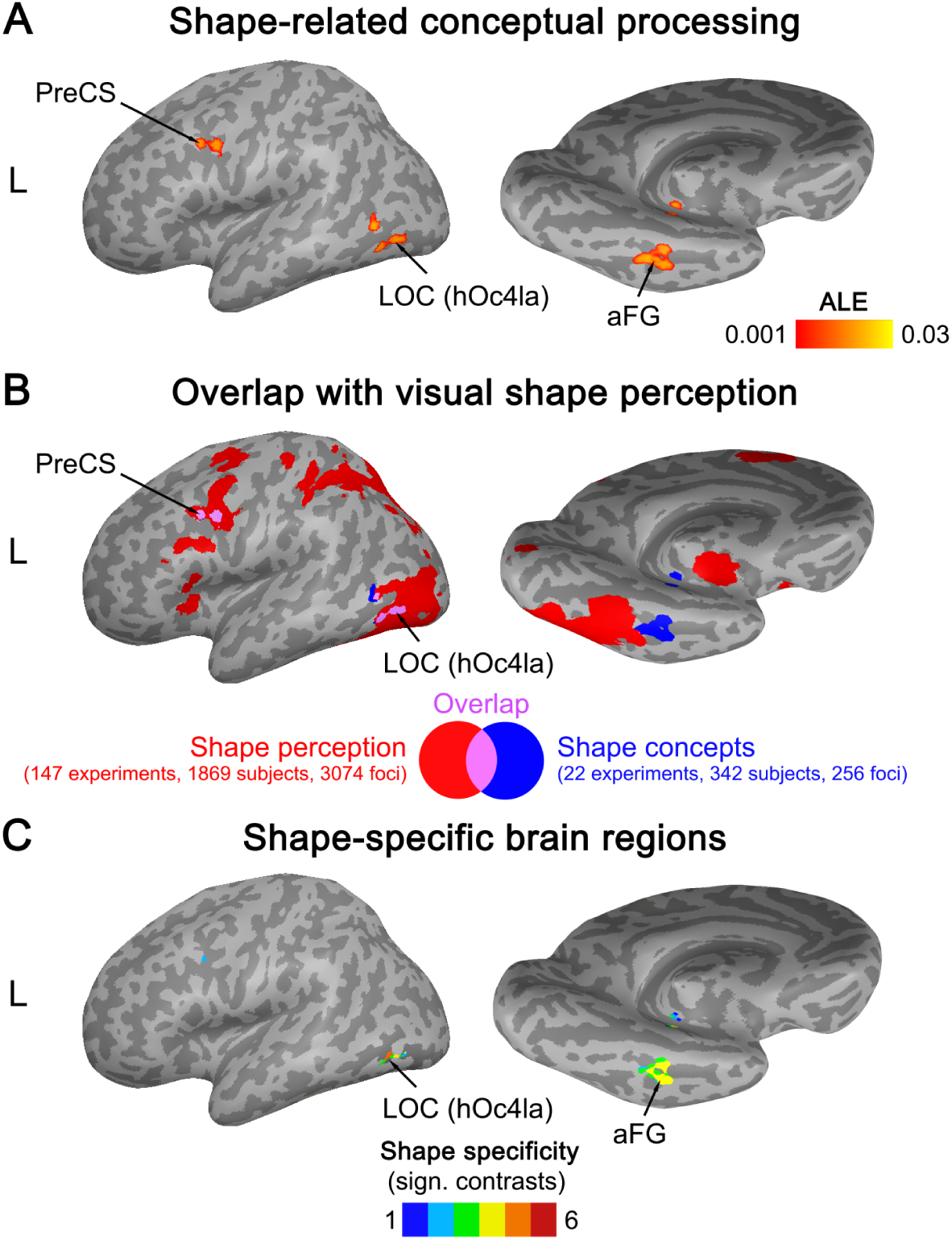
(A) ALE meta-analytic map for conceptual processing related to visual shape (thresholded at a voxel-wise p < 0.001, cluster-wise p < 0.05 FWE-corrected). (B) Overlap (purple) between meta-analytic maps for shape-related conceptual processing (blue) and real visual shape perception (red). (C) Regions showing consistent engagement for conceptual processing related to visual shape and no other modality, and higher activation likelihood for shape than the other modalities (number of significant contrasts is displayed).

Real visual shape perception robustly engaged the bilateral early visual cortex (V1/V2/V3/V4), LOC (hOc4la/p), IPS/SPL, FG, PreCS extending into PMC, IFG, and insula (Figure 4B red; Table S11). Shape-related conceptual processing overlapped with real visual shape perception in left PreCS and LOC (hOc4la), whereas the FG clusters were directly adjacent but non-overlapping (Figure 4B purple; Table S12).

Left LOC (hOc4la) and aFG showed evidence for shape specificity, with consistent engagement selectively for shape (and no other modality) and higher activation likelihood for shape than most other modalities (Figure 4C; Table S13).

A supplementary analysis that excluded low-level contrasts (e.g. shape-related words > fixation) provided even stronger evidence that left LOC (hOc4la) was robustly and selectively engaged in shape-related conceptual processing (Figure S3). However, left aFG did not show significant convergence in this analysis.

### Motion

Motion-related conceptual processing consistently recruited the left pSTS extending into pMTG and aSMG, as well as left IFG and dorsal FG (Figure 5A; Table S14).

**Figure 5.**
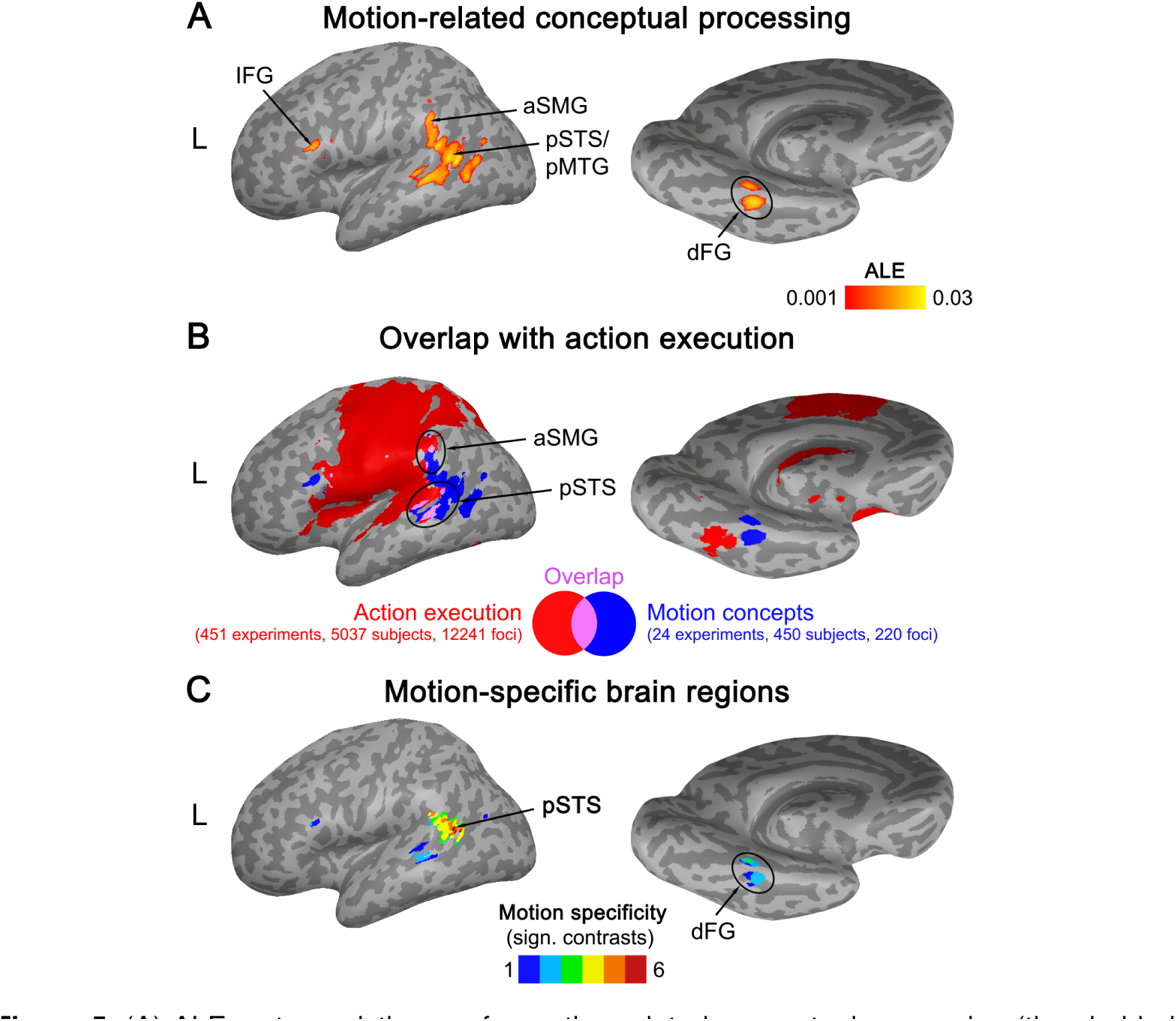
(A) ALE meta-analytic map for motion-related conceptual processing (thresholded at a voxel-wise p < 0.001, cluster-wise p < 0.05 FWE-corrected). (B) Overlap (purple) between meta-analytic maps for motion-related conceptual processing (blue) and real action execution (red). (C) Regions showing consistent engagement for conceptual processing related to motion and no other modality, and higher activation likelihood for motion than the other modalities (number of significant contrasts is displayed).

Motion-related conceptual processing did not overlap with real motion perception, which robustly recruited the bilateral early visual cortex (V1/V2), LOC, area V5/MT, SPL/IPS, PMC, and SMA (Figure S8; Table S15). However, motion-related conceptual processing overlapped with real action execution in left pSTS and aSMG (Figure 5B; Table S16).

Left pSTS was motion-specific (Figure 5C; Table S17): A cluster in left pSTS was consistently engaged only for motion (and no other modality) and showed significantly higher activation likelihood for motion than all other modalities. Left dFG showed weaker evidence for motion specificity, with significant convergence only for motion and higher activation likelihood for motion than some, but not all, other modalities.

A supplementary meta-analysis excluding low-level contrasts (e.g. motion words > fixation) revealed highly similar results (Figure S4), confirming that left pSTS and dFG are robustly engaged for motion-related conceptual processing, where left pSTS is motion-specific. In contrast to the full analysis, left IFG was not consistently engaged.

### Color

Color-related conceptual processing robustly engaged the left IPS and ventral FG (vFG) across neuroimaging experiments (Figure 6A; Table S18).

**Figure 6.**
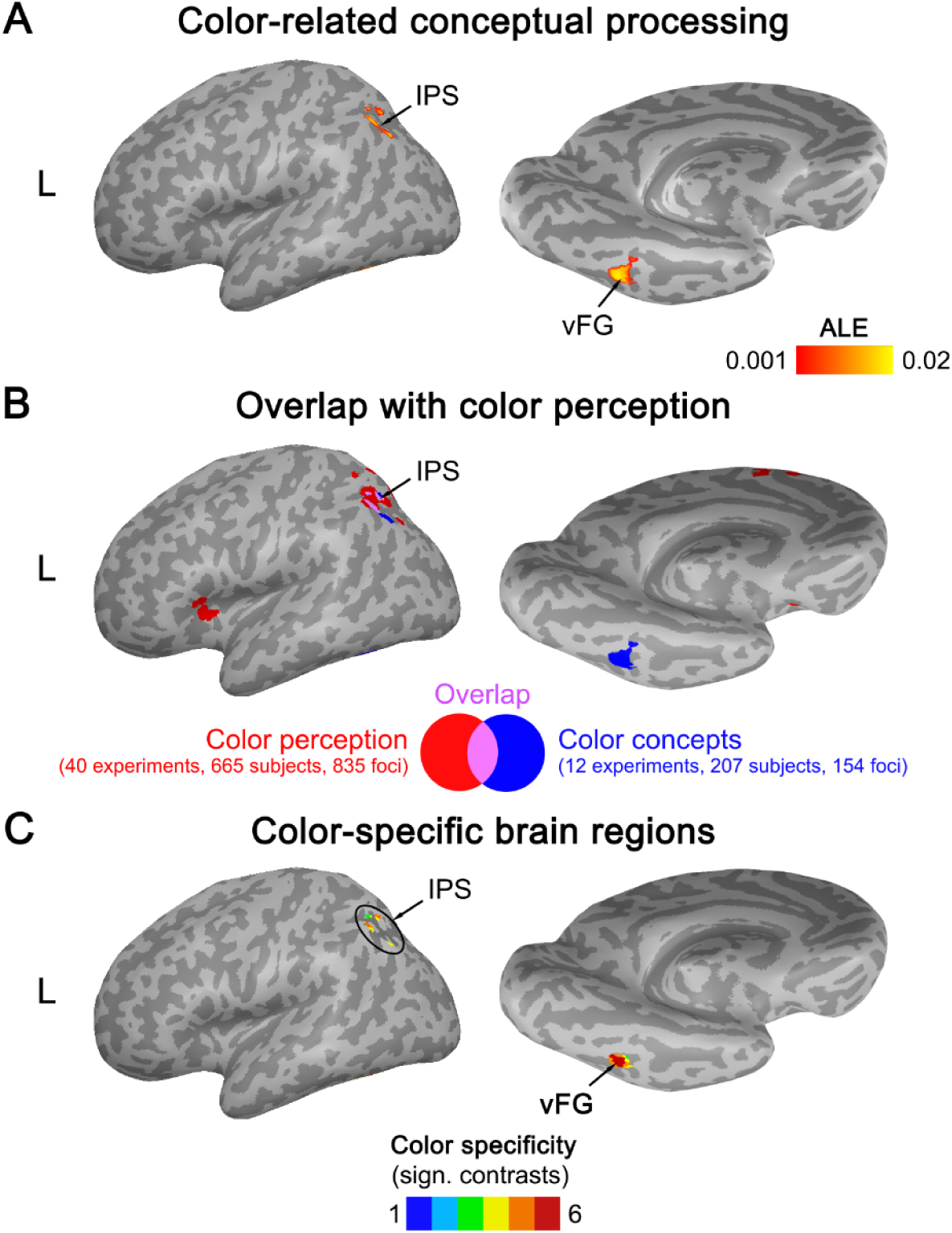
(A) ALE meta-analytic map for color-related conceptual processing (thresholded at a voxel-wise p < 0.001, cluster-wise p < 0.05 FWE-corrected). (B) Overlap (purple) between meta-analytic maps for color-related conceptual processing (blue) and real color perception (red). (C) Regions showing consistent engagement for conceptual processing related to color and no other modality, and higher activation likelihood for color than the other modalities (number of significant contrasts is displayed).

Real color perception consistently activated bilateral IPS (extending into SPL), insula, pre-SMA, left precuneus, and right LOC (Figure 6B red; Table S19). Color-related conceptual processing and real color perception overlapped in left IPS (Figure 5B; Table S20). The left vFG region engaged for color concepts was not consistently engaged for color perception.

Left vFG was color-specific (Figure 6C; Table S21), with significant convergence selectively for color (and no other modality) and higher activation likelihood for color than all other modalities. Left IPS showed weaker evidence for color specificity: This area was consistently engaged only for color, and more consistently activated for color than several, albeit not all, other modalities.

A supplementary analysis excluding low-level contrasts (e.g. color words > fixation) provided qualitatively identical results (Figure S3), confirming that color-related conceptual processing robustly engages left IPS and vFG, where vFG is color-specific.

### Olfaction-Gustation

Conceptual processing related to olfaction-gustation was associated with consistent activation in left orbitofrontal cortex (OFC) (Figure 7A; Table S22).

**Figure 7.**
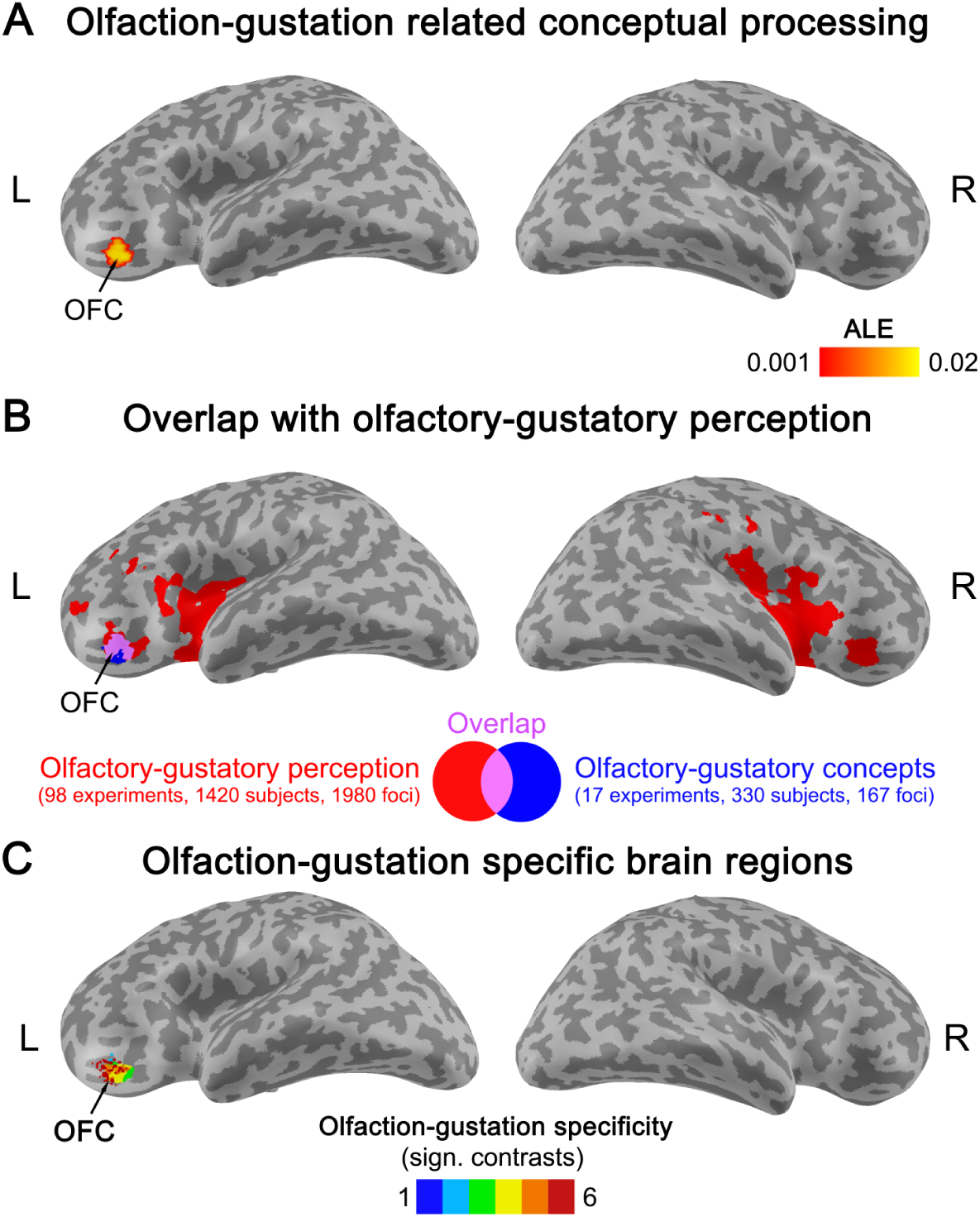
(A) ALE meta-analytic map for conceptual processing related to olfaction-gustation (thresholded at a voxel-wise p < 0.001, cluster-wise p < 0.05 FWE-corrected). (B) Overlap (purple) between meta-analytic maps for olfaction-gustation related conceptual processing (blue) and real olfactory-gustatory perception (red). (C) Regions showing consistent engagement for conceptual processing related to olfaction-gustation and no other modality, and higher activation likelihood for olfaction-gustation than the other modalities (number of significant contrasts is displayed).

Real olfactory-gustatory perception robustly activated the bilateral OFC, insula, IFG, anterior cingulate, thalamus, basal forebrain, as well as the amygdala, hippocampus, entorhinal cortex, and subiculum (Figure 7B red; Table S23). Conceptual processing related to olfaction-gustation and real olfactory-gustatory perception overlapped in the left OFC (Figure 7B purple; Table S24).

Left OFC was specific to olfactory-gustatory conceptual processing (Figure 7C; Table S25): This area showed significant convergence only for olfaction-gustation (and no other modality) and higher activation likelihood for olfaction-gustation than every other modality.

A supplementary analysis that excluded low-level contrasts (e.g. food words > rest) yielded highly similar results (Figure S6), confirming that left OFC is robustly and selectively engaged in conceptual processing related to olfaction-gustation.

### Emotion

Emotion-related conceptual processing consistently activated the bilateral amygdala and medial prefrontal cortex (mPFC), as well as the left angular gyrus (AG) and temporal pole (TP) (Figure 8A; Table S26).

**Figure 8.**
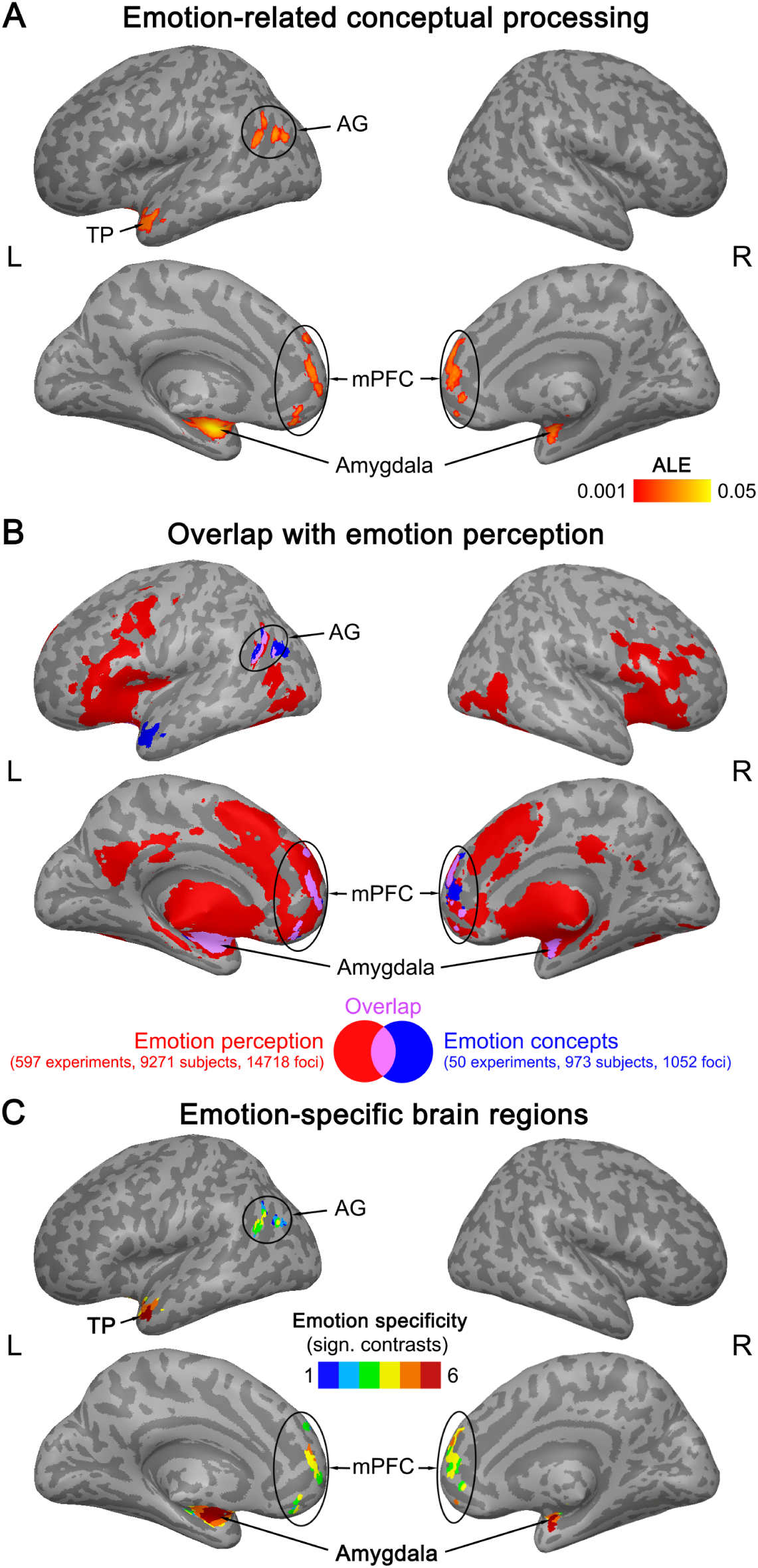
(A) ALE meta-analytic map for emotion-related conceptual processing (thresholded at a voxel-wise p < 0.001, cluster-wise p < 0.05 FWE-corrected). (B) Overlap (purple) between meta-analytic maps for emotion-related conceptual processing (blue) and real emotion perception (red). (C) Regions showing consistent engagement for conceptual processing related to emotion and no other modality, and higher activation likelihood for emotion than the other modalities (number of significant contrasts is displayed).

Real emotion perception robustly recruited the bilateral amygdala and hippocampus, thalamus, basal forebrain, IFG and insula, PreCS, FG, LOC, cingulate and mPFC (Figure 8B red; Table S27). Emotion-related conceptual processing overlapped with real emotion perception in the bilateral amygdala, mPFC, and left AG (Figure 8B purple; Table S28). Left TP was not consistently engaged for real emotion perception.

Bilateral amygdala and left TP were emotion-specific (Figure 8C; Table S29), showing significant engagement selectively for emotion (and no other modality) and higher activation likelihood for emotion than all other modalities. Weaker evidence for emotion specificity was found in left AG and bilateral mPFC, which were consistently engaged exclusively for emotion and more consistently engaged for emotion than several, but not all, other modalities.

A supplementary analysis without low-level contrasts (e.g. emotionally-connotated words > fixation) provided similar results (Figure S7), confirming the consistent engagement of left AG, bilateral amygdala and mPFC, as well as the emotion specificity of bilateral amygdala. However, left TP did not show significant convergence in this analysis.

### Multimodal convergence zones

To identify “multimodal” brain regions consistently engaged for conceptual processing related to multiple modalities, we performed conjunction analyses between all possible combinations of modalities. We found that several brain regions were recruited for two (“bimodal”), or three (“trimodal”) modalities (Figure 9A). No region was engaged for more than three modalities.

**Figure 9.**
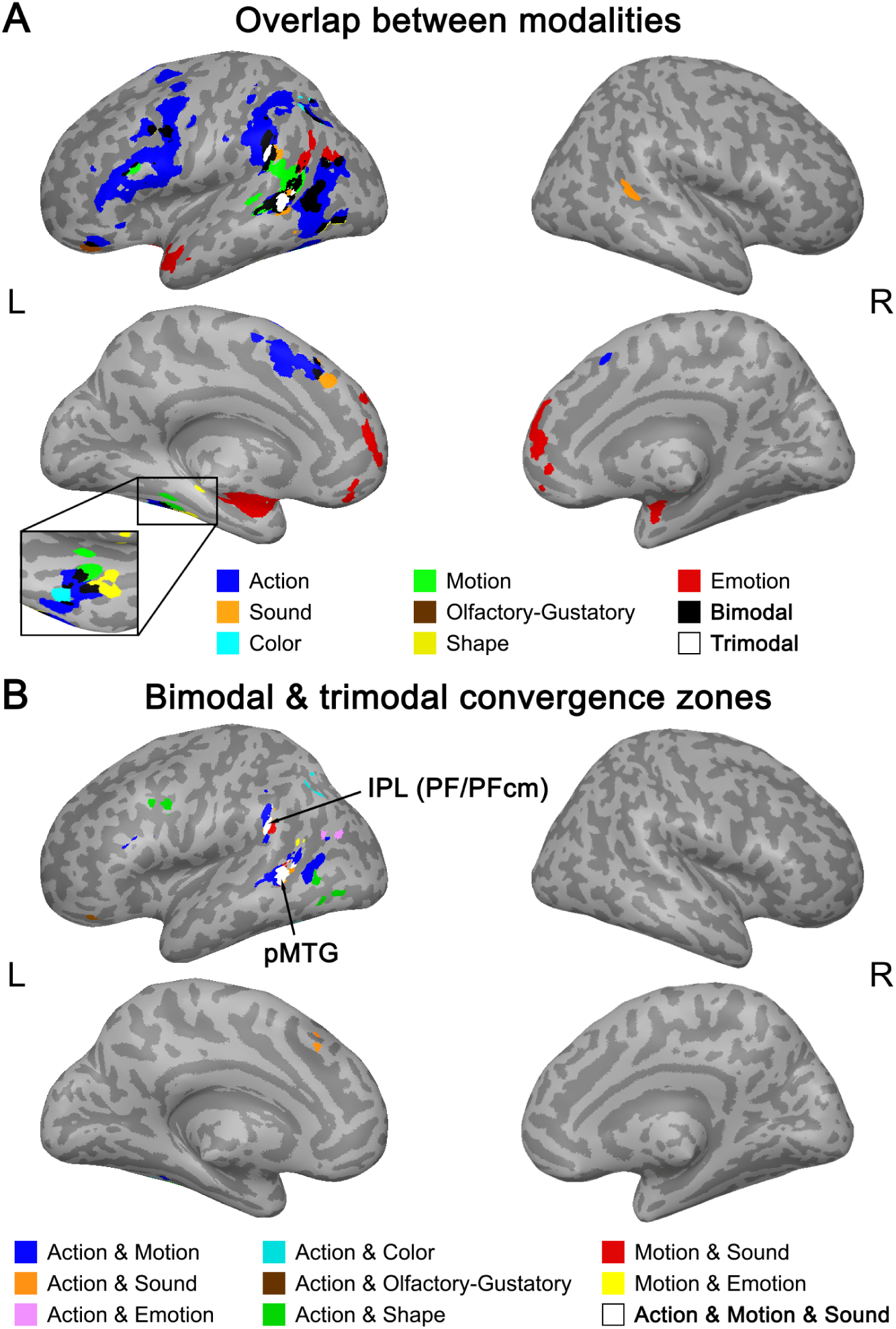
(A) Overlap between meta-analytic maps for conceptual processing related to the different perceptual-motor modalities. All maps were thresholded at a voxel-wise p < 0.001 and a cluster-wise p < 0.05 FWE-corrected. (B) Multimodal convergence zones that were consistently engaged for multiple modalities. Trimodal regions in left IPL and pMTG were surrounded by bimodal areas.

In particular, the left posterior middle temporal gyrus (pMTG) and inferior parietal lobe (IPL; area PF/PFcm) contained “trimodal” convergence zones engaged for action, motion, and sound (Figure 9B; Table S30). These trimodal areas were surrounded by “bimodal” regions for action and motion (blue), motion and sound (red), and action and sound (orange).

Other bimodal regions were distributed throughout the left hemisphere—for action and shape (green; LOC, PreCS, aFG), action and color (cyan; IPS, vFG), action and emotion (purple; pAG), action and olfaction-gustation (brown; OFC), and motion and emotion (yellow, TPJ).

## Discussion

Here, we investigated the neural basis of conceptual processing in the healthy human brain. Specifically, we (1) tested the grounded cognition hypothesis that conceptual processing consistently recruits modality-specific perceptual-motor regions, and (2) investigated whether and which modalities overlap in multimodal convergence zones. To this end, we performed an activation likelihood estimation (ALE) meta-analysis across 212 functional neuroimaging experiments on conceptual processing related to 7 perceptual-motor modalities: action, sound, visual shape, motion, color, olfaction-gustation, and emotion.

We found that conceptual processing consistently engages brain regions that are also activated during real perceptual-motor experience of the same modalities. These perceptual-motor areas exhibit a strong modality specificity, that is, a higher activation likelihood for the respective modality than for the other modalities. These results support grounded cognition theories: Conceptual processing robustly recruits modality-specific perceptual-motor regions.

In addition to modality-specific perceptual-motor areas, we identified multimodal convergence zones that are robustly engaged for multiple modalities. In particular, the left inferior parietal lobe (IPL) and posterior middle temporal gyrus (pMTG) are activated for three modalities: action, motion, and sound. These “trimodal” areas are surrounded by “bimodal” areas engaged for two modalities. Taken together, our findings support “hybrid theories” of conceptual processing which propose an involvement of both modality-specific and cross-modal brain regions.

### Modality-specific perceptual-motor regions

#### Action

Action-related conceptual processing overlapped with real action execution in high-level somatomotor regions, including left aSMG/S1, IPS, PMC, and (pre-)SMA. These regions are recruited during real tool use (Goldenberg, 2009; Johnson-Frey, 2004; Lewis, 2006), as well as during action observation, pantomime and imagery (Hardwick et al., 2018; Papitto et al., 2019). Left aSMG/S1 showed the strongest evidence for action specificity, that is, a higher activation likelihood for conceptual processing related to action than all other modalities. Left aSMG is implicated in the visual-motor control of object-directed actions (Haaland et al., 2000; Turella and Lingnau, 2014). In particular, ideomotor apraxia—a deficit in producing skilled object-directed movements (Culham and Valyear, 2006)—is specifically associated with damage in and near left aSMG (Buxbaum et al., 2005a, 2005b; Haaland et al., 2000). These observations suggest that left aSMG stores high-level representations of object-use motor skills (Culham and Valyear, 2006; Johnson-Frey, 2004; van Elk et al., 2014).

Notably, action comprises both motor and somatosensory processes, which are notoriously difficult to disentangle in neuroimaging experiments. Crucially, both motor and somatosensory areas are involved in object-directed actions (Hardwick et al., 2018; van Elk et al., 2014) as well as action-related conceptual processing (Desai et al., 2010; Fernandino et al., 2016a). Nonetheless, motor and somatosensory areas may play distinct roles in action knowledge processing, representing the movement-vs. touch-related components of object-directed actions, respectively. Future studies should aim to disentangle the motor and somatosensory components of action concepts. Overall, our results indicate that action-related conceptual processing involves high-level somatomotor representations that are also engaged during real object-directed actions.

#### Sound

Sound-related conceptual processing and real auditory perception overlapped in the bilateral pSTS. The pSTS is implicated in high-level auditory processes, such as environmental sound recognition (Lewis et al., 2004), sound recall (Wheeler et al., 2000), music imagery (Zatorre et al., 1996), and voice perception (Belin et al., 2000; Specht and Reul, 2003). Right pSTS showed the strongest evidence for sound specificity, with a higher activation likelihood for sound than all other modalities. This finding converges with a right-hemispheric dominance for non-linguistic environmental sounds, such as music (Halpern et al., 2004), human voices (Belin et al., 2000), and animal vocalizations (Lewis et al., 2009). In contrast, speech perception is strongly left-lateralized (Friederici, 2012, 2002). This suggests that auditory features of concepts comprise high-level environmental sound representations (e.g. barking), and not internal verbalizations (e.g. “dog”). Overall, these results indicate that sound-related conceptual processing involves brain regions that are also engaged in high-level auditory processing.

#### Visual shape

Both shape-related conceptual processing and real visual shape perception consistently activated the left LOC (area hOc4la). Left LOC (hOc4la) was also shape-specific, being selectively and more consistently recruited for shape than for the other modalities. The LOC is part of extrastriate visual cortex, situated downstream from early visual cortex (V1-V3) and implicated in visual object perception (Malikovic et al., 2016; Weiner and Grill-Spector, 2011). The LOC responds to differences in the perceived 3D shape of objects (Kourtzi et al., 2003; Murray et al., 2002), as well as the 2D shapes of familiar and illusory objects (Stanley and Rubin, 2005). Area hOc4la of the LOC can be structurally and functionally distinguished from its neighbors hOc4lp and V5/MT: While both hOc4la and hOc4lp are associated with visual shape perception, only hOc4la contains object-selective representations (Sayres and Grill-Spector, 2008). In contrast to V5/MT, hOc4la is not motion-sensitive, but strongly responds to images of intact vs. scrambled objects (Larsson and Heeger, 2006). Taken together, these results suggest that shape-related conceptual processing recruits high-level representations of object shape.

Left aFG was also consistently engaged in shape-related conceptual processing. This converges with several neuroimaging studies indicating that the shape properties of animals and tools are represented in the FG in both perceptual and conceptual tasks (for a review, see Martin, 2007). However, in this meta-analysis, the FG area engaged in conceptual processing did not overlap with, but lay directly anterior to the FG area engaged in real visual shape perception. This may reflect abstraction from low-level visual shape information (Thompson-Schill, 2003; see discussion of “anterior shift” below).

#### Motion

Motion-related conceptual processing did not overlap with real visual motion perception, but with action execution in left pSTS and aSMG. At first glance, this result might suggest that motion concepts are more related to movement execution than perception. This view is supported by the fact that left aSMG is implicated in tool-use motor skills (Johnson-Frey, 2004; van Elk et al., 2014; see section on Action).

However, left pSTS is rather implicated in movement perception than in movement execution (Pitcher and Ungerleider, 2021). Indeed, a cluster within left pSTS showed strong evidence for motion specificity, that is, a higher activation likelihood for motion than for all other modalities, including action. Moreover, this motion-specific cluster did not overlap with real action execution. Left pSTS is known to be specialized for biological (i.e. human and animal) motion perception (for a review, see Saygin, 2012). The pSTS receives input from the classical motion-sensitive area V5/MT which does not distinguish different types of movement (Boussaoud et al., 1990; Ungerleider and Desimone, 1986). Together with these previous findings, our results suggest that motion-related conceptual processing involves high-level representations of (biological or action-related) motion.

#### Color

Color-related conceptual processing consistently engaged the left IPS and vFG. Remarkably, overlap with real color perception was found in left IPS, but not in vFG. Left IPS is not typically implicated in color perception. However, left IPS is implicated in storing individual object features in visual working memory, including color features (Galeano Weber et al., 2016; Xu, 2007). Thus, the left IPS recruitment during color-related conceptual processing may reflect the online working-memory storage of color representations, rather than color representation *per se*. In line with this view, left IPS showed only weak evidence for color specificity and virtually the same region was also engaged in action-related conceptual processing.

In contrast, left vFG showed strong evidence for color specificity. Left vFG is specifically implicated in active color perception (e.g. color discrimination tasks; Beauchamp, 1999; Simmons et al., 2007), and receives input from area V4 implicated in passive color perception (Conway, 2009). However, left vFG was not consistently activated during real color perception in our meta-analysis. As a potential explanation, the *BrainMap* dataset possibly included too few active color perception tasks to engage left vFG. Notably, several individual studies demonstrated direct activation overlap between color-related conceptual processing and active color perception in left vFG (Hsu et al., 2012, 2011; Simmons et al., 2007). Therefore, we conclude that color-related conceptual processing involves high-level color perception areas.

It should be noted that our dataset for color-related conceptual processing comprised only 12 independent experiments. A minimum of ∼17 experiments is generally recommended for ALE meta-analyses to ensure that results are not driven by single studies (Müller et al., 2018). Therefore, we analyzed the contribution of experiments to each cluster, and found that 4 experiments each contributed to the IPS and vFG clusters, indicating that these clusters were not completely idiosyncratic. Nonetheless, the results for color-related conceptual processing should be taken with some caution.

#### Olfaction-Gustation

Conceptual processing related to olfaction-gustation and real olfactory-gustatory perception overlapped in left OFC. Left OFC was also highly specific to olfactory-gustatory processing, with significant engagement selectively for olfaction-gustation and a higher activation likelihood for olfaction-gustation than for all other modalities.

The OFC is implicated in both olfaction and gustation, receiving input from both primary olfactory and gustatory cortices (de Araujo et al., 2003; Small and Prescott, 2005). Specifically, the OFC is involved in the recognition of odors and flavors, as well as the computation of their reward value, which translates to degrees of (un-)pleasantness on the behavioral level (Kemmerer, 2014; Small et al., 2007). Therefore, our results indicate that conceptual processing related to olfaction-gustation involves high-level components of the olfactory-gustatory system.

#### Emotion

Emotion-related conceptual processing overlapped with real emotion perception in the left TPJ, bilateral mPFC and amygdala. The TPJ is involved in social-emotional processes, particularly in theory-of-mind or mentalizing—representing the mental states of others (Saxe and Kanwisher, 2003; Schurz et al., 2020; Van Overwalle and Baetens, 2009). The mPFC is implicated in social-emotional event representation and simulation (Benoit et al., 2011; Schacter et al., 2017). Finally, the amygdala is well-known for its central role in emotion perception (Cheung et al., 2019; LeDoux, 2007), especially fear and anxiety (Davis, 1992). Unsurprisingly, the amygdala showed the strongest evidence for emotion specificity out of all regions.

However, strong emotion specificity was also found in the left TP. The TP is implicated in high-level social-emotional processing, receiving affective input from the amygdala and mPFC (Olson et al., 2007; Ross and Olson, 2010). In our meta-analysis, this region was not consistently engaged in real emotion perception, suggesting that it stores highly abstract social-emotional representations. Moreover, although the TP is located in the ATL, our findings support the view that the TP is not cross-modal, but emotion-specific (cf. Binder and Desai, 2011). Crucially, however, this result does not refute the idea that the ATL contains the cross-modal hub of the conceptual system. It only implies that the cross-modal hub is not located in the TP, but it could be located in other parts of the ATL. Indeed, some previous work suggests that the most critical and modality-invariant hub region is located in the anterior fusiform and/or inferior temporal gyri (Binney et al., 2010; Lambon Ralph et al., 2016; Mion et al., 2010).

In contrast to the other modalities, emotion is a modality of *internal*, not external, perception. Nonetheless, emotion constitutes a crucial type of experiential information for many concepts, especially abstract concepts (Kousta et al., 2011; Ulrich et al., in press). Abstract concepts like ‘love’, ‘argument’ or ‘nightmare’ have strong associations to affective experience (Kiefer and Harpaintner, 2020; Vigliocco et al., 2014). Additional modalities of internal experience, such as introspection and mentalizing (Barsalou and Wiemer-Hastings, 2005; Borghi et al., 2019) as well as social constellations and interactions (Wilson-Mendenhall et al., 2013) might also contribute to the grounding of abstract concepts. Future studies and meta-analyses of (abstract) conceptual representation may consider a wider range of internal perceptual modalities.

*Conceptual processing engages high-level, rather than low-level, perceptual-motor regions* Overall, we found strong evidence for a consistent involvement of modality-specific perceptual-motor areas in conceptual processing. Importantly, however, conceptual processing mainly recruited *high-level* (e.g. secondary or association), rather than low-level (e.g. primary) regions of the modality-specific perceptual-motor systems. For instance, sound-related conceptual-processing engaged the right pSTS, not primary auditory cortex. This indicates that low-level perceptual-motor areas are not consistently recruited across conceptual tasks (Fernandino et al., 2016a; Martin, 2016; Thompson-Schill, 2003). However, several individual studies found conceptual effects in low-level perceptual-motor areas (e.g. Harpaintner et al., 2020; Hauk et al., 2004). As a potential explanation for the lack of consistent recruitment, the engagement of low-level perceptual-motor areas might be particularly task-dependent. Various authors propose that low-level perceptual-motor areas are selectively engaged when the task explicitly requires the retrieval of detailed perceptual-motor information (Binder and Desai, 2011; Kemmerer, 2015; Willems and Casasanto, 2011). This view is supported by several functional neuroimaging studies (Hoenig et al., 2008; Hsu et al., 2011; Kuhnke et al., 2020b; van Dam et al., 2012). Moreover, low-level perceptual-motor areas may influence the activity of higher-level cortical areas via functional connections (e.g. Kuhnke et al., 2021), even if low-level areas are not strongly activated themselves (Fiori et al., 2018; Ward et al., 2010). More generally, the task dependence of the retrieval of perceptual-motor features, and of the resulting engagement of modality-specific areas is a crucial issue for theories of conceptual processing (Kiefer and Pulvermüller, 2012; Kuhnke et al., 2020b; Yee and Thompson-Schill, 2016). Future meta-analyses could directly compare brain activation during implicit tasks (where conceptual access is merely incidental; e.g. lexical decision) vs. explicit tasks (which require the retrieval of conceptual-semantic information; e.g. semantic decision).

Notably, not all perceptual-motor areas were modality-specific, and vice versa. This indicates that perceptual-motor involvement and modality specificity are two distinct issues, and it was crucial to analyze them separately. As an example, the left mid-FG contained a fine-grained parcellation into modality-specific regions for color (ventral), motion (dorsal), shape (anterior), and action (middle) (see Figure 9A). However, none of these regions overlapped with real perceptual-motor experience. The FG regions engaged in conceptual processing generally lay directly anterior to the regions engaged in perception or action. This finding is in line with the “anterior shift” hypothesis that brain regions involved in conceptual processing often lie directly anterior to perceptual-motor areas, potentially reflecting abstraction from basic perceptual-motor information (Thompson-Schill, 2003).

### Multimodal convergence zones for conceptual processing

In addition to modality-specific areas, we identified several multimodal convergence zones that were consistently engaged for conceptual processing related to multiple modalities. In particular, the left IPL and pMTG were robustly engaged for 3 modalities: action, sound, and motion. These “trimodal” areas were surrounded by “bimodal” areas engaged for 2 modalities. Bimodal areas were in turn surrounded by modality-specific areas. This concentric anatomical organization suggests a neural hierarchy, where modality-specific areas converge onto bimodal areas which converge onto trimodal areas (Damasio, 1989; Margulies et al., 2016; Mesulam, 1998). Such a hierarchy of convergence zones may implement multiple levels of abstraction from low-level perceptual-motor information, in line with several previous proposals (Binder and Desai, 2011; Fernandino et al., 2016a; Kuhnke et al., 2021, 2020b; Simmons and Barsalou, 2003).

As an alternative explanation, could multimodal overlap reflect domain-general executive control regions? We believe this to be highly unlikely. First, if multimodal areas were indeed domain-general, they should be engaged for all modalities, not just two (bimodal) or three (trimodal). Second, contrasts included in our meta-analysis largely did not differ in executive demand, but compared two well-matched experimental conditions to isolate modality-specific activity (e.g. action > non-action words). While low-level contrasts (e.g. action words > rest) could differ in control demands, our supplementary analysis without such contrasts yielded highly similar results. Third, control regions are expected to show stronger activation for harder tasks (Noonan et al., 2013). In contrast, a recent large-scale fMRI study (N = 172) revealed that multimodal IPL shows the *opposite* relationship: lower activity for harder tasks (Kuhnke et al., 2022). Moreover, left IPL was not engaged in a recent meta-analysis of “semantic control”—the controlled retrieval of conceptual information (Jackson, 2021). In contrast to left IPL, however, left pMTG is robustly recruited for semantic control (Hodgson et al., 2021; Jackson, 2021). It is possible that left pMTG supports the controlled retrieval of conceptual representations, rather than conceptual representation *per se*.

As a further alternative, could multimodal overlap reflect spreading of activation from one modality to another (e.g. the sound of a dog reactivates its visual shape; Reilly et al., 2016a)? While such “cross-modality spreading” cannot be completely excluded, it is unlikely to explain all multimodal activations, especially in the trimodal IPL and pMTG. Individual studies found multimodal effects in left IPL and pMTG, even when the individual modalities were controlled for (Kuhnke et al., 2020b; Tong et al., 2022). Many experiments included in this meta-analysis similarly isolated modality-specific activity, while controlling for other modalities (e.g. Fernandino et al., 2016b; Goldberg et al., 2006). However, some bimodal activations might reflect the retrieval of a common knowledge type that is relevant for both modalities. For instance, overlap between action and visual shape may either reflect genuine bimodal visuo-tactile shape representations (Amedi et al., 2001) or visual shape information that is also retrieved during object-directed actions (van Elk et al., 2014).

Indeed, our results suggest that there are numerous multimodal areas involving action, and less involving combinations of other modalities. This finding supports the view that action is a core component of human cognition, and action and perception are tightly interlinked (Buxbaum et al., 2005b; Tomasello et al., 2017). However, our dataset for action-related conceptual processing also comprised more experiments (N = 74) than other modalities. Thus, analyses on action had a higher statistical power than other modalities, which might also increase the likelihood to detect overlap. Future work may identify additional multimodal convergence zones for other combinations of modalities.

#### Multimodal vs. amodal hubs

Notably, the anterior temporal lobe (ATL) did not emerge as a multimodal convergence zone, even though the ATL is widely considered as the key cross-modal hub of the conceptual system (for a review, see Lambon Ralph et al., 2016). In support of this view, evidence from semantic dementia (Jefferies, 2013; Patterson et al., 2007), functional neuroimaging (Rice et al., 2015; Visser et al., 2010), and TMS (Pobric et al., 2010a, 2010b) indicates a crucial role of the ATL in processing various types of conceptual information. As a potential explanation for the lack of consistent ATL engagement in our meta-analysis, we propose that the ATL is an “amodal” hub. That is, the ATL completely abstracts away from modality-specific perceptual-motor information to highly abstract conceptual representations (Lambon Ralph et al., 2010; Patterson and Lambon Ralph, 2016). This renders the “amodal” ATL insensitive to modality-specific conceptual content. In other words, we assume the ATL to be equally involved in the processing of all concepts (Jefferies, 2013; Lambon Ralph et al., 2016). However, our meta-analysis focused on contrasts that aim to isolate conceptual processing related to a certain perceptual-motor modality (e.g. action > non-action words). As the ATL is equally engaged for both sides of the contrast (e.g. for both action and non-action words), ATL activation is canceled out. As an alternative explanation, the ATL is known to suffer from susceptibility-induced signal dropout in fMRI (Devlin et al., 2000; Weiskopf et al., 2006). Therefore, it is possible that many studies could not measure ATL activity with a sufficient signal-to-noise ratio to detect modality-specific effects. Notably, however, ATL activation is consistently observed in meta-analyses of general conceptual contrasts (e.g. words > pseudowords; Binder et al., 2009; Hodgson et al., 2021; Jackson, 2021). Thus, it seems more likely that the ATL was not engaged for individual modalities as the ATL is amodal. Finally, while both left and right ATL seem to be engaged in conceptual processing, there may be subtle differences in hemispheric specialization (Jung and Lambon Ralph, 2016). For instance, a previous meta-analysis found ATL activations to be left-lateralized for written input and word retrieval (Rice et al., 2015).

In contrast to the “amodal” ATL, “multimodal” hubs like left IPL and pMTG retain modality-specific perceptual-motor information about the individual modalities that they bind (Fernandino et al., 2016b; Kuhnke et al., 2022, 2021, 2020b; Reilly et al., 2016b; Seghier, 2013). Hence, these regions are sensitive to modality-specific conceptual information related to several modalities.

The multimodal—amodal hub theory is supported by several studies. For example, Kuhnke et al. (2020b) demonstrated that left IPL and pMTG respond to both sound and action features of concepts when these are task-relevant. In contrast, the ATL was not engaged for individual features, but for general conceptual information (words > pseudowords). In a follow-up study (Kuhnke et al., 2021), left IPL was functionally coupled with auditory brain regions during sound feature retrieval, and with somatomotor regions during action feature retrieval. In contrast, the ATL interacted with other high-level cross-modal areas, but not modality-specific cortices. In line with these results, Fernandino et al. (2016a) found that activity in the IPL during word reading correlated with the strength of sensory-motor associations for all modalities tested (action, sound, shape, color, motion). Again, ATL activity did not correlate with individual sensory-motor associations. Finally, TMS over left IPL (Ishibashi et al., 2011; Kuhnke et al., 2020a; Pobric et al., 2010a) and pMTG (Davey et al., 2015; Whitney et al., 2012) can selectively disrupt the retrieval of individual task-relevant semantic features. In contrast, TMS over ATL typically impairs semantic processing for all types of concepts (Pobric et al., 2010a, 2010b).

### Evidence for hybrid theories of conceptual processing

Theories of conceptual representation can be organized on a continuum between strong embodied and strong amodal views (Kiefer and Harpaintner, 2020; Meteyard et al., 2012). Strong embodied views—the extreme version of grounded cognition theories—hold that conceptual processing relies exclusively on distributed and interconnected modality-specific perceptual-motor areas (e.g. Allport, 1985). In contrast, strong amodal views assume that concepts consist entirely of abstract, amodal symbols represented outside the perceptual-motor systems (Fodor, 1975; Pylyshyn, 1984).

Our results oppose both of these extremes. In contrast to strong embodied views, we found consistent engagement of multimodal convergence zones, not only modality-specific perceptual-motor regions. Contrary to strong amodal views, we found that conceptual processing robustly recruits modality-specific perceptual-motor areas. Moreover, conceptual processing engages multimodal, not only amodal, hubs. Taken together, our results support so-called “hybrid theories” of conceptual processing, which assume an involvement of both modality-specific perceptual-motor cortices and cross-modal convergence zones (Binder and Desai, 2011; Fernandino et al., 2016a; Kiefer and Harpaintner, 2020; Reilly et al., 2016b; Simmons and Barsalou, 2003).

Based on our findings, we now propose a new model of the conceptual system—a refined and extended version of our previous account (Kuhnke et al., 2021, 2020b). According to this model, conceptual processing relies on a hierarchical neural architecture from modality-specific to bimodal to trimodal regions, up to an amodal hub in the ATL (Figure 10). At the functional level, the neural hierarchy implements abstraction of conceptual representations from basic perceptual-motor information (Binder and Desai, 2011; Fernandino et al., 2016a; Kiefer and Harpaintner, 2020). At the structural level, representational convergence is implemented via a concentric anatomical organization, where trimodal areas are surrounded by bimodal areas which are surrounded by modality-specific areas (Damasio, 1989; Margulies et al., 2016; Mesulam, 1998). Our model can account for all key results of the current meta-analysis: (1) The consistent recruitment of modality-specific perceptual-motor regions, (2) the overlap of multiple modalities in multimodal (i.e. bimodal and trimodal) convergence zones, and (3) the absence of ATL recruitment for modality-specific contrasts, despite overwhelming evidence for a crucial role of the ATL in semantic cognition. Moreover, our model is supported by a recent computational modeling study which revealed that the core functions of the conceptual system—conceptual abstraction and flexibility—are best achieved by a hierarchical multi-level architecture composed of a modality-specific layer, an intermediate layer (∼multimodal regions), and a single top-level hub (∼amodal ATL) (Jackson et al., 2021; also see Garagnani and Pulvermüller, 2016).

**Figure 10.**
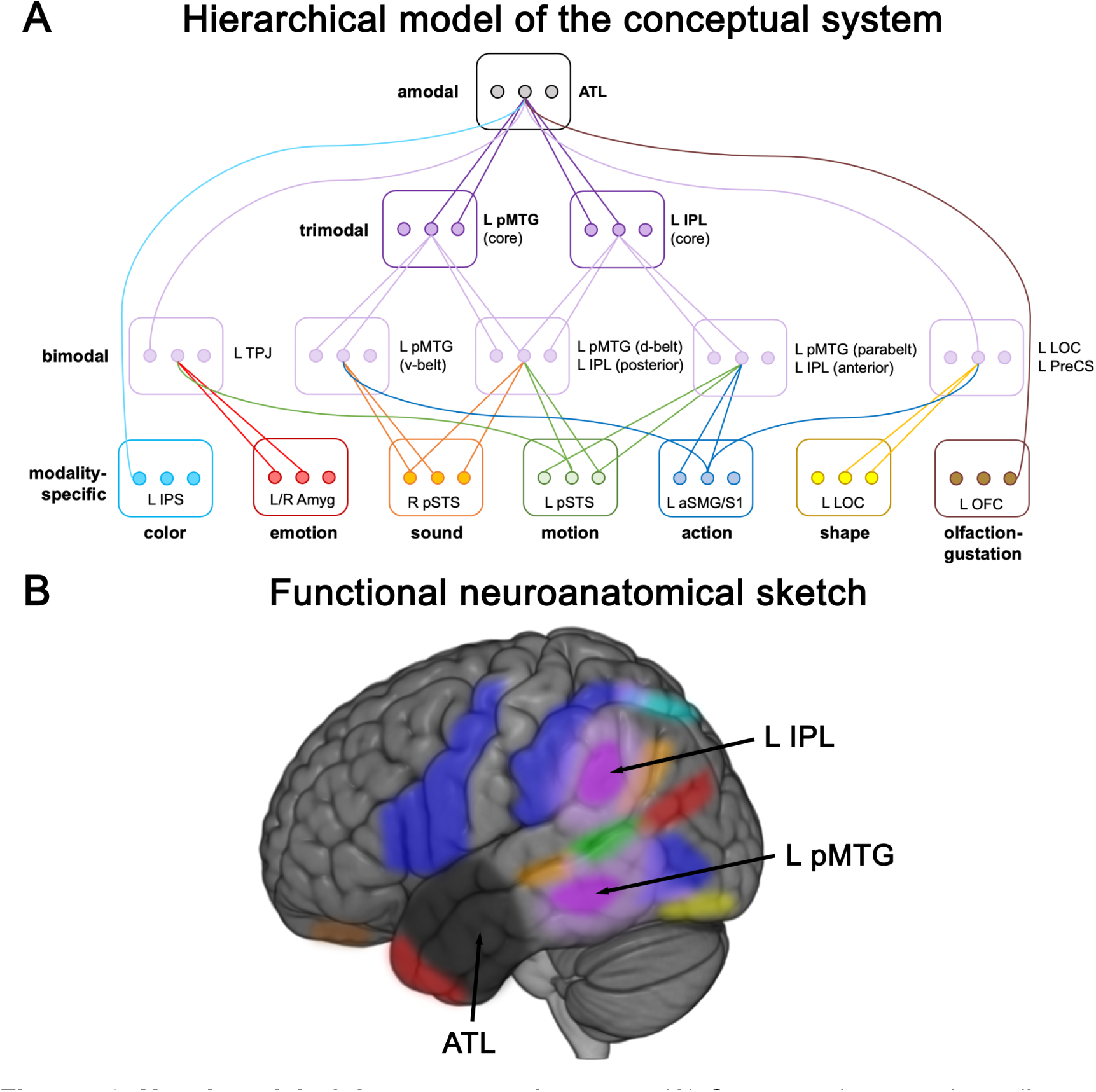
Novel model of the conceptual system. (A) Conceptual processing relies on a hierarchical neural architecture from modality-specific to bimodal to trimodal areas, up to an amodal hub. Boxes represent brain regions and dots represent individual representational units that converge onto a more abstract representation at a higher level. (B) Functional neuroanatomical sketch illustrating the proposed locations of the different components of our model on the cerebral cortex. The color code is identical to panel A.

Our model is related to two prominent theories of conceptual representation: The “hub-and-spokes” (Lambon Ralph et al., 2016; Patterson et al., 2007) and “embodied abstraction” (Binder and Desai, 2011; Fernandino et al., 2016a) models. According to the hub-and-spokes model, modality-specific “spoke” regions converge onto a single cross-modal “hub” in the ATL. “Graded” versions of the hub-and-spokes model suggest that different ATL subregions may be weighted towards (combinations of) individual modalities depending on their proximity to and connectivity with modality-specific cortices (Binney et al., 2012; Lambon Ralph et al., 2016). In contrast, the embodied abstraction model proposes a hierarchy of cross-modal convergence zones in the inferior parietal, temporal, and medial prefrontal cortices. In line with embodied abstraction, our model assumes multiple levels of cross-modal convergence zones. In line with the hub-and-spokes model, our model proposes the ATL to constitute the most abstract cross-modal hub. However, in contrast to both theories, our model distinguishes among cross-modal areas between “multimodal” regions which retain modality-specific information and the “amodal” ATL which does not. In addition, our model makes more precise functional-anatomical predictions: (1) Multimodal areas are restricted to bimodal and trimodal zones, whereas no multimodal area binds more than three modalities. (2) Trimodal areas are exclusively located in the left IPL and pMTG. (3) Trimodal, bimodal and modality-specific areas show a concentric anatomical organization. (4) Only the ATL functions as amodal hub. These predictions could be directly tested in future research to further develop and refine theories of conceptual representation in the human brain.

For methodological reasons, this meta-analysis selectively included voxel-wise activation-based neuroimaging analyses. Two other types of analyses that provide crucial information about the neural bases of conceptual processing are multivariate pattern analyses (MVPA) and connectivity analyses. MVPA—including decoding and representational similarity analysis (RSA)—tests for information represented in fine-grained, multi-voxel activity patterns (Haxby et al., 2014; Norman et al., 2006). RSA has recently been used to relate computational models of semantics to the brain, which revealed that a grounded perceptual-motor model better explains brain representations (including in multimodal regions) than taxonomic categories or distributional information (Fernandino et al., 2022; Tong et al., 2022). These findings clearly corroborate our results and our model. Moreover, functional and effective connectivity analyses can assess how the various modality-specific, multimodal and amodal brain regions work together during conceptual tasks (Chai et al., 2016; Chiou et al., 2018; Jackson et al., 2016; Kuhnke et al., 2021; Wang et al., 2017). In addition, electrophysiological measures with a high temporal resolution (such as EEG/MEG) provide invaluable information about the time course of conceptual knowledge retrieval (Hauk, 2016; Kiefer et al., 2022). Finally, neuropsychological lesion studies and non-invasive brain stimulation are essential to assess the causal relevance of different brain structures for conceptual processing (Bergmann and Hartwigsen, 2021; Price and Friston, 2002). In particular, evidence for a causal role of modality-specific perceptual-motor regions is still scarce, especially for modalities other than action (Hauk and Tschentscher, 2013; Trumpp et al., 2013; Vukovic et al., 2017). Only through the combination of these complementary sources of evidence can we arrive at a comprehensive understanding of the neural bases of conceptual processing.

## Conclusion

In conclusion, this meta-analysis of >200 functional neuroimaging studies revealed that conceptual processing robustly recruits both modality-specific perceptual-motor regions and multimodal convergence zones. These results support “hybrid theories” of conceptual processing which propose an involvement of both modality-specific and cross-modal cortices. We propose a novel model of the conceptual system, according to which conceptual processing relies on a hierarchical neural architecture from modality-specific to bimodal to trimodal areas (left IPL, pMTG) up to an amodal hub in the ATL.

## Statements & Declarations

## Acknowledgements

We wish to thank Dana Richter and Lisa Reimund for their help in literature search and data extraction from relevant articles.

## Author Contributions (CRediT)

**Philipp Kuhnke**: Conceptualization, Investigation, Data curation, Formal analysis, Methodology, Visualization, Writing—original draft, Writing—review and editing;

**Marie C. Beaupain**: Investigation, Data curation, Methodology, Writing—review and editing;

**Johannes Arola**: Investigation, Data curation, Methodology;

**Markus Kiefer**: Conceptualization, Writing—review and editing.

**Gesa Hartwigsen**: Conceptualization, Funding acquisition, Supervision, Project administration, Writing—review and editing.

## Data Statement

All meta-analytic maps are openly available via the ANIMA database (Reid et al., 2016): https://anima.fz-juelich.de/studies/Kuhnke_2023_Conceptual_Processing.

## Appendix

**Table A1.** Acronyms for brain regions.

|  |  |  |  |
| --- | --- | --- | --- |
| a (prefix) | anterior | PCC | posterior cingulate cortex |
| p (prefix) | posterior | PFC | prefrontal cortex |
| d (prefix) | dorsal | mPFC | medial PFC |
| v (prefix) | ventral | dmPFC | dorsomedial PFC |
| A1 | primary auditory cortex | vmPFC | ventromedial PFC |
| ACC | anterior cingulate cortex | PMC | premotor cortex |
| AG | angular gyrus | PMd | dorsal PMC |
| ATL | anterior temporal lobe | PMv | ventral PMC |
| FG | fusiform gyrus | PreCS | precentral sulcus |
| IFG | inferior frontal gyrus | S1 | primary somatosensory cortex |
| IPL | inferior parietal lobe | SMA | supplementary motor area |
| IPS | intraparietal sulcus | SMG | supramarginal gyrus |
| LOC | lateral occipital cortex | SPL | superior parietal lobe |
| LTO | lateral temporal-occipital junction | STG | superior temporal gyrus |
| M1 | primary motor cortex | STS | superior temporal sulcus |
| MCC | middle cingulate cortex | TP | temporal pole |
| MFG | middle frontal gyrus | TPJ | temporoparietal junction |
| MTG | middle temporal gyrus | V1/V2 | primary/secondary visual cortex |
| OFC | orbitofrontal cortex | V5/MT | middle temporal visual area |

## References

Allport, D.A., 1985. Distributed memory, modular subsystems and dysphasia, in: Newman, S.K., Epstein, R. (Eds.), Current Perspectives in Dysphasia. Churchill Livingstone.

Amedi, A., Malach, R., Hendler, T., Peled, S., Zohary, E., 2001. Visuo-haptic object-related activation in the ventral visual pathway. Nat. Neurosci. 4, 324–330. https://doi.org/10.1038/85201

Barsalou, L.W., 2008. Grounded Cognition. Annu. Rev. Psychol. 59, 617–645. https://doi.org/10.1146/annurev.psych.59.103006.093639

Barsalou, L.W., Wiemer-Hastings, K., 2005. Situating Abstract Concepts, in: Pecher, D., Zwaan, R.A. (Eds.), Grounding Cognition: The Role of Perception and Action in Memory, Language, and Thinking. pp. 129–163. https://doi.org/10.1017/CBO9780511499968

Beauchamp, M.S., 1999. An fMRI Version of the Farnsworth-Munsell 100-Hue Test Reveals Multiple Color-selective Areas in Human Ventral Occipitotemporal Cortex. Cereb. Cortex 9, 257–263. https://doi.org/10.1093/cercor/9.3.257

Bedny, M., Caramazza, A., Grossman, E., Pascual-Leone, A., Saxe, R., 2008. Concepts are more than percepts: the case of action verbs. J Neurosci 28, 11347–11353. https://doi.org/10.1523/JNEUROSCI.3039-08.2008

Belin, P., Zatorre, R.J., Lafaille, P., Ahad, P., Pike, B., 2000. Voice-selective areas in human auditory cortex. Nature 403, 309–312. https://doi.org/10.1038/35002078

Benoit, R.G., Gilbert, S.J., Burgess, P.W., 2011. A Neural Mechanism Mediating the Impact of Episodic Prospection on Farsighted Decisions. J. Neurosci. 31, 6771–6779. https://doi.org/10.1523/JNEUROSCI.6559-10.2011

Bergmann, T.O., Hartwigsen, G., 2021. Inferring Causality from Noninvasive Brain Stimulation in Cognitive Neuroscience. J. Cogn. Neurosci. 33, 195–225. https://doi.org/10.1162/jocn_a_01591

Binder, J.R., Desai, R.H., 2011. The neurobiology of semantic memory. Trends Cogn. Sci. 15, 527–536. https://doi.org/10.1016/j.tics.2011.10.001

Binder, J.R., Desai, R.H., Graves, W.W., Conant, L.L., 2009. Where Is the Semantic System? A Critical Review and Meta-Analysis of 120 Functional Neuroimaging Studies. Cereb. Cortex 19, 2767–2796. https://doi.org/10.1093/cercor/bhp055

Binney, R.J., Embleton, K. V., Jefferies, E., Parker, G.J.M., Lambon Ralph, M.A., 2010. The ventral and inferolateral aspects of the anterior temporal lobe are crucial in semantic memory: Evidence from a novel direct comparison of distortion-corrected fMRI, rTMS, and semantic dementia. Cereb. Cortex 20, 2728–2738. https://doi.org/10.1093/cercor/bhq019

Binney, R.J., Parker, G.J.M., Lambon Ralph, M.A., 2012. Convergent Connectivity and Graded Specialization in the Rostral Human Temporal Lobe as Revealed by Diffusion-Weighted Imaging Probabilistic Tractography. J. Cogn. Neurosci. 24, 1998–2014. https://doi.org/10.1162/jocn_a_00263

Borghi, A.M., Barca, L., Binkofski, F., Castelfranchi, C., Pezzulo, G., Tummolini, L., 2019. Words as social tools: Language, sociality and inner grounding in abstract concepts. Phys. Life Rev. 29, 120–153. https://doi.org/10.1016/j.plrev.2018.12.001

Boussaoud, D., Ungerleider, L.G., Desimone, R., 1990. Pathways for motion analysis: Cortical connections of the medial superior temporal and fundus of the superior temporal visual areas in the macaque. J. Comp. Neurol. 296, 462–495. https://doi.org/10.1002/cne.902960311

Buxbaum, L.J., Johnson-Frey, S.H., Bartlett-Williams, M., 2005a. Deficient internal models for planning hand-object interactions in apraxia. Neuropsychologia 43, 917–929. https://doi.org/10.1016/j.neuropsychologia.2004.09.006

Buxbaum, L.J., Kyle, K.M., Menon, R., 2005b. On beyond mirror neurons: Internal representations subserving imitation and recognition of skilled object-related actions in humans. Cogn. Brain Res. 25, 226–239. https://doi.org/10.1016/j.cogbrainres.2005.05.014

Chai, L.R., Mattar, M.G., Blank, I.A., Fedorenko, E., Bassett, D.S., 2016. Functional Network Dynamics of the Language System. Cereb. Cortex 26, 4148–4159. https://doi.org/10.1093/cercor/bhw238

Chao, L.L., Haxby, J. V, Martin, A., 1999. Attribute-based neural substrates in temporal cortex for perceiving and knowing about objects. Nat. Neurosci. 2, 913–919. https://doi.org/10.1038/13217

Cheung, V.K.M., Harrison, P.M.C., Meyer, L., Pearce, M.T., Haynes, J.-D., Koelsch, S., 2019. Uncertainty and Surprise Jointly Predict Musical Pleasure and Amygdala, Hippocampus, and Auditory Cortex Activity. Curr. Biol. 1–9. https://doi.org/10.1016/j.cub.2019.09.067

Chiou, R., Humphreys, G.F., Jung, J.Y., Lambon Ralph, M.A., 2018. Controlled semantic cognition relies upon dynamic and flexible interactions between the executive ‘semantic control’ and hub-and-spoke ‘semantic representation’ systems. Cortex 103, 100–116. https://doi.org/10.1016/j.cortex.2018.02.018

Cieslik, E.C., Seidler, I., Laird, A.R., Fox, P.T., Eickhoff, S.B., 2016. Different involvement of subregions within dorsal premotor and medial frontal cortex for pro-and antisaccades. Neurosci. Biobehav. Rev. 68, 256–269. https://doi.org/10.1016/j.neubiorev.2016.05.012

Conway, B.R., 2009. Color vision, cones, and color-coding in the cortex. Neuroscientist 15, 274–290. https://doi.org/10.1177/1073858408331369

Culham, J.C., Valyear, K.F., 2006. Human parietal cortex in action. Curr. Opin. Neurobiol. 16, 205–212. https://doi.org/10.1016/j.conb.2006.03.005

Damasio, A.R., 1989. The Brain Binds Entities and Events by Multiregional Activation from Convergence Zones. Neural Comput. 1, 123–132. https://doi.org/10.1162/neco.1989.1.1.123

Davey, J., Cornelissen, P.L., Thompson, H.E., Sonkusare, S., Hallam, G., Smallwood, J., Jefferies, E., 2015. Automatic and Controlled Semantic Retrieval: TMS Reveals Distinct Contributions of Posterior Middle Temporal Gyrus and Angular Gyrus. J. Neurosci. 35, 15230–15239. https://doi.org/10.1523/JNEUROSCI.4705-14.2015

Davis, M., 1992. The Role of the Amygdala in Fear and Anxiety. Annu. Rev. Neurosci. 15, 353–375. https://doi.org/10.1146/annurev.ne.15.030192.002033

de Araujo, I.E.T., Rolls, E.T., Kringelbach, M.L., McGlone, F., Phillips, N., 2003. Taste-olfactory convergence, and the representation of the pleasantness of flavour, in the human brain. Eur. J. Neurosci. 18, 2059–2068. https://doi.org/10.1046/j.1460-9568.2003.02915.x

Desai, R.H., Binder, J.R., Conant, L.L., Seidenberg, M.S., 2010. Activation of Sensory-Motor Areas in Sentence Comprehension. Cereb. Cortex 20, 468–478. https://doi.org/10.1093/cercor/bhp115

Desai, R.H., Reilly, M., van Dam, W., 2018. The multifaceted abstract brain. Philos. Trans. R. Soc. B Biol. Sci. 373, 20170122. https://doi.org/10.1098/rstb.2017.0122

Devlin, J.T., Russell, R.P., Davis, M.H., Price, C.J., Wilson, J., Moss, H.E., Matthews, P.M., Tyler, L.K., 2000. Susceptibility-Induced Loss of Signal: Comparing PET and fMRI on a Semantic Task. Neuroimage 11, 589–600. https://doi.org/10.1006/nimg.2000.0595

Diveica, V., Koldewyn, K., Binney, R.J., 2021. Establishing a Role of the Semantic Control Network in Social Cognitive Processing: A Meta-analysis of Functional Neuroimaging Studies Veronica. bioRxiv Prepr. 1–71. https://doi.org/10.1101/2021.04.01.437961

Eickhoff, S.B., Bzdok, D., Laird, A.R., Kurth, F., Fox, P.T., 2012. Activation likelihood estimation meta-analysis revisited. Neuroimage 59, 2349–2361. https://doi.org/10.1016/j.neuroimage.2011.09.017

Eickhoff, S.B., Bzdok, D., Laird, A.R., Roski, C., Caspers, S., Zilles, K., Fox, P.T., 2011. Co-activation patterns distinguish cortical modules, their connectivity and functional differentiation. Neuroimage 57, 938–949. https://doi.org/10.1016/j.neuroimage.2011.05.021

Eickhoff, S.B., Laird, A.R., Grefkes, C., Wang, L.E., Zilles, K., Fox, P.T., 2009. Coordinate-based activation likelihood estimation meta-analysis of neuroimaging data: A random-effects approach based on empirical estimates of spatial uncertainty. Hum. Brain Mapp. 30, 2907–2926. https://doi.org/10.1002/hbm.20718

Eickhoff, S.B., Nichols, T.E., Laird, A.R., Hoffstaedter, F., Amunts, K., Fox, P.T., Bzdok, D., Eickhoff, C.R., 2016. Behavior, sensitivity, and power of activation likelihood estimation characterized by massive empirical simulation. Neuroimage 137, 70–85. https://doi.org/10.1016/j.neuroimage.2016.04.072

Fernandino, L., Binder, J.R., Desai, R.H., Pendl, S.L., Humphries, C.J., Gross, W.L., Conant, L.L., Seidenberg, M.S., 2016a. Concept Representation Reflects Multimodal Abstraction: A Framework for Embodied Semantics. Cereb. Cortex 26, 2018–2034. https://doi.org/10.1093/cercor/bhv020

Fernandino, L., Humphries, C.J., Conant, L.L., Seidenberg, M.S., Binder, J.R., 2016b. Heteromodal Cortical Areas Encode Sensory-Motor Features of Word Meaning. J. Neurosci. 36, 9763–9769. https://doi.org/10.1523/JNEUROSCI.4095-15.2016

Fernandino, L., Tong, J.Q., Conant, L.L., Humphries, C.J., Binder, J.R., 2022. Decoding the information structure underlying the neural representation of concepts. Proc. Natl. Acad. Sci. U. S. A. 119, 1–11. https://doi.org/10.1073/pnas.2108091119

Fiori, V., Kunz, L., Kuhnke, P., Marangolo, P., Hartwigsen, G., 2018. Transcranial direct current stimulation (tDCS) facilitates verb learning by altering effective connectivity in the healthy brain. Neuroimage 181, 550–559. https://doi.org/10.1016/j.neuroimage.2018.07.040

Fodor, J.A., 1975. The Language of Thought. Harvard University Press, Cambridge, MA.

Fox, P.T., Lancaster, J.L., Laird, A.R., Eickhoff, S.B., 2014. Meta-analysis in human neuroimaging: Computational modeling of large-scale databases. Annu. Rev. Neurosci. 37, 409–434. https://doi.org/10.1146/annurev-neuro-062012-170320

Friederici, A.D., 2012. The cortical language circuit: From auditory perception to sentence comprehension. Trends Cogn. Sci. 16, 262–268. https://doi.org/10.1016/j.tics.2012.04.001

Friederici, A.D., 2002. Towards a neural basis of auditory sentence processing. Trends Cogn. Sci. 6, 78–84.

Galeano Weber, E.M., Peters, B., Hahn, T., Bledowski, C., Fiebach, C.J., 2016. Superior Intraparietal Sulcus Controls the Variability of Visual Working Memory Precision. J. Neurosci. 36, 5623–5635. https://doi.org/10.1523/JNEUROSCI.1596-15.2016

Garagnani, M., Pulvermüller, F., 2016. Conceptual grounding of language in action and perception: A neurocomputational model of the emergence of category specificity and semantic hubs. Eur. J. Neurosci. 43, 721–737. https://doi.org/10.1111/ejn.13145

Giacobbe, C., Raimo, S., Cropano, M., Santangelo, G., 2022. Neural correlates of embodied action language processing: a systematic review and meta-analytic study. Brain Imaging Behav. https://doi.org/10.1007/s11682-022-00680-3

Goldberg, R.F., Perfetti, C.A., Schneider, W., 2006. Perceptual Knowledge Retrieval Activates Sensory Brain Regions. J. Neurosci. 26, 4917–4921. https://doi.org/10.1523/JNEUROSCI.5389-05.2006

Goldenberg, G., 2009. Apraxia and the parietal lobes. Neuropsychologia 47, 1449–1459. https://doi.org/10.1016/j.neuropsychologia.2008.07.014

Haaland, K.Y., Harrington, D.L., Knight, R.T., 2000. Neural representations of skilled movement. Brain 123, 2306–2313. https://doi.org/10.1093/brain/123.11.2306

Halpern, A.R., Zatorre, R.J., Bouffard, M., Johnson, J.A., 2004. Behavioral and neural correlates of perceived and imagined musical timbre. Neuropsychologia 42, 1281– 1292. https://doi.org/10.1016/j.neuropsychologia.2003.12.017

Hardwick, R.M., Caspers, S., Eickhoff, S.B., Swinnen, S.P., 2018. Neural correlates of action: Comparing meta-analyses of imagery, observation, and execution. Neurosci. Biobehav. Rev. 94, 31–44. https://doi.org/10.1016/j.neubiorev.2018.08.003

Harpaintner, M., Sim, E., Trumpp, N.M., Ulrich, M., Kiefer, M., 2020. The grounding of abstract concepts in the motor and visual system: An fMRI study. Cortex 124, 1–22. https://doi.org/10.1016/j.cortex.2019.10.014

Hauk, O., 2016. Only time will tell – why temporal information is essential for our neuroscientific understanding of semantics. Psychon. Bull. Rev. 23, 1072–1079. https://doi.org/10.3758/s13423-015-0873-9

Hauk, O., Johnsrude, I., Pulvermüller, F., 2004. Somatotopic Representation of Action Words in Human Motor and Premotor Cortex. Neuron 41, 301–307. https://doi.org/10.1016/S0896-6273(03)00838-9

Hauk, O., Tschentscher, N., 2013. The Body of Evidence: What Can Neuroscience Tell Us about Embodied Semantics? Front. Psychol. 4, 1–14. https://doi.org/10.3389/fpsyg.2013.00050

Haxby, J. V., Connolly, A.C., Guntupalli, J.S., 2014. Decoding Neural Representational Spaces Using Multivariate Pattern Analysis. Annu. Rev. Neurosci. 37, 435–456. https://doi.org/10.1146/annurev-neuro-062012-170325

Hodgson, V.J., Lambon Ralph, M.A., Jackson, R.L., 2021. Multiple dimensions underlying the functional organization of the language network. Neuroimage 241, 118444. https://doi.org/10.1016/j.neuroimage.2021.118444

Hoenig, K., Sim, E.-J., Bochev, V., Herrnberger, B., Kiefer, M., 2008. Conceptual Flexibility in the Human Brain: Dynamic Recruitment of Semantic Maps from Visual, Motor, and Motion-related Areas. J. Cogn. Neurosci. 20, 1799–1814. https://doi.org/10.1162/jocn.2008.20123

Hsu, N.S., Frankland, S.M., Thompson-Schill, S.L., 2012. Chromaticity of color perception and object color knowledge. Neuropsychologia 50, 327–333. https://doi.org/10.1016/j.neuropsychologia.2011.12.003

Hsu, N.S., Kraemer, D.J.M., Oliver, R.T., Schlichting, M.L., Thompson-Schill, S.L., 2011. Color, context, and cognitive style: variations in color knowledge retrieval as a function of task and subject variables. J. Cogn. Neurosci. 23, 2544–57. https://doi.org/10.1162/jocn.2011.21619

Ishibashi, R., Lambon Ralph, M.A., Saito, S., Pobric, G., 2011. Different roles of lateral anterior temporal lobe and inferior parietal lobule in coding function and manipulation tool knowledge: Evidence from an rTMS study. Neuropsychologia 49, 1128–1135. https://doi.org/10.1016/j.neuropsychologia.2011.01.004

Jackson, R.L., 2021. The neural correlates of semantic control revisited. Neuroimage 224. https://doi.org/10.1016/j.neuroimage.2020.117444

Jackson, R.L., Hoffman, P., Pobric, G., Lambon Ralph, M.A., 2016. The semantic network at work and rest: Differential connectivity of anterior temporal lobe subregions. J. Neurosci. 36, 1490–1501. https://doi.org/10.1523/JNEUROSCI.2999-15.2016

Jackson, R.L., Rogers, T.T., Lambon Ralph, M.A., 2021. Reverse-engineering the cortical architecture for controlled semantic cognition. Nat. Hum. Behav. 860528. https://doi.org/10.1038/s41562-020-01034-z

Jefferies, E., 2013. The neural basis of semantic cognition: Converging evidence from neuropsychology, neuroimaging and TMS. Cortex 49, 611–625. https://doi.org/10.1016/j.cortex.2012.10.008

Johnson-Frey, S.H., 2004. The neural bases of complex tool use in humans. Trends Cogn. Sci. 8, 71–78. https://doi.org/10.1016/j.tics.2003.12.002

Jung, J., Lambon Ralph, M.A., 2016. Mapping the Dynamic Network Interactions Underpinning Cognition: A cTBS-fMRI Study of the Flexible Adaptive Neural System for Semantics. Cereb. Cortex 26, 3580–3590. https://doi.org/10.1093/cercor/bhw149

Kemmerer, D., 2015. Are the motor features of verb meanings represented in the precentral motor cortices? Yes, but within the context of a flexible, multilevel architecture for conceptual knowledge. Psychon. Bull. Rev. 22, 1068–1075. https://doi.org/10.3758/s13423-014-0784-1

Kemmerer, D., 2014. Cognitive neuroscience of language. Psychology Press, New York and London.

Kiefer, M., Barsalou, L.W., 2013. Grounding the Human Conceptual System in Perception, Action, and Internal States, in: Prinz, W., Beisert, M., Herwig, A. (Eds.), Tutorials in Action Science. The MIT Press, Cambridge, pp. 381–407. https://doi.org/10.7551/mitpress/9780262018555.003.0015

Kiefer, M., Harpaintner, M., 2020. Varieties of abstract concepts and their grounding in perception or action. Open Psychol. 2, 119–137. https://doi.org/10.1515/psych-2020- 0104

Kiefer, M., Pielke, L., Trumpp, N.M., 2022. Differential temporo-spatial pattern of electrical brain activity during the processing of abstract concepts related to mental states and verbal associations. Neuroimage 252, 119036. https://doi.org/10.1016/j.neuroimage.2022.119036

Kiefer, M., Pulvermüller, F., 2012. Conceptual representations in mind and brain: Theoretical developments, current evidence and future directions. Cortex 48, 805–825. https://doi.org/10.1016/j.cortex.2011.04.006

Kiefer, M., Sim, E.-J., Herrnberger, B., Grothe, J., Hoenig, K., 2008. The Sound of Concepts: Four Markers for a Link between Auditory and Conceptual Brain Systems. J. Neurosci. 28, 12224–12230. https://doi.org/10.1523/JNEUROSCI.3579-08.2008

Kompa, N.A., 2021. Language and embodiment—Or the cognitive benefits of abstract representations. Mind Lang. 36, 27–47. https://doi.org/10.1111/mila.12266

Kompa, N.A., Mueller, J.L., 2020. How Abstract (Non-embodied) Linguistic Representations Augment Cognitive Control. Front. Psychol. 11. https://doi.org/10.3389/fpsyg.2020.01597

Kourtzi, Z., Erb, M., Grodd, W., Bülthoff, H.H., 2003. Representation of the perceived 3-D object shape in the human lateral occipital complex. Cereb. Cortex 13, 911–920. https://doi.org/10.1093/cercor/13.9.911

Kousta, S.T., Vigliocco, G., Vinson, D.P., Andrews, M., Del Campo, E., 2011. The Representation of Abstract Words: Why Emotion Matters. J. Exp. Psychol. Gen. 140, 14–34. https://doi.org/10.1037/a0021446

Kuhnke, P., Beaupain, M.C., Cheung, V.K.M., Weise, K., Kiefer, M., Hartwigsen, G., 2020a. Left posterior inferior parietal cortex causally supports the retrieval of action knowledge. Neuroimage 219, 117041. https://doi.org/10.1016/j.neuroimage.2020.117041

Kuhnke, P., Chapman, C.A., Cheung, V.K.M., Turker, S., Graessner, A., Martin, S., Williams, K.A., Hartwigsen, G., 2022. The role of the angular gyrus in semantic cognition: a synthesis of five functional neuroimaging studies. Brain Struct. Funct. https://doi.org/10.1007/s00429-022-02493-y

Kuhnke, P., Kiefer, M., Hartwigsen, G., 2021. Task-Dependent Functional and Effective Connectivity during Conceptual Processing. Cereb. Cortex 31, 3475–3493. https://doi.org/10.1093/cercor/bhab026

Kuhnke, P., Kiefer, M., Hartwigsen, G., 2020b. Task-Dependent Recruitment of Modality-Specific and Multimodal Regions during Conceptual Processing. Cereb. Cortex 30, 3938–3959. https://doi.org/10.1093/cercor/bhaa010

Lambon Ralph, M.A., 2013. Neurocognitive insights on conceptual knowledge and its breakdown. Philos. Trans. R. Soc. B Biol. Sci. 369, 20120392–20120392. https://doi.org/10.1098/rstb.2012.0392

Lambon Ralph, M.A., Jefferies, E., Patterson, K., Rogers, T.T., 2016. The neural and computational bases of semantic cognition. Nat. Rev. Neurosci. 18, 42–55. https://doi.org/10.1038/nrn.2016.150

Lambon Ralph, M.A., Sage, K., Jones, R.W., Mayberry, E.J., 2010. Coherent concepts are computed in the anterior temporal lobes. Proc. Natl. Acad. Sci. 107, 2717–2722. https://doi.org/10.1073/pnas.0907307107

Lancaster, J.L., Tordesillas-Gutiérrez, D., Martinez, M., Salinas, F., Evans, A., Zilles, K., Mazziotta, J.C., Fox, P.T., 2007. Bias between MNI and Talairach coordinates analyzed using the ICBM-152 brain template. Hum. Brain Mapp. 28, 1194–1205. https://doi.org/10.1002/hbm.20345

Langner, R., Eickhoff, S.B., 2013. Sustaining attention to simple tasks: A meta-analytic review of the neural mechanisms of vigilant attention. Psychol. Bull. 139, 870–900. https://doi.org/10.1037/a0030694

Larsson, J., Heeger, D.J., 2006. Two Retinotopic Visual Areas in Human Lateral Occipital Cortex. J. Neurosci. 26, 13128–13142. https://doi.org/10.1523/JNEUROSCI.1657-06.2006

LeDoux, J., 2007. The amygdala. Curr. Biol. 17, R868–R874. https://doi.org/10.1016/j.cub.2007.08.005

Lewis, J.W., 2006. Cortical Networks Related to Human Use of Tools. Neurosci. 12, 211– 231. https://doi.org/10.1177/1073858406288327

Lewis, J.W., Talkington, W.J., Walker, N.A., Spirou, G.A., Jajosky, A., Frum, C., Brefczynski-Lewis, J.A., 2009. Human Cortical Organization for Processing Vocalizations Indicates Representation of Harmonic Structure as a Signal Attribute. J. Neurosci. 29, 2283– 2296. https://doi.org/10.1523/JNEUROSCI.4145-08.2009

Lewis, J.W., Wightman, F.L., Brefczynski, J.A., Phinney, R.E., Binder, J.R., DeYoe, E.A., 2004. Human Brain Regions Involved in Recognizing Environmental Sounds. Cereb. Cortex 14, 1008–1021. https://doi.org/10.1093/cercor/bhh061

Mahon, B.Z., 2015. What is embodied about cognition? Lang. Cogn. Neurosci. 30, 420–429. https://doi.org/10.1080/23273798.2014.987791

Mahon, B.Z., Caramazza, A., 2008. A critical look at the embodied cognition hypothesis and a new proposal for grounding conceptual content. J. Physiol. 102, 59–70. https://doi.org/10.1016/j.jphysparis.2008.03.004

Malikovic, A., Amunts, K., Schleicher, A., Mohlberg, H., Kujovic, M., Palomero-Gallagher, N., Eickhoff, S.B., Zilles, K., 2016. Cytoarchitecture of the human lateral occipital cortex: mapping of two extrastriate areas hOc4la and hOc4lp. Brain Struct. Funct. 221, 1877– 1897. https://doi.org/10.1007/s00429-015-1009-8

Margulies, D.S., Ghosh, S.S., Goulas, A., Falkiewicz, M., Huntenburg, J.M., Langs, G., Bezgin, G., Eickhoff, S.B., Castellanos, F.X., Petrides, M., Jefferies, E., Smallwood, J., 2016. Situating the default-mode network along a principal gradient of macroscale cortical organization. Proc. Natl. Acad. Sci. 113, 12574–12579. https://doi.org/10.1073/pnas.1608282113

Martin, A., 2016. GRAPES—Grounding representations in action, perception, and emotion systems: How object properties and categories are represented in the human brain. Psychon. Bull. Rev. 23, 979–990. https://doi.org/10.3758/s13423-015-0842-3

Martin, A., 2007. The Representation of Object Concepts in the Brain. Annu. Rev. Psychol. 58, 25–45. https://doi.org/10.1146/annurev.psych.57.102904.190143

Mesulam, M.M., 1998. From sensation to cognition. Brain 121, 1013–1052. https://doi.org/10.1093/brain/121.6.1013

Meteyard, L., Cuadrado, S.R., Bahrami, B., Vigliocco, G., 2012. Coming of age: A review of embodiment and the neuroscience of semantics. Cortex 48, 788–804. https://doi.org/10.1016/j.cortex.2010.11.002

Mion, M., Patterson, K., Acosta-Cabronero, J., Pengas, G., Izquierdo-Garcia, D., Hong, Y.T., Fryer, T.D., Williams, G.B., Hodges, J.R., Nestor, P.J., 2010. What the left and right anterior fusiform gyri tell us about semantic memory. Brain 133, 3256–3268. https://doi.org/10.1093/brain/awq272

Müller, V.I., Cieslik, E.C., Laird, A.R., Fox, P.T., Radua, J., Mataix-Cols, D., Tench, C.R., Yarkoni, T., Nichols, T.E., Turkeltaub, P.E., Wager, T.D., Eickhoff, S.B., 2018. Ten simple rules for neuroimaging meta-analysis. Neurosci. Biobehav. Rev. 84, 151–161. https://doi.org/10.1016/j.neubiorev.2017.11.012

Murray, S.O., Olshausen, B.A., Woods, D.L., 2002. Processing shape, motion, and three-dimensional shape-from-motion in the human cortex. J. Vis. 2, 508–516. https://doi.org/10.1167/2.7.303

Nichols, T., Brett, M., Andersson, J., Wager, T., Poline, J.-B., 2005. Valid conjunction inference with the minimum statistic. Neuroimage 25, 653–660. https://doi.org/10.1016/j.neuroimage.2004.12.005

Noonan, K.A., Jefferies, E., Visser, M., Lambon Ralph, M.A., 2013. Going beyond Inferior Prefrontal Involvement in Semantic Control: Evidence for the Additional Contribution of Dorsal Angular Gyrus and Posterior Middle Temporal Cortex. J. Cogn. Neurosci. 25, 1824–1850. https://doi.org/10.1162/jocn_a_00442

Norman, K.A., Polyn, S.M., Detre, G.J., Haxby, J. V., 2006. Beyond mind-reading: multi-voxel pattern analysis of fMRI data. Trends Cogn. Sci. 10, 424–430. https://doi.org/10.1016/j.tics.2006.07.005

Olson, I.R., Plotzker, A., Ezzyat, Y., 2007. The Enigmatic temporal pole: A review of findings on social and emotional processing. Brain 130, 1718–1731. https://doi.org/10.1093/brain/awm052

Page, M.J., Moher, D., Bossuyt, P.M., Boutron, I., Hoffmann, T.C., Mulrow, C.D., Shamseer, L., Tetzlaff, J.M., Akl, E.A., Brennan, S.E., Chou, R., Glanville, J., Grimshaw, J.M., Hróbjartsson, A., Lalu, M.M., Li, T., Loder, E.W., Mayo-Wilson, E., McDonald, S., McGuinness, L.A., Stewart, L.A., Thomas, J., Tricco, A.C., Welch, V.A., Whiting, P., McKenzie, J.E., 2021. PRISMA 2020 explanation and elaboration: updated guidance and exemplars for reporting systematic reviews. BMJ 372, n160. https://doi.org/10.1136/bmj.n160

Papitto, G., Friederici, A.D., Zaccarella, E., 2019. The topographical organization of motor processing: An ALE meta-analysis on six action domains and the relevance of Broca’s region. Neuroimage 116321. https://doi.org/10.1016/j.neuroimage.2019.116321

Patterson, K., Lambon Ralph, M.A., 2016. The hub-and-spoke hypothesis of semantic memory, in: Neurobiology of Language. Elsevier, Amsterdam, pp. 765–775.

Patterson, K., Nestor, P.J., Rogers, T.T., 2007. Where do you know what you know? The representation of semantic knowledge in the human brain. Nat. Rev. Neurosci. 8, 976– 987. https://doi.org/10.1038/nrn2277

Pitcher, D., Ungerleider, L.G., 2021. Evidence for a Third Visual Pathway Specialized for Social Perception. Trends Cogn. Sci. 25, 100–110. https://doi.org/10.1016/j.tics.2020.11.006

Pobric, G., Jefferies, E., Lambon Ralph, M.A., 2010a. Category-Specific versus Category-General Semantic Impairment Induced by Transcranial Magnetic Stimulation. Curr. Biol. 20, 964–968. https://doi.org/10.1016/j.cub.2010.03.070

Pobric, G., Jefferies, E., Lambon Ralph, M.A., 2010b. Amodal semantic representations depend on both anterior temporal lobes: Evidence from repetitive transcranial magnetic stimulation. Neuropsychologia 48, 1336–1342. https://doi.org/10.1016/j.neuropsychologia.2009.12.036

Postle, N., McMahon, K.L., Ashton, R., Meredith, M., de Zubicaray, G.I., 2008. Action word meaning representations in cytoarchitectonically defined primary and premotor cortices. Neuroimage 43, 634–644. https://doi.org/10.1016/j.neuroimage.2008.08.006

Price, C.J., Friston, K.J., 2002. Degeneracy and cognitive anatomy. Trends Cogn. Sci. 6, 416–421. https://doi.org/10.1016/S1364-6613(02)01976-9

Pylyshyn, Z.W., 1984. Computation and Cognition: Toward a Foundation for Cognitive Science, Psychological Review. The MIT Press. https://doi.org/10.1016/0004- 3702(89)90062-3

Raposo, A., Moss, H.E., Stamatakis, E.A., Tyler, L.K., 2009. Modulation of motor and premotor cortices by actions, action words and action sentences. Neuropsychologia 47, 388–396. https://doi.org/10.1016/j.neuropsychologia.2008.09.017

Reid, A.T., Bzdok, D., Genon, S., Langner, R., Müller, V.I., Eickhoff, C.R., Hoffstaedter, F., Cieslik, E.-C., Fox, P.T., Laird, A.R., Amunts, K., Caspers, S., Eickhoff, S.B., 2016. ANIMA: A data-sharing initiative for neuroimaging meta-analyses. Neuroimage 124, 1245–1253. https://doi.org/10.1016/j.neuroimage.2015.07.060

Reilly, J., Garcia, A., Binney, R.J., 2016a. Does the sound of a barking dog activate its corresponding visual form? An fMRI investigation of modality-specific semantic access. Brain Lang. 159, 45–59. https://doi.org/10.1016/j.bandl.2016.05.006

Reilly, J., Peelle, J.E., Garcia, A., Crutch, S.J., 2016b. Linking somatic and symbolic representation in semantic memory: the dynamic multilevel reactivation framework. Psychon. Bull. Rev. 23, 1002–1014. https://doi.org/10.3758/s13423-015-0824-5

Rice, G.E., Lambon Ralph, M.A., Hoffman, P., 2015. The Roles of Left Versus Right Anterior Temporal Lobes in Conceptual Knowledge: An ALE Meta-analysis of 97 Functional Neuroimaging Studies. Cereb. Cortex 25, 4374–4391. https://doi.org/10.1093/cercor/bhv024

Ross, L.A., Olson, I.R., 2010. Social cognition and the anterior temporal lobes. Neuroimage 49, 3452–3462. https://doi.org/10.1016/j.neuroimage.2009.11.012

Rottschy, C., Langner, R., Dogan, I., Reetz, K., Laird, A.R., Schulz, J.B., Fox, P.T., Eickhoff, S.B., 2012. Modelling neural correlates of working memory: A coordinate-based meta-analysis. Neuroimage 60, 830–846. https://doi.org/10.1016/j.neuroimage.2011.11.050

Saxe, R., Kanwisher, N., 2003. People thinking about thinking peopleThe role of the temporo-parietal junction in “theory of mind.” Neuroimage 19, 1835–1842. https://doi.org/10.1016/S1053-8119(03)00230-1

Saygin, A.P., 2012. Sensory and Motor Brain Areas Supporting Biological Motion Perception: Neuropsychological and Neuroimaging Studies, in: People Watching: Social, Perceptual, and Neurophysiological Studies of Body Perception. Oxford University Press, pp. 369–387. https://doi.org/10.1093/acprof:oso/9780195393705.003.0021

Sayres, R., Grill-Spector, K., 2008. Relating Retinotopic and Object-Selective Responses in Human Lateral Occipital Cortex. J. Neurophysiol. 100, 249–267. https://doi.org/10.1152/jn.01383.2007

Schacter, D.L., Benoit, R.G., Szpunar, K.K., 2017. Episodic future thinking: mechanisms and functions. Curr. Opin. Behav. Sci. 17, 41–50. https://doi.org/10.1016/j.cobeha.2017.06.002

Schurz, M., Radua, J., Tholen, M.G., Maliske, L., Margulies, D.S., Mars, R.B., Sallet, J., Kanske, P., 2020. Toward a hierarchical model of social cognition: A neuroimaging meta-analysis and integrative review of empathy and theory of mind. Psychol. Bull. https://doi.org/10.1037/bul0000303

Seghier, M.L., 2013. The angular gyrus: Multiple functions and multiple subdivisions. Neuroscientist 19, 43–61. https://doi.org/10.1177/1073858412440596

Simmons, W.K., Barsalou, L.W., 2003. The Similarity-in-Topography Principle: Reconciling Theories of Conceptual Deficits. Cogn. Neuropsychol. 20, 451–486. https://doi.org/10.1080/02643290342000032

Simmons, W.K., Ramjee, V., Beauchamp, M.S., McRae, K., Martin, A., Barsalou, L.W., 2007. A common neural substrate for perceiving and knowing about color. Neuropsychologia 45, 2802–2810. https://doi.org/10.1016/j.neuropsychologia.2007.05.002

Small, D.M., Bender, G., Veldhuizen, M.G., Rudenga, K., Nachtigal, D., Felsted, J., 2007. The role of the human orbitofrontal cortex in taste and flavor processing. Ann. N. Y. Acad. Sci. 1121, 136–51. https://doi.org/10.1196/annals.1401.002

Small, D.M., Prescott, J., 2005. Odor/taste integration and the perception of flavor. Exp. Brain Res. 166, 345–357. https://doi.org/10.1007/s00221-005-2376-9

Specht, K., Reul, J., 2003. Functional segregation of the temporal lobes into highly differentiated subsystems for auditory perception: An auditory rapid event-related fMRI-task. Neuroimage 20, 1944–1954. https://doi.org/10.1016/j.neuroimage.2003.07.034

Stanley, D.A., Rubin, N., 2005. Rapid detection of salient regions: Evidence from apparent motion. J. Vis. 5, 4. https://doi.org/10.1167/5.9.4

Thompson-Schill, S.L., 2003. Neuroimaging studies of semantic memory: inferring “how” from “where.” Neuropsychologia 41, 280–292. https://doi.org/10.1016/S0028-3932(02)00161-6

Tomasello, R., Garagnani, M., Wennekers, T., Pulvermüller, F., 2017. Brain connections of words, perceptions and actions: A neurobiological model of spatio-temporal semantic activation in the human cortex. Neuropsychologia 98, 111–129. https://doi.org/10.1016/j.neuropsychologia.2016.07.004

Tong, J., Binder, J.R., Humphries, C., Mazurchuk, S., Conant, L.L., Fernandino, L., 2022. A Distributed Network for Multimodal Experiential Representation of Concepts. J. Neurosci. 42, 7121–7130. https://doi.org/10.1523/JNEUROSCI.1243-21.2022

Trumpp, N.M., Kliese, D., Hoenig, K., Haarmeier, T., Kiefer, M., 2013. Losing the sound of concepts: Damage to auditory association cortex impairs the processing of sound-related concepts. Cortex 49, 474–486. https://doi.org/10.1016/j.cortex.2012.02.002

Turella, L., Lingnau, A., 2014. Neural correlates of grasping. Front. Hum. Neurosci. 8, 1–8. https://doi.org/10.3389/fnhum.2014.00686

Turkeltaub, P.E., Eickhoff, S.B., Laird, A.R., Fox, M., Wiener, M., Fox, P., 2012. Minimizing within-experiment and within-group effects in activation likelihood estimation meta-analyses. Hum. Brain Mapp. 33, 1–13. https://doi.org/10.1002/hbm.21186

Ulrich, M., Harpaintner, M., Trumpp, N.M., Berger, A., Kiefer, M., in press. Academic training increases grounding of scientific concepts in experiential brain systems. Cereb. Cortex. https://doi.org/10.1093/cercor/bhac449

Ungerleider, L.G., Desimone, R., 1986. Cortical connections of visual area MT in the macaque. J. Comp. Neurol. 248, 190–222. https://doi.org/10.1002/cne.902480204

van Dam, W.O., van Dijk, M., Bekkering, H., Rueschemeyer, S.A., 2012. Flexibility in embodied lexical-semantic representations. Hum. Brain Mapp. 33, 2322–2333. https://doi.org/10.1002/hbm.21365

van Elk, M., van Schie, H., Bekkering, H., 2014. Action semantics: A unifying conceptual framework for the selective use of multimodal and modality-specific object knowledge. Phys. Life Rev. 11, 220–250. https://doi.org/10.1016/j.plrev.2013.11.005

Van Overwalle, F., Baetens, K., 2009. Understanding others’ actions and goals by mirror and mentalizing systems: A meta-analysis. Neuroimage 48, 564–584. https://doi.org/10.1016/j.neuroimage.2009.06.009

Vigliocco, G., Kousta, S.T., Della Rosa, P.A., Vinson, D.P., Tettamanti, M., Devlin, J.T., Cappa, S.F., 2014. The neural representation of abstract words: The role of emotion. Cereb. Cortex 24, 1767–1777. https://doi.org/10.1093/cercor/bht025

Visser, M., Jefferies, E., Lambon Ralph, M.A., 2010. Semantic Processing in the Anterior Temporal Lobes: A Meta-analysis of the Functional Neuroimaging Literature. J. Cogn. Neurosci. 22, 1083–1094. https://doi.org/10.1162/jocn.2009.21309

Vukovic, N., Feurra, M., Shpektor, A., Myachykov, A., Shtyrov, Y., 2017. Primary motor cortex functionally contributes to language comprehension: An online rTMS study. Neuropsychologia 96, 222–229. https://doi.org/10.1016/j.neuropsychologia.2017.01.025

Wang, Y., Collins, J.A., Koski, J., Nugiel, T., Metoki, A., Olson, I.R., 2017. Dynamic neural architecture for social knowledge retrieval. Proc. Natl. Acad. Sci. 114, E3305–E3314. https://doi.org/10.1073/pnas.1621234114

Ward, N.S., Bestmann, S., Hartwigsen, G., Weiss, M.M., Christensen, L.O.D., Frackowiak, R.S.J., Rothwell, J.C., Siebner, H.R., 2010. Low-Frequency Transcranial Magnetic Stimulation over Left Dorsal Premotor Cortex Improves the Dynamic Control of Visuospatially Cued Actions. J. Neurosci. 30, 9216–9223. https://doi.org/10.1523/JNEUROSCI.4499-09.2010

Watson, C.E., Cardillo, E.R., Ianni, G.R., Chatterjee, A., 2013. Action Concepts in the Brain: An Activation Likelihood Estimation Meta-analysis. J. Cogn. Neurosci. 25, 1191–1205. https://doi.org/10.1162/jocn_a_00401

Weiner, K.S., Grill-Spector, K., 2011. Not one extrastriate body area: Using anatomical landmarks, hMT+, and visual field maps to parcellate limb-selective activations in human lateral occipitotemporal cortex. Neuroimage 56, 2183–2199. https://doi.org/10.1016/j.neuroimage.2011.03.041

Weiskopf, N., Hutton, C., Josephs, O., Deichmann, R., 2006. Optimal EPI parameters for reduction of susceptibility-induced BOLD sensitivity losses: A whole-brain analysis at 3 T and 1.5 T. Neuroimage 33, 493–504. https://doi.org/10.1016/j.neuroimage.2006.07.029

Wheeler, M.E., Petersen, S.E., Buckner, R.L., 2000. Memory’s echo: Vivid remembering reactivates sensory-specific cortex. Proc. Natl. Acad. Sci. 97, 11125–11129. https://doi.org/10.1073/pnas.97.20.11125

Whitney, C., Kirk, M., O’Sullivan, J., Lambon Ralph, M.A., Jefferies, E., 2012. Executive Semantic Processing Is Underpinned by a Large-scale Neural Network: Revealing the Contribution of Left Prefrontal, Posterior Temporal, and Parietal Cortex to Controlled Retrieval and Selection Using TMS. J. Cogn. Neurosci. 24, 133–147. https://doi.org/10.1162/jocn_a_00123

Willems, R.M., Casasanto, D., 2011. Flexibility in Embodied Language Understanding. Front. Psychol. 2, 116. https://doi.org/10.3389/fpsyg.2011.00116

Wilson-Mendenhall, C.D., Simmons, W.K., Martin, A., Barsalou, L.W., 2013. Contextual Processing of Abstract Concepts Reveals Neural Representations of Nonlinguistic Semantic Content. J. Cogn. Neurosci. 25, 920–935. https://doi.org/10.1162/jocn_a_00361

Xu, Y., 2007. The Role of the Superior Intraparietal Sulcus in Supporting Visual Short-Term Memory for Multifeature Objects. J. Neurosci. 27, 11676–11686. https://doi.org/10.1523/JNEUROSCI.3545-07.2007

Yarkoni, T., Poldrack, R.A., Van Essen, D.C., Wager, T.D., 2010. Cognitive neuroscience 2.0: building a cumulative science of human brain function. Trends Cogn. Sci. 14, 489– 496. https://doi.org/10.1016/j.tics.2010.08.004

Yee, E., Thompson-Schill, S.L., 2016. Putting concepts into context. Psychon. Bull. Rev. 23, 1015–1027. https://doi.org/10.3758/s13423-015-0948-7

Zatorre, R.J., Halpern, A.R., Perry, D.W., Meyer, E., Evans, A.C., 1996. Hearing in the Mind’s Ear: A PET Investigation of Musical Imagery and Perception. J. Cogn. Neurosci. 8, 29–46. https://doi.org/10.1162/jocn.1996.8.1.29

Zvyagintsev, M., Clemens, B., Chechko, N., Mathiak, K.A., Sack, A.T., Mathiak, K., 2013. Brain networks underlying mental imagery of auditory and visual information. Eur. J. Neurosci. 37, 1421–1434. https://doi.org/10.1111/ejn.12140

