## Supplementary Material for "Meta-analytic evidence for a novel hierarchical model of conceptual processing"

#### Supplementary Materials and Methods

**Table S1. Checklist for neuroimaging meta-analysis following the guidelines by Müller et al. (2018).**

| | |
|---|---|
| The research question is specifically defined | YES, and it includes the following contrasts:<br>• conceptual processing related to a certain perceptual-motor modality > high-level control condition (e.g. for action: action words > non-action words)<br>• conceptual processing related to a certain perceptual-motor modality > low-level baseline (e.g. for action: action words > fixation) |
| The literature search was systematic | YES, it included the following search queries in the following databases:<br>• **Review articles:** manual reference tracing<br>• **PubMed**: (((functional magnetic resonance imaging OR fMRI OR positron emission tomography OR PET OR neuroimaging)) AND (semantics OR semantic memory OR concepts OR conceptual OR knowledge)) AND (embodied OR embodiment OR grounded OR sensory OR motor OR perceptual)<br>• **PubMed**: ((((functional magnetic resonance imaging OR fMRI OR positron emission tomography OR PET OR neuroimaging)) AND (semantics OR semantic memory OR concepts OR conceptual OR knowledge)) AND (embodied OR embodiment OR grounded OR sensory OR perceptual) AND (tactile OR gustatory OR olfactory OR smell OR odor OR haptic OR taste))<br>• **PubMed**: (((((functional magnetic resonance imaging OR fMRI OR positron emission tomography OR PET OR neuroimaging)) AND (semantics OR semantic memory OR concepts OR conceptual OR knowledge)) AND (embodied OR embodiment OR grounded OR sensory OR perceptual) AND (tactile OR gustatory OR olfactory OR smell OR odor OR haptic OR taste OR somatosensory OR olfactive OR touch OR touching OR nose OR aroma OR aromatic OR flavour OR flavor OR perfume OR odoriferant OR emotion OR emotional OR emotive OR feeling OR valence OR affective OR social)))<br>• **Web of Science**: (ALL= neuroimaging conceptual) AND DOCUMENT TYPES: (Article)<br>• **Web of Science**: (ALL=(fmri OR functional magnetic resonance imaging OR positron emission tomography OR PET) AND ALL=(semantics OR semantic memory OR concepts OR conceptual OR knowledge) AND ALL=(embodied OR embodiment OR grounded OR sensory OR perceptual OR social) NOT ALL=(lesion OR damage OR autism))<br>• **Web of Science**: (ALL=(fmri OR functional magnetic resonance imaging OR positron emission tomography OR PET) AND TI=(color OR visual OR haptic OR tactile OR olfaction or smell OR motion) AND ALL=(conceptual OR semantic* OR semantic memory OR conceptual memory))<br>• **BrainMap (Sleuth)**: Normal mapping AND experiments - context: emotion<br>• **BrainMap (Sleuth)**: Experiments- Imaging modality-fMRI AND conditions- stimulus modality: gustatory<br>• **BrainMap (Sleuth)**: Normal mapping AND stimulus modality: olfactory<br>• **BrainMap (Sleuth)**: Experiments- Imaging modality-fMRI AND conditions- stimulus modality: auditory<br>• **BrainMap (Sleuth):** Normal mapping AND stimulus modality: tactile |

| | |
|---|---|
| Detailed inclusion and exclusion criteria are included | YES, and the criteria were:<br><br>**Inclusion**:<br>● **Participants**: healthy human adults; native speakers; right-handed<br>● **Neuroimaging methods**: fMRI, PET<br>● **Data analysis**: whole-brain, univariate (voxel-wise) activation analyses<br>● **Contrasts**: probing conceptual processing related to a certain perceptual–motor modality (action, color, emotion, motion, olfaction/gustation, shape, sound)<br><br>**Exclusion**:<br>● **Participants**: non-human subjects; children; patients with neurological or psychiatric disorders; non-native speakers; left-handed subjects<br>● **Data analyses**: search space was not whole-brain (e.g. region-of-interest analyses; partial brain coverage); analysis was not voxel-wise (e.g. multivariate); non-activation-based (e.g. functional connectivity) analyses<br>● **Contrasts**: non-conceptual contrasts; contrasts with potential perceptual difference in same modality as targeted conceptual modality; contrasts probing multiple modalities at once; contrasts probing non-included conceptual modalities/features |
| Sample overlap was taken into account | YES, using the following method:<br><br>Multiple contrasts from the same sample and study were pooled, i.e. treated as one single experiment (as recommended by Turkeltaub et al., 2012). |
| All experiments use the same search coverage | YES, the search coverage is the following:<br><br>Only coordinates from whole-brain analyses were included. ROI analyses and experiments with limited (non-whole-brain) coverage were excluded (as recommended by Müller et al., 2018). |
| Studies are converted to a common reference space | YES, using the following conversion(s):<br><br>Coordinates in Talairach space were transformed to MNI space using the Lancaster transform as implemented in *GingerALE* v3.0 (Matlab function *tal2icbm_spm.m*). |
| Data extraction have been conducted by two investigators (ideal case) or double checked by the same investigator (state how double-checking was performed) | YES, the following authors:<br><br>● PK, MB, JA, MK, GH checked inclusion/exclusion criteria<br>● MB, JA extracted coordinates<br>● MB, JA extracted other info: study metadata (e.g. number of subjects, task type, stimulus type, contrast type, reference space)<br>● PK, MB, JA double-checked the following data: correctness of coordinates (focusing on sign), study metadata |

| | |
|---|---|
| The paper includes a table with at least the references, basic study description (e.g., for fMRI task: stimuli), contrasts and basic sample descriptions (e.g., size, mean age and gender distribution, specific characteristics) of the included studies, source of information (e.g. contact with authors), reference space | YES, and also the following data:<br><br>See Tables S31-S37 for tables summarizing the relevant information for each included study. Studies have been organized according to modality, method used (fMRI or PET), sample size (number of subjects), task, stimulus modality, contrast(s) included in the analysis, coordinate space used in the original paper for reporting results (Talairach or MNI), number of foci, and contrast type (high- or low-level). |
| The study protocol was previously registered and all analyses planned beforehand, including the methods and parameters used for inference, correction for multiple testing, etc. | 1) Inclusion and exclusion criteria were defined beforehand based on explicit conceptual considerations (see above and in Methods section).<br><br>2) Any non-planned analyses are clearly labelled as supplementary or exploratory in the paper.<br><br>3) The meta-analysis used the recommended default methods and parameters of the *GingerALE* software (details elaborated in the Materials and Methods section; also see Müller et al., 2018). |
| The meta-analysis includes diagnostics | YES, the following:<br><br>● Robustness check: In addition to our primary analyses (which included all contrasts), we performed a supplementary analysis that excluded low-level contrasts (e.g. for action: action words > fixation; see Methods and Results sections for details). |

### Supplementary Results

#### Coordinate Tables

The following tables report significant clusters of the ALE analyses, i.e. brain regions consistently activated across neuroimaging studies. All analyses were thresholded at a voxel-wise $p < 0.001$ and a cluster-wise $p < 0.05$ FWE-corrected using Monte Carlo simulation (10,000 permutations). Coordinates are in MNI space. We report peaks more than 8 mm apart in clusters larger than 50 $mm^3$. Anatomical labels were determined using the SPM Anatomy toolbox (Version 2.2c; Eickhoff et al., 2005), the Harvard-Oxford atlas distributed with FSL (http://www.fmrib.ox.ac.uk/fsl/), and the human motor area template (http://lrnlab.org/; Mayka et al., 2006).

| | | | |
|---|---|---|---|
| L | left | OFC | orbitofrontal cortex |
| R | right | PCC | posterior cingulate cortex |
| a (prefix) | anterior | PFC | prefrontal cortex |
| p (prefix) | posterior | mPFC | medial PFC |
| d (prefix) | dorsal | dmPFC | dorsomedial PFC |
| v (prefix) | ventral | vmPFC | ventromedial PFC |
| A1 | primary auditory cortex | PMC | premotor cortex |
| ACC | anterior cingulate cortex | PMd | dorsal PMC |
| AG | angular gyrus | PMv | ventral PMC |
| FG | fusiform gyrus | PreCS | precentral sulcus |
| IFG | inferior frontal gyrus | S1 | primary somatosensory cortex |
| IPL | inferior parietal lobe | SMA | supplementary motor area |
| IPS | intraparietal sulcus | SMG | supramarginal gyrus |
| LOC | lateral occipital cortex | SPL | superior parietal lobe |
| LTO | lateral temporal-occipital junction | STG | superior temporal gyrus |
| M1 | primary motor cortex | STS | superior temporal sulcus |
| MCC | middle cingulate cortex | TP | temporal pole |
| MFG | middle frontal gyrus | TPJ | temporoparietal junction |
| MTG | middle temporal gyrus | V1/V2 | primary/secondary visual cortex |

**Table S2. ALE results for action-related conceptual processing.**

| Region | Cluster size (mm$^3$) | x | y | z | ALE | P | Z |
|---|---|---|---|---|---|---|---|
| L IFG / PMC | 16880 | | | | | | |
| L IFG (op) | | -48 | 6 | 24 | 0.056 | 2.26E-13 | 7.239 |
| L IFG (tri) | | -46 | 34 | 8 | 0.039 | 1.43E-08 | 5.550 |
| L IFG (tri) | | -48 | 20 | 0 | 0.033 | 4.17E-07 | 4.927 |
| L IFG (tri) | | -54 | 20 | 8 | 0.033 | 4.34E-07 | 4.919 |
| L IFG (op) | | -58 | 4 | 12 | 0.032 | 1.12E-06 | 4.730 |
| L IFG (tri) | | -46 | 20 | 24 | 0.031 | 1.49E-06 | 4.672 |
| L PMd | | -46 | 6 | 44 | 0.029 | 6.24E-06 | 4.369 |
| L PMd | | -46 | 2 | 44 | 0.028 | 9.11E-06 | 4.286 |
| L IFG (tri) | | -48 | 40 | -2 | 0.021 | 2.92E-04 | 3.439 |
| L LTO (pMTG/pSTS/pITG) | 13632 | | | | | | |
| L pMTG | | -54 | -58 | 0 | 0.057 | 5.96E-14 | 7.418 |
| L pMTG | | -46 | -62 | 12 | 0.031 | 1.96E-06 | 4.616 |
| L FG4 | | -38 | -44 | -20 | 0.030 | 2.70E-06 | 4.549 |
| L pSTS | | -56 | -38 | 0 | 0.029 | 4.26E-06 | 4.452 |
| L AG (PGp) | | -42 | -66 | 22 | 0.028 | 8.29E-06 | 4.307 |
| L aSMG / S1 | 10984 | | | | | | |
| L aSMG (PFt) | | -60 | -32 | 34 | 0.069 | 1.05E-17 | 8.488 |
| L S1 (area 2) | | -46 | -42 | 48 | 0.046 | 1.39E-10 | 6.311 |
| L/R (pre-)SMA | 6504 | | | | | | |
| L pre-SMA | | -4 | 12 | 52 | 0.037 | 3.57E-08 | 5.388 |
| L PMd | | -22 | 4 | 58 | 0.030 | 2.90E-06 | 4.533 |
| L pre-SMA | | -4 | 28 | 42 | 0.028 | 9.11E-06 | 4.286 |
| L PMd | | -28 | -4 | 54 | 0.027 | 1.21E-05 | 4.222 |
| L pre-SMA | | -10 | 4 | 62 | 0.027 | 1.85E-05 | 4.126 |
| L pre-SMA | | -12 | 16 | 60 | 0.022 | 2.47E-04 | 3.484 |
| L IPS (hIP3) | 1512 | -34 | -64 | 42 | 0.031 | 1.61E-06 | 4.656 |
| L OFC (Fo3) | 1008 | -30 | 36 | -14 | 0.037 | 6.10E-08 | 5.290 |

**Table S3. ALE results for real action execution.**

| Region | Cluster size (mm$^3$) | x | y | z | ALE | P | Z |
|---|---|---|---|---|---|---|---|
| L/R M1/S1, IFG/PMC, SMA, aSMG/IPS, pSTG/STS | 275560 | | | | | | |
| L SMA | | -2 | 0 | 56 | 0.379 | 0 | ∞ |
| L Thalamus (prefrontal) | | -12 | -18 | 4 | 0.305 | 0 | ∞ |
| L M1 (4a) | | -38 | -20 | 56 | 0.273 | 0 | ∞ |
| R Thalamus (premotor) | | 14 | -18 | 6 | 0.242 | 0 | ∞ |
| L Putamen | | -24 | -4 | 4 | 0.217 | 2.79E-39 | 13.060 |
| R PMv | | 52 | -6 | 36 | 0.207 | 4.97E-36 | 12.477 |
| L PMd | | -46 | -8 | 44 | 0.202 | 2.54E-34 | 12.160 |
| R Putamen | | 24 | 2 | 4 | 0.200 | 1.19E-33 | 12.033 |
| L M1 (4p) | | -52 | -8 | 32 | 0.192 | 2.80E-31 | 11.574 |

| | | | | | | | |
|---|---|---|---|---|---|---|---|
| L PMv | | -56 | -4 | 26 | 0.190 | 1.10E-30 | 11.456 |
| L/R Cerebellum | 33160 | | | | | | |
| R Cerebellum (VI) | | 20 | -56 | -22 | 0.232 | 4.80E-44 | 13.870 |
| L Cerebellum (VI) | | -22 | -60 | -24 | 0.166 | 5.60E-24 | 10.030 |
| L Cerebellum (VI) | | -26 | -58 | -24 | 0.164 | 2.17E-23 | 9.896 |
| L Cerebellum (VI) | | -14 | -62 | -20 | 0.162 | 4.25E-23 | 9.828 |
| R Cerebellum (V) | | 4 | -62 | -16 | 0.158 | 7.78E-22 | 9.531 |
| R FG2 | | 42 | -68 | -16 | 0.072 | 1.27E-04 | 3.659 |

**Table S4. Overlap between meta-analytic maps for action-related conceptual processing and real action execution.**

| Region | Cluster size (mm$^3$) | x | y | z | ALE | P | Z |
|---|---|---|---|---|---|---|---|
| L IFG / PMC | 8384 | | | | | | |
| L PMv | | -48 | 6 | 24 | 0.056 | 2.26E-13 | 7.239 |
| L IFG (tri) | | -46 | 20 | -2 | 0.032 | 7.57E-07 | 4.809 |
| L IFG (tri) | | -54 | 18 | 10 | 0.032 | 1.11E-06 | 4.732 |
| L IFG (op) | | -58 | 4 | 12 | 0.032 | 1.13E-06 | 4.730 |
| L PMd | | -46 | 6 | 44 | 0.029 | 6.24E-06 | 4.369 |
| L PMd | | -46 | 2 | 44 | 0.028 | 9.11E-06 | 4.286 |
| L IFG (tri) | | -44 | 18 | 24 | 0.027 | 1.22E-05 | 4.220 |
| L aSMG/IPS, S1 | 6872 | | | | | | |
| L aSMG (PFt) | | -58 | -30 | 34 | 0.067 | 5.50E-17 | 8.294 |
| L S1 (area 2) | | -46 | -42 | 48 | 0.046 | 1.39E-10 | 6.311 |
| L/R (pre-)SMA / PMd | 5176 | | | | | | |
| L pre-SMA | | -4 | 12 | 52 | 0.037 | 3.57E-08 | 5.388 |
| L PMd | | -22 | 2 | 58 | 0.028 | 8.86E-06 | 4.292 |
| L PMd | | -28 | -4 | 54 | 0.027 | 1.21E-05 | 4.222 |
| L pre-SMA | | -10 | 4 | 62 | 0.027 | 1.85E-05 | 4.126 |
| L PMd | | -16 | 2 | 54 | 0.026 | 2.07E-05 | 4.099 |
| L pre-SMA | | -4 | 26 | 42 | 0.026 | 2.86E-05 | 4.024 |
| L pSTS | 112 | | | | | | |
| L pSTS | | -56 | -36 | 2 | 0.024 | 6.54E-05 | 3.825 |
| L pSTS | | -54 | -40 | 4 | 0.022 | 1.83E-04 | 3.564 |
| L FG3 | 88 | -34 | -50 | -20 | 0.024 | 9.34E-05 | 3.736 |

**Table S5. Contrasts between conceptual processing related to action and all other modalities (inclusively masked by regions significant for action and no other modality).**

| Region | Cluster size (mm$^3$) | x | y | z | Action specificity (sign. contrasts out of 6) |
|---|---|---|---|---|---|
| L IFG / PMC | 11800 | | | | |
| L IFG (orb) | | -50 | 18 | -4 | 4 |
| L IFG (op) | | -46 | 16 | -2 | 3 |
| L IFG (tri) | | -54 | 22 | -6 | 2 |

| | | | | | |
|---|---|---|---|---|---|
| L IFG (orb) | | -48 | 20 | -8 | 1 |
| L aSMG / IPS | 9752 | | | | |
| L aSMG (PFop) | | -60 | -30 | 30 | 6 |
| L aSMG (PFop) | | -58 | -30 | 24 | 5 |
| L aSMG (PFcm) | | -54 | -42 | 22 | 4 |
| L aSMG (PFcm) | | -54 | -42 | 18 | 3 |
| L aSMG (PFcm) | | -58 | -42 | 16 | 2 |
| L pITG / LTO | 7424 | | | | |
| L pITG | | -56 | -58 | -8 | 6 |
| L pITG | | -58 | -54 | -8 | 5 |
| L pITG | | -56 | -62 | -12 | 4 |
| L pITG | | -50 | -54 | -16 | 3 |
| L FG2 | | -50 | -60 | -16 | 2 |
| L LTO | | -60 | -50 | 0 | 1 |
| L/R (pre-)SMA / PMC | 4872 | | | | |
| R pre-SMA | | 2 | 14 | 50 | 5 |
| L pre-SMA | | 0 | 10 | 48 | 4 |
| L PMd | | -16 | 0 | 50 | 3 |
| L pre-SMA | | -8 | 8 | 48 | 2 |
| L MCC | | -2 | 12 | 38 | 1 |
| L IPS / AG | 768 | | | | |
| L AG (PGa) | | -36 | -70 | 40 | 3 |
| L AG | | -38 | -64 | 38 | 2 |
| L IPS (hIP3) | | -38 | -60 | 38 | 1 |
| L FG4 | 512 | -40 | -48 | -24 | 1 |
| L IFG (orb) | 408 | | | | |
| L IFG (orb) | | -36 | 38 | -16 | 2 |
| L IFG (orb) | | -30 | 34 | -18 | 1 |
| L MTG | 360 | | | | |
| L MTG | | -56 | -36 | -8 | 2 |
| L MTG | | -60 | -38 | -6 | 1 |

**Table S6. ALE results for sound-related conceptual processing.**

| Region | Cluster size (mm³) | x | y | z | ALE | P | Z |
|---|---|---|---|---|---|---|---|
| L pSTS/pMTG | 1136 | -60 | -44 | 0 | 0.020 | 1.11E-06 | 4.733 |
| L SMG | 1008 | | | | | | |
| L SMG (PF) | | -60 | -42 | 28 | 0.018 | 7.11E-06 | 4.340 |
| L SMG (PFm) | | -60 | -52 | 26 | 0.017 | 1.03E-05 | 4.257 |
| R pSTS | 856 | 46 | -42 | 10 | 0.019 | 4.09E-06 | 4.460 |
| L dmPFC | 744 | | | | | | |
| L dmPFC | | -6 | 34 | 38 | 0.018 | 7.87E-06 | 4.318 |
| L dmPFC | | -2 | 28 | 44 | 0.014 | 1.48E-04 | 3.618 |

**Table S7. ALE results for real auditory perception.**

| Region | Cluster size ($mm^3$) | x | y | z | ALE | P | Z |
|---|---|---|---|---|---|---|---|
| L AC, STG/MTG, IFG/Insula, MFG/PreCS | 58360 | | | | | | |
| L STG | | -56 | -20 | 4 | 0.175 | 0 | ∞ |
| L IPL (PFcm) | | -46 | -34 | 12 | 0.113 | 1.41E-24 | 10.166 |
| L Insula | | -34 | 22 | 0 | 0.074 | 4.59E-13 | 7.142 |
| L IFG (tri) | | -46 | 20 | 22 | 0.057 | 9.25E-09 | 5.625 |
| L IFG (op) | | -44 | 8 | 28 | 0.056 | 1.61E-08 | 5.529 |
| L PMd | | -50 | -6 | 50 | 0.056 | 1.97E-08 | 5.494 |
| L PreCS | | -54 | 6 | 16 | 0.051 | 2.37E-07 | 5.037 |
| L IFG (tri) | | -38 | 42 | 4 | 0.049 | 4.82E-07 | 4.899 |
| L IFG (tri) | | -50 | 22 | 10 | 0.048 | 1.08E-06 | 4.738 |
| L IFG (orb) | | -46 | 28 | -4 | 0.045 | 3.01E-06 | 4.526 |
| L IFG (tri) | | -40 | 44 | 6 | 0.045 | 4.13E-06 | 4.458 |
| R AC, STG/MTC, IFG/Insula, MFG/PreCS | 53968 | | | | | | |
| R STG | | 58 | -16 | 2 | 0.145 | 1.39E-35 | 12.395 |
| R STG (TE 3) | | 62 | -26 | 8 | 0.072 | 1.59E-12 | 6.969 |
| R Insula | | 34 | 22 | -2 | 0.069 | 1.33E-11 | 6.664 |
| L Thalamus (prefrontal) | | -12 | -16 | 4 | 0.063 | 4.34E-10 | 6.132 |
| R IFG (op) | | 48 | 14 | 26 | 0.062 | 4.96E-10 | 6.111 |
| R PreCS | | 54 | 10 | 34 | 0.060 | 1.75E-09 | 5.906 |
| R PMd | | 54 | 0 | 46 | 0.054 | 4.32E-08 | 5.353 |
| R TP (TE 3) | | 54 | 6 | -16 | 0.053 | 6.10E-08 | 5.290 |
| R Putamen | | 24 | 4 | 2 | 0.052 | 9.81E-08 | 5.203 |
| R PMv | | 44 | 2 | 40 | 0.048 | 8.05E-07 | 4.797 |
| R IFG (op) | | 48 | 12 | 4 | 0.046 | 2.85E-06 | 4.537 |
| L/R pre-SMA | 9104 | -2 | 12 | 52 | 0.098 | 8.83E-20 | 9.027 |
| R IPS | 1864 | | | | | | |
| R IPS (hIP2) | | 42 | -44 | 44 | 0.053 | 9.23E-08 | 5.214 |
| R IPS (hIP3) | | 40 | -54 | 50 | 0.045 | 4.15E-06 | 4.457 |

**Table S8. Overlap between meta-analytic maps for sound-related conceptual processing and real auditory perception.**

| Region | Cluster size ($mm^3$) | x | y | z | ALE | P | Z |
|---|---|---|---|---|---|---|---|
| R pSTS | 400 | 46 | -42 | 10 | 0.019 | 4.09E-06 | 4.460 |
| L pSTS | 224 | -58 | -42 | 2 | 0.017 | 1.61E-05 | 4.157 |

**Table S9. Contrasts between conceptual processing related to sound and all other modalities (inclusively masked by regions significant for sound and no other modality).**

| Region | Cluster size ($mm^3$) | x | y | z | Sound specificity (sign. contrasts out of 6) |
|---|---|---|---|---|---|
| R pSTS | 856 | 46 | -44 | 6 | 6 |

| | | | | | |
|---|---|---|---|---|---|
| L pMTG | 568 | -58 | -42 | -10 | 5 |
| L dmPFC | 528 | | | | |
| L dmPFC | | -6 | 32 | 34 | 5 |
| L dmPFC | | -2 | 28 | 32 | 4 |
| L dmPFC | | 0 | 26 | 40 | 2 |
| L pSMG | 432 | | | | |
| L pSMG (PFm) | | -62 | -48 | 22 | 5 |

**Table S10. ALE results for shape-related conceptual processing.**

| Region | Cluster size (mm$^3$) | x | y | z | ALE | P | Z |
|---|---|---|---|---|---|---|---|
| L aFG / Hippocampus | 1368 | | | | | | |
| L aFG (FG3) | | -40 | -34 | -22 | 0.022 | 1.43E-07 | 5.133 |
| L Hippocampus (CA3) | | -32 | -26 | -12 | 0.016 | 1.23E-05 | 4.219 |
| L LOC (hOc4la) | 1160 | -52 | -66 | -4 | 0.020 | 5.90E-07 | 4.859 |
| L PreCS | 832 | -44 | 2 | 32 | 0.016 | 1.31E-05 | 4.205 |

**Table S11. ALE results for real visual shape perception.**

| Region | Cluster size (mm$^3$) | x | y | z | ALE | P | Z |
|---|---|---|---|---|---|---|---|
| Early visual cortex (V1/V2/V3/V4), R LOC (hOc4la/p), IPS/SPL, FG | 46744 | | | | | | |
| R SPL (7A) | | 26 | -64 | 52 | 0.091 | 1.44E-19 | 8.973 |
| R FG4 | | 42 | -54 | -20 | 0.090 | 3.71E-19 | 8.868 |
| R LOC (hOc4la) | | 42 | -76 | -8 | 0.086 | 4.84E-18 | 8.578 |
| R FG2 | | 46 | -64 | -12 | 0.085 | 9.94E-18 | 8.494 |
| R LOC (hOc4lp) | | 38 | -86 | 0 | 0.066 | 2.68E-12 | 6.896 |
| R IPS (hIP3) | | 32 | -52 | 52 | 0.065 | 7.37E-12 | 6.750 |
| R LOC | | 32 | -76 | 32 | 0.061 | 9.65E-11 | 6.367 |
| R IPS (hIP3) | | 40 | -42 | 46 | 0.056 | 1.18E-09 | 5.970 |
| L V1 (hOc1) | | -12 | -94 | -2 | 0.043 | 1.42E-06 | 4.682 |
| R V1 (hOc1) | | 8 | -86 | 4 | 0.041 | 3.90E-06 | 4.471 |
| R LOC | | 34 | -82 | 18 | 0.034 | 1.16E-04 | 3.680 |
| L LOC (hOc4la), FG | 20632 | | | | | | |
| L LOC (hOc4la) | | -40 | -80 | -8 | 0.084 | 3.00E-17 | 8.365 |
| L FG4 | | -38 | -56 | -18 | 0.078 | 1.53E-15 | 7.886 |
| L LOC (hOc4la) | | -46 | -68 | -10 | 0.074 | 2.34E-14 | 7.540 |
| L Cerebellum (VIIa crus I) | | -40 | -68 | -28 | 0.030 | 5.79E-04 | 3.249 |
| L IPS/SPL, S1 | 18128 | | | | | | |
| L SPL (7A) | | -24 | -66 | 48 | 0.077 | 3.82E-15 | 7.775 |
| L S1 (area 2) | | -40 | -34 | 44 | 0.067 | 2.49E-12 | 6.906 |
| L SPL (7A) | | -16 | -60 | 58 | 0.059 | 3.16E-10 | 6.182 |
| L IPS (hIP1) | | -36 | -48 | 46 | 0.056 | 1.33E-09 | 5.952 |
| L SPL (7PC) | | -36 | -50 | 60 | 0.046 | 4.02E-07 | 4.935 |
| L LOC | | -30 | -86 | 26 | 0.039 | 8.79E-06 | 4.293 |

| | | | | | | | |
|---|---|---|---|---|---|---|---|
| L PreCS/MFG, IFG/Insula | 14808 | | | | | | |
| L PreCS | | -48 | 8 | 30 | 0.072 | 5.92E-14 | 7.419 |
| L MFG | | -26 | -2 | 56 | 0.062 | 4.18E-11 | 6.494 |
| L Insula | | -34 | 22 | 2 | 0.060 | 1.31E-10 | 6.320 |
| L IFG (tri) | | -44 | 22 | 18 | 0.048 | 1.48E-07 | 5.127 |
| L MFG | | -40 | 4 | 44 | 0.045 | 4.81E-07 | 4.899 |
| L IFG (tri) | | -44 | 20 | -2 | 0.030 | 6.54E-04 | 3.214 |
| L/R pre-SMA | 7864 | 4 | 12 | 52 | 0.085 | 1.71E-17 | 8.431 |
| R PMd | 4720 | 30 | -4 | 54 | 0.062 | 4.66E-11 | 6.478 |
| R IFG/PMC | 4560 | | | | | | |
| R IFG (op) | | 50 | 8 | 28 | 0.071 | 2.02E-13 | 7.254 |
| R PMd | | 46 | 8 | 42 | 0.031 | 3.49E-04 | 3.391 |
| R IFG/Insula | 4280 | | | | | | |
| R Insula | | 36 | 24 | -4 | 0.050 | 4.93E-08 | 5.329 |
| R IFG (tri) | | 44 | 30 | 8 | 0.041 | 3.74E-06 | 4.479 |
| L Hippocampus/Amygdala | 1768 | -20 | -8 | -16 | 0.059 | 2.29E-10 | 6.233 |
| R Hippocampus/Amygdala | 1600 | 22 | -6 | -16 | 0.055 | 2.12E-09 | 5.875 |

**Table S12. Overlap between meta-analytic maps for shape-related conceptual processing and real visual shape perception.**

| **Region** | **Cluster size ($mm^3$)** | **x** | **y** | **z** | **ALE** | **P** | **Z** |
|---|---|---|---|---|---|---|---|
| L LOC (hOc4la) | 880 | -52 | -66 | -4 | 0.020 | 5.90E-07 | 4.859 |
| L PreCS | 728 | -44 | 2 | 32 | 0.016 | 1.31E-05 | 4.205 |

**Table S13. Contrasts between conceptual processing related to visual shape and all other modalities (inclusively masked by regions significant for shape and no other modality).**

| **Region** | **Cluster size ($mm^3$)** | **x** | **y** | **z** | **Shape specificity (sign. contrasts out of 6)** |
|---|---|---|---|---|---|
| L aFG / Hippocampus | 1232 | | | | |
| L aFG (FG4) | | -40 | -34 | -26 | 4 |
| L Hippocampus (DG) | | -30 | -30 | -14 | 3 |
| L LOC (hOc4la) | 120 | | | | |
| L LOC (hOc4la) | | -52 | -70 | -8 | 5 |
| L LOC (hOc4la) | | -50 | -66 | -12 | 3 |
| L PreCS | 80 | -44 | 0 | 32 | 5 |

**Table S14. ALE results for motion-related conceptual processing.**

| **Region** | **Cluster size ($mm^3$)** | **x** | **y** | **z** | **ALE** | **P** | **Z** |
|---|---|---|---|---|---|---|---|
| L pSTS/pMTG, aSMG | 8000 | | | | | | |
| L pSTS | | -54 | -50 | 6 | 0.027 | 2.02E-09 | 5.883 |
| L pMTG | | -56 | -58 | 4 | 0.025 | 1.44E-08 | 5.548 |
| L pSTS | | -62 | -52 | 10 | 0.024 | 4.16E-08 | 5.360 |
| L pSTS | | -54 | -34 | 2 | 0.020 | 6.63E-07 | 4.836 |

| | | | | | | | |
|---|---|---|---|---|---|---|---|
| L aSMG (PFcm) | | -58 | -42 | 24 | 0.018 | 3.75E-06 | 4.479 |
| L aSMG (PFt) | | -60 | -34 | 36 | 0.016 | 1.27E-05 | 4.212 |
| L AG (PGp) | | -54 | -64 | 22 | 0.015 | 3.26E-05 | 3.993 |
| L dFG (FG3) | 888 | -30 | -40 | -18 | 0.022 | 1.50E-07 | 5.123 |
| L IFG | 800 | | | | | | |
| L IFG (tri) | | -52 | 24 | 20 | 0.017 | 9.88E-06 | 4.268 |
| L IFG (op) | | -52 | 16 | 22 | 0.012 | 2.32E-04 | 3.501 |

**Table S15. ALE results for real visual motion perception.**

| Region | Cluster size (mm$^3$) | x | y | z | ALE | P | Z |
|---|---|---|---|---|---|---|---|
| L/R SPL/IPS, LOC | 51616 | | | | | | |
| L SPL (7PC) | | -30 | -54 | 54 | 0.108 | 2.24E-29 | 11.192 |
| R SPL (7A) | | 22 | -62 | 56 | 0.084 | 3.37E-20 | 9.132 |
| R LOC (hOc4la / V5) | | 46 | -70 | 0 | 0.083 | 5.86E-20 | 9.072 |
| L SPL (7A) | | -20 | -64 | 58 | 0.070 | 9.22E-16 | 7.956 |
| L LOC | | -24 | -84 | 24 | 0.058 | 6.76E-12 | 6.763 |
| R LOC | | 30 | -70 | 40 | 0.050 | 1.22E-09 | 5.966 |
| R LOC | | 26 | -74 | 38 | 0.047 | 5.59E-09 | 5.712 |
| R IPS (hIP2) | | 42 | -44 | 46 | 0.046 | 1.36E-08 | 5.559 |
| L SPL (7A) | | -4 | -60 | 56 | 0.038 | 1.56E-06 | 4.662 |
| R LOC | | 28 | -78 | 28 | 0.037 | 2.30E-06 | 4.582 |
| L/R PMC, (pre-)SMA | 31712 | | | | | | |
| L pre-SMA | | 0 | 4 | 58 | 0.101 | 2.21E-26 | 10.563 |
| R PMd | | 42 | -4 | 50 | 0.098 | 2.55E-25 | 10.331 |
| R PMd | | 26 | 0 | 56 | 0.077 | 5.54E-18 | 8.562 |
| R PMv | | 50 | 0 | 36 | 0.048 | 3.40E-09 | 5.796 |
| R MCC | | 6 | 22 | 32 | 0.047 | 7.56E-09 | 5.660 |
| L PMC | 14120 | | | | | | |
| L PMd | | -32 | -4 | 52 | 0.091 | 5.65E-23 | 9.800 |
| L PMd | | -42 | -2 | 46 | 0.070 | 9.29E-16 | 7.956 |
| L PMv | | -50 | 2 | 40 | 0.061 | 9.10E-13 | 7.048 |
| L Thalamus/Putamen, Insula | 8248 | | | | | | |
| L Thalamus (prefrontal) | | -12 | -16 | 6 | 0.056 | 1.75E-11 | 6.624 |
| L Insula | | -32 | 18 | 4 | 0.043 | 7.70E-08 | 5.248 |
| L Putamen | | -22 | 12 | 2 | 0.038 | 1.60E-06 | 4.658 |
| L Putamen | | -24 | -2 | 8 | 0.037 | 2.50E-06 | 4.565 |
| L Insula | | -38 | 16 | -6 | 0.033 | 2.53E-05 | 4.053 |
| R Thalamus/Putamen, Insula | 5904 | | | | | | |
| R Putamen | | 24 | 10 | 4 | 0.049 | 2.35E-09 | 5.857 |
| R Thalamus (parietal) | | 16 | -16 | 12 | 0.039 | 1.07E-06 | 4.740 |
| R Thalamus (prefrontal) | | 8 | -22 | 8 | 0.034 | 1.06E-05 | 4.253 |
| R Insula | | 36 | 22 | 6 | 0.034 | 1.32E-05 | 4.203 |
| R Thalamus (prefrontal) | | 12 | -14 | -2 | 0.030 | 9.26E-05 | 3.738 |

| | | | | | | | |
|---|---|---|---|---|---|---|---|
| L LOC (hOc4la / V5) | 4640 | -44 | -76 | 4 | 0.064 | 1.08E-13 | 7.338 |
| L V1/V2 | 3640 | | | | | | |
| L V1 (hOc1) | | -12 | -88 | -2 | 0.049 | 2.03E-09 | 5.882 |
| L V2 (hOc2) | | -4 | -84 | -8 | 0.031 | 7.40E-05 | 3.794 |
| L Cerebellum (VI) | | -10 | -76 | -16 | 0.029 | 1.90E-04 | 3.553 |
| R V1 | 2024 | 8 | -88 | 2 | 0.047 | 9.66E-09 | 5.618 |

**Table S16. Overlap between meta-analytic maps for motion-related conceptual processing and real action execution.**

| Region | Cluster size ($mm^3$) | x | y | z | ALE | P | Z |
|---|---|---|---|---|---|---|---|
| L pSTS | 464 | -54 | -34 | 2 | 0.020 | 6.30E-07 | 4.836 |
| L aSMG | 232 | | | | | | |
| L aSMG (PFt) | | -58 | -34 | 36 | 0.015 | 2.91E-05 | 4.020 |
| L aSMG (PFcm) | | -58 | -38 | 26 | 0.013 | 1.21E-04 | 3.670 |
| L aSMG (PF/PFcm) | | -58 | -40 | 20 | 0.011 | 7.45E-04 | 3.177 |

**Table S17. Contrasts between conceptual processing related to motion and all other modalities (inclusively masked by regions significant for motion and no other modality).**

| Region | Cluster size ($mm^3$) | x | y | z | Motion specificity (sign. contrasts out of 6) |
|---|---|---|---|---|---|
| L pSTS / pMTG | 2544 | | | | |
| L pSTS | | -62 | -56 | 8 | 6 |
| L pSTS | | -58 | -52 | 8 | 5 |
| L pMTG | | -64 | -52 | 4 | 4 |
| L pSTS | | -50 | -56 | 16 | 3 |
| L dFG | 808 | | | | |
| L dFG (FG3) | | -32 | -38 | -16 | 3 |
| L dFG (FG3) | | -30 | -38 | -24 | 2 |
| L dFG (FG3) | | -28 | -44 | -18 | 1 |
| L STS | 544 | | | | |
| L STS | | -58 | -32 | -2 | 2 |
| L STS | | -56 | -28 | 0 | 1 |
| L IFG (tri) | 184 | | | | |
| L IFG (tri) | | -54 | 28 | 18 | 2 |
| L IFG (tri) | | -54 | 22 | 20 | 1 |
| L pMTG | 152 | -54 | -64 | 18 | 3 |

**Table S18. ALE results for color-related conceptual processing.**

| Region | Cluster size ($mm^3$) | x | y | z | ALE | P | Z |
|---|---|---|---|---|---|---|---|
| L IPS (hIP3) | 1048 | -36 | -62 | 46 | 0.018 | 6.51E-07 | 4.839 |
| L vFG (FG4) | 896 | -42 | -50 | -12 | 0.016 | 3.73E-06 | 4.480 |

**Table S19. ALE results for real color perception.**

| Region | Cluster size (mm$^3$) | x | y | z | ALE | P | Z |
|---|---|---|---|---|---|---|---|
| L IPS | 3816 | | | | | | |
| L IPS (hIP3) | | -34 | -56 | 50 | 0.047 | 6.51E-13 | 7.094 |
| L IPS (hIP3) | | -28 | -64 | 42 | 0.030 | 1.87E-07 | 5.082 |
| R IPS | 3360 | | | | | | |
| R IPS (hIP3) | | 30 | -56 | 52 | 0.030 | 1.71E-07 | 5.099 |
| R IPS (hIP2) | | 42 | -40 | 44 | 0.029 | 2.92E-07 | 4.997 |
| R IPS (hIP3) | | 32 | -58 | 46 | 0.029 | 3.61E-07 | 4.955 |
| R IPS (hIP2) | | 44 | -40 | 52 | 0.022 | 2.39E-05 | 4.066 |
| L Insula | 1808 | -32 | 22 | 4 | 0.028 | 9.15E-07 | 4.771 |
| R Insula | 1536 | 34 | 22 | 0 | 0.028 | 6.84E-07 | 4.830 |
| L pre-SMA | 1416 | | | | | | |
| L pre-SMA | | 0 | 16 | 48 | 0.028 | 7.76E-07 | 4.805 |
| L pre-SMA | | -6 | 12 | 58 | 0.019 | 2.33E-04 | 3.499 |
| L SPL (7A) | | -18 | -68 | 60 | 0.039 | 2.23E-10 | 6.237 |
| R LOC (hOc4lp) | 1168 | 34 | -88 | 0 | 0.028 | 6.36E-07 | 4.844 |

**Table S20. Overlap between meta-analytic maps for color-related conceptual processing and real color perception.**

| Region | Cluster size (mm$^3$) | x | y | z | ALE | P | Z |
|---|---|---|---|---|---|---|---|
| L IPS (hIP3) | 648 | -36 | -60 | 44 | 0.018 | 6.68E-07 | 4.834 |

**Table S21. Contrasts between conceptual processing related to color and all other modalities (inclusively masked by regions significant for color and no other modality).**

| Region | Cluster size (mm$^3$) | x | y | z | Color specificity (sign. contrasts out of 6) |
|---|---|---|---|---|---|
| L vFG (FG4) | 744 | -44 | -52 | -18 | 6 |
| L IPS | 560 | | | | |
| L IPS (hIP3) | | -34 | -60 | 46 | 5 |
| L IPS (hIP1) | | -40 | -60 | 46 | 4 |

**Table S22. ALE results for olfaction-gustation related conceptual processing.**

| Region | Cluster size (mm$^3$) | x | y | z | ALE | P | Z |
|---|---|---|---|---|---|---|---|
| L OFC | 1744 | | | | | | |
| L OFC (Fo3) | | -24 | 36 | -14 | 0.019 | 6.98E-07 | 4.826 |
| L OFC (Fo3) | | -26 | 40 | -16 | 0.018 | 9.85E-07 | 4.756 |

**Table S23. ALE results for real olfactory-gustatory perception.**

| Region | Cluster size (mm$^3$) | x | y | z | ALE | P | Z |
|---|---|---|---|---|---|---|---|
| L/R OFC, Amygdala / Hippocampus, Thalamus, Insula | 77536 | | | | | | |

| | | | | | | | |
|---|---|---|---|---|---|---|---|
| L Amygdala (LB) | | -20 | -4 | -20 | 0.096 | 2.37E-25 | 10.338 |
| R Amygdala (LB) | | 24 | 0 | -18 | 0.092 | 5.36E-24 | 10.035 |
| R Thalamus (prefrontal) | | 8 | -8 | -2 | 0.067 | 1.91E-15 | 7.862 |
| L Basal Forebrain (Ch 1-3) | | -6 | 2 | -14 | 0.059 | 6.40E-13 | 7.097 |
| L Insula | | -32 | 18 | 2 | 0.055 | 9.39E-12 | 6.715 |
| R Insula | | 34 | 24 | -4 | 0.053 | 3.39E-11 | 6.525 |
| R Thalamus | | 8 | 2 | -2 | 0.052 | 6.35E-11 | 6.431 |
| R Insula (Id1) | | 38 | -10 | -12 | 0.049 | 6.79E-10 | 6.060 |
| R Insula | | 36 | 16 | -8 | 0.046 | 2.95E-09 | 5.820 |
| L Insula | | -34 | 2 | 0 | 0.045 | 5.75E-09 | 5.707 |
| R OFC (Fo3) | | 28 | 32 | -10 | 0.044 | 8.80E-09 | 5.634 |
| L/R ACC | 3624 | | | | | | |
| R ACC | | 2 | 44 | -2 | 0.049 | 6.84E-10 | 6.059 |
| L ACC (area 33) | | -2 | 32 | 0 | 0.036 | 1.07E-06 | 4.740 |

**Table S24. Overlap between meta-analytic maps for olfaction-gustation related conceptual processing and real olfactory-gustatory perception.**

| Region | Cluster size ($mm^3$) | x | y | z | ALE | P | Z |
|---|---|---|---|---|---|---|---|
| L OFC | 744 | | | | | | |
| L OFC (Fo3) | | -26 | 40 | -16 | 0.018 | 9.85E-07 | 4.756 |
| L OFC (Fo3) | | -24 | 34 | -16 | 0.017 | 1.82E-06 | 4.631 |

**Table S25. Contrasts between conceptual processing related to olfaction-gustation and all other modalities (inclusively masked by regions significant for olfaction-gustation and no other modality).**

| Region | Cluster size ($mm^3$) | x | y | z | Olfaction-gustation specificity (sign. contrasts out of 6) |
|---|---|---|---|---|---|
| L OFC | 1528 | | | | |
| L OFC (Fo3) | | -18 | 38 | -14 | 6 |
| L OFC (Fo3) | | -22 | 40 | -18 | 5 |
| L OFC (Fo3) | | -22 | 34 | -20 | 4 |
| L OFC (Fo3) | | -22 | 30 | -18 | 3 |

**Table S26. ALE results for emotion-related conceptual processing.**

| Region | Cluster size ($mm^3$) | x | y | z | ALE | P | Z |
|---|---|---|---|---|---|---|---|
| L/R dmPFC | 3576 | | | | | | |
| R dmPFC | | 6 | 54 | 22 | 0.027 | 1.13E-06 | 4.728 |
| R dmPFC (Fp2) | | 4 | 56 | 6 | 0.025 | 5.26E-06 | 4.406 |
| L dmPFC | | -6 | 62 | 36 | 0.020 | 9.46E-05 | 3.733 |
| L Amygdala / Hippocampus | 3368 | | | | | | |
| L Amygdala (LB) | | -22 | -4 | -18 | 0.048 | 7.05E-13 | 7.083 |
| L Hippocampus (CA3) | | -26 | -18 | -16 | 0.018 | 4.51E-04 | 3.319 |

| | | | | | | | |
|---|---|---|---|---|---|---|---|
| L AG | 2168 | | | | | | |
| L AG (PGp) | | -48 | -64 | 26 | 0.031 | 1.54E-07 | 5.119 |
| L AG (PGa) | | -54 | -58 | 22 | 0.024 | 1.02E-05 | 4.261 |
| R Amygdala (LB) | 2064 | 24 | -2 | -24 | 0.034 | 1.71E-08 | 5.519 |
| L/R vmPFC | 1952 | | | | | | |
| L vmPFC (Fp2) | | -2 | 48 | -18 | 0.021 | 4.94E-05 | 3.893 |
| R vmPFC (Fp2) | | 2 | 48 | -12 | 0.021 | 6.30E-05 | 3.834 |
| L vmPFC (Fp2) | | 2 | 56 | -18 | 0.020 | 1.29E-04 | 3.655 |

**Table S27. ALE results for real emotion perception.**

| Region | Cluster size ($mm^3$) | x | y | z | ALE | P | Z |
|---|---|---|---|---|---|---|---|
| L/R Amygdala / Hippocampus, Thalamus, IFG / Insula | 159328 | | | | | | |
| L Amygdala (CM) | | -20 | -4 | -16 | 0.336 | 0 | ∞ |
| L Pallidum | | -12 | 8 | -6 | 0.300 | 0 | ∞ |
| R Amygdala | | 24 | -4 | -16 | 0.283 | 0 | ∞ |
| R Caudate | | 12 | 10 | -4 | 0.269 | 3.40E-44 | 13.895 |
| L Insula | | -32 | 22 | -2 | 0.264 | 1.44E-42 | 13.624 |
| R Insula | | 36 | 22 | -2 | 0.246 | 2.84E-37 | 12.703 |
| R Thalamus (prefrontal) | | 6 | -14 | 8 | 0.181 | 2.13E-20 | 9.181 |
| L Thalamus (prefrontal) | | -10 | -16 | 8 | 0.178 | 7.79E-20 | 9.041 |
| L Brainstem | | -4 | -26 | -6 | 0.156 | 3.37E-15 | 7.791 |
| R IFG (op) | | 46 | 8 | 26 | 0.150 | 5.85E-14 | 7.420 |
| L PreCS | | -44 | 6 | 30 | 0.143 | 1.08E-12 | 7.024 |
| L Cingulate, mPFC | 60464 | | | | | | |
| L dmPFC | | -2 | 12 | 54 | 0.201 | 4.48E-25 | 10.277 |
| R MCC | | 4 | 22 | 38 | 0.189 | 2.45E-22 | 9.650 |
| R dmPFC | | 4 | 20 | 42 | 0.188 | 3.83E-22 | 9.604 |
| L vmPFC (Fp2) | | 0 | 54 | -8 | 0.180 | 3.27E-20 | 9.135 |
| L dmPFC (Fp2) | | -2 | 56 | 12 | 0.161 | 3.56E-16 | 8.077 |
| R PCC | | 2 | -26 | 34 | 0.159 | 7.31E-16 | 7.973 |
| L dmPFC (Fp2) | | -2 | 58 | 20 | 0.159 | 7.87E-16 | 7.973 |
| L ACC | | 0 | 42 | 6 | 0.146 | 3.21E-13 | 7.191 |
| L vmPFC (s32) | | -4 | 40 | -12 | 0.140 | 3.16E-12 | 6.872 |
| L PCC | | -4 | -52 | 26 | 0.133 | 5.79E-11 | 6.445 |
| L MCC | | 0 | -4 | 44 | 0.099 | 1.05E-05 | 4.255 |
| R LOC / FG | 10536 | | | | | | |
| R LOC (hOc4la) | | 46 | -74 | -4 | 0.138 | 8.26E-12 | 6.734 |
| R FG4 | | 44 | -54 | -18 | 0.109 | 4.51E-07 | 4.912 |
| R FG4 | | 44 | -50 | -20 | 0.108 | 5.62E-07 | 4.869 |
| R FG1 | | 32 | -62 | -18 | 0.107 | 7.42E-07 | 4.814 |
| R FG2 | | 44 | -62 | -12 | 0.103 | 3.17E-06 | 4.514 |
| R V3v (hOc3v) | | 24 | -90 | -8 | 0.092 | 7.63E-05 | 3.787 |

| | | | | | | | |
|---|---|---|---|---|---|---|---|
| R V3v (hOc3v) | | 20 | -82 | -12 | 0.090 | 1.28E-04 | 3.657 |
| L AG, LOC / FG | 9920 | | | | | | |
| L FG4 | | -42 | -60 | -16 | 0.157 | 1.77E-15 | 7.870 |
| L LOC (hOc4la) | | -42 | -76 | -2 | 0.116 | 3.69E-08 | 5.382 |
| L AG (PGa) | | -48 | -58 | 28 | 0.111 | 2.12E-07 | 5.058 |
| L LOC | | -52 | -66 | 6 | 0.105 | 1.69E-06 | 4.646 |
| L AG (PGa) | | -54 | -58 | 16 | 0.087 | 2.86E-04 | 3.445 |

**Table S28. Overlap between meta-analytic maps for emotion-related conceptual processing and real emotion perception.**

| Region | Cluster size ($mm^3$) | x | y | z | ALE | P | Z |
|---|---|---|---|---|---|---|---|
| L Amygdala / Hippocampus | 3360 | | | | | | |
| L Amygdala (LB) | | -22 | -4 | -18 | 0.048 | 3.09E-12 | 6.875 |
| L Hippocampus (CA2) | | -32 | -16 | -16 | 0.022 | 6.51E-05 | 3.826 |
| L/R dmPFC | 3168 | | | | | | |
| R dmPFC (Fp2) | | 6 | 54 | 6 | 0.029 | 1.15E-06 | 4.725 |
| R dmPFC | | 6 | 54 | 22 | 0.028 | 2.20E-06 | 4.592 |
| L dmPFC | | -8 | 58 | 12 | 0.024 | 1.89E-05 | 4.121 |
| L dmPFC (Fp2) | | -4 | 60 | 4 | 0.022 | 6.40E-05 | 3.830 |
| R Amygdala (LB) | 1888 | 24 | -2 | -24 | 0.034 | 4.47E-08 | 5.347 |
| L/R vmPFC | 1184 | | | | | | |
| L vmPFC (Fp2) | | -2 | 48 | -18 | 0.021 | 9.20E-05 | 3.740 |
| R vmPFC (Fp2) | | 2 | 48 | -12 | 0.021 | 1.16E-04 | 3.682 |
| R vmPFC (Fp2) | | 4 | 50 | -10 | 0.021 | 1.27E-04 | 3.657 |
| L vmPFC (Fp2) | | 2 | 56 | -18 | 0.020 | 2.28E-04 | 3.506 |
| R vmPFC | | 8 | 46 | -6 | 0.019 | 2.83E-04 | 3.447 |
| L AG | 568 | | | | | | |
| L AG (PGp) | | -48 | -64 | 24 | 0.029 | 9.88E-07 | 4.756 |
| L AG (PGa) | | -54 | -58 | 20 | 0.024 | 2.24E-05 | 4.081 |

**Table S29. Contrasts between conceptual processing related to emotion and all other modalities (inclusively masked by regions significant for emotion and no other modality).**

| Region | Cluster size ($mm^3$) | x | y | z | Emotion specificity (sign. contrasts out of 6) |
|---|---|---|---|---|---|
| L/R dmPFC | 3968 | | | | |
| L dmPFC (Fp2) | | -4 | 58 | 6 | 5 |
| R dmPFC (Fp2) | | 6 | 54 | 2 | 4 |
| R dmPFC (Fp2) | | 8 | 60 | 8 | 3 |
| L Amygdala / Hippocampus | 3448 | | | | |
| L Amygdala (LB) | | -26 | -6 | -22 | 6 |
| L Amygdala (LB) | | -24 | -6 | -28 | 5 |
| L Hippocampus (CA2) | | -34 | -18 | -16 | 4 |
| L Hippocampus (CA3) | | -30 | -18 | -18 | 3 |
| R Amygdala | 1920 | | | | |

| | | | | | |
|---|---|---|---|---|---|
| R Amygdala (LB) | | 24 | -4 | -28 | 6 |
| R Amygdala (LB) | | 20 | -4 | -26 | 5 |
| L/R vmPFC | 1392 | | | | |
| L vmPFC (Fo1) | | -2 | 50 | -22 | 5 |
| L vmPFC (Fp2) | | 0 | 48 | -18 | 4 |
| L vmPFC (Fo1) | | -2 | 44 | -22 | 3 |
| L TP | 1368 | | | | |
| L TP | | -50 | 10 | -32 | 6 |
| L TP | | -52 | 4 | -26 | 5 |
| L TP | | -44 | 8 | -20 | 4 |
| L AG | 1240 | | | | |
| L AG (PGa) | | -54 | -56 | 20 | 4 |
| L AG (PGp) | | -48 | -70 | 26 | 2 |

**Table S30. Multimodal convergence zones (overlap between the meta-analytic maps for all conceptual modalities).**

| Region | Cluster size ($mm^3$) | x | y | z | ALE | P | Z |
|---|---|---|---|---|---|---|---|
| *Trimodal: Action & Sound & Motion* | | | | | | | |
| L pMTG (core) | 264 | | | | | | |
| L pMTG | | -58 | -40 | 0 | 0.014 | 8.87E-05 | 3.749 |
| L pMTG | | -56 | -46 | 2 | 0.014 | 1.48E-04 | 3.618 |
| L IPL (PFcm) | 256 | -60 | -42 | 26 | 0.017 | 1.83E-05 | 4.127 |
| *Bimodal: Action & Sound* | | | | | | | |
| L dmPFC | 208 | -2 | 28 | 42 | 0.014 | 1.67E-04 | 3.587 |
| L pMTG (belt) | 168 | -58 | -40 | -6 | 0.013 | 2.02E-04 | 3.537 |
| L IPL (PF) | 72 | -56 | -38 | 28 | 0.012 | 5.93E-04 | 3.242 |
| *Bimodal: Action & Color* | | | | | | | |
| L IPS (hIP3) | 488 | -36 | -62 | 44 | 0.018 | 6.97E-07 | 4.826 |
| L FG4 | 88 | -36 | -46 | -20 | 0.009 | 7.04E-04 | 3.193 |
| L FG4 | 64 | -46 | -52 | -16 | 0.010 | 3.12E-04 | 3.421 |
| *Bimodal: Action & Emotion* | | | | | | | |
| L AG (PGp) | 328 | -46 | -64 | 22 | 0.023 | 2.91E-05 | 4.020 |
| *Bimodal: Action & Motion* | | | | | | | |
| L pMTG (posterior) | 2144 | -56 | -56 | 4 | 0.024 | 4.02E-08 | 5.366 |
| L IPL (PF/PFt) | 904 | -58 | -44 | 16 | 0.014 | 1.07E-04 | 3.702 |
| L pMTG (anterior) | 496 | -62 | -38 | -4 | 0.011 | 8.02E-04 | 3.155 |
| L IFG (tri) | 376 | -52 | 22 | 16 | 0.011 | 6.14E-04 | 3.232 |
| L FG3 | 72 | -34 | -42 | -22 | 0.011 | 6.95E-04 | 3.197 |

| | | | | | | | |
|---|---|---|---|---|---|---|---|
| *Bimodal: Action & Olfaction-Gustation* | | | | | | | |
| L OFC (Fo3) | 216 | -28 | 36 | -14 | 0.014 | 3.28E-05 | 3.991 |
| *Bimodal: Action & Shape* | | | | | | | |
| L LOC (hOc4la) | 1024 | -52 | -66 | -4 | 0.020 | 5.90E-07 | 4.859 |
| L PreCS | 736 | -44 | 2 | 28 | 0.010 | 9.67E-04 | 3.100 |
| L FG4 | 104 | -40 | -42 | -22 | 0.010 | 9.78E-04 | 3.106 |
| *Bimodal: Motion & Sound* | | | | | | | |
| L IPL (PFm/PF) | 248 | -60 | -44 | 26 | 0.016 | 3.29E-05 | 3.991 |
| L pMTG (belt) | 72 | -58 | -46 | 2 | 0.012 | 4.09E-04 | 3.346 |
| L pMTG (belt) | 56 | -62 | -42 | -4 | 0.011 | 4.74E-04 | 3.305 |
| *Bimodal: Emotion & Motion* | | | | | | | |
| L TPJ (PGp/PGa) | 176 | -54 | -60 | 20 | 0.011 | 4.51E-04 | 3.320 |

#### Supplementary analyses without low-level contrasts

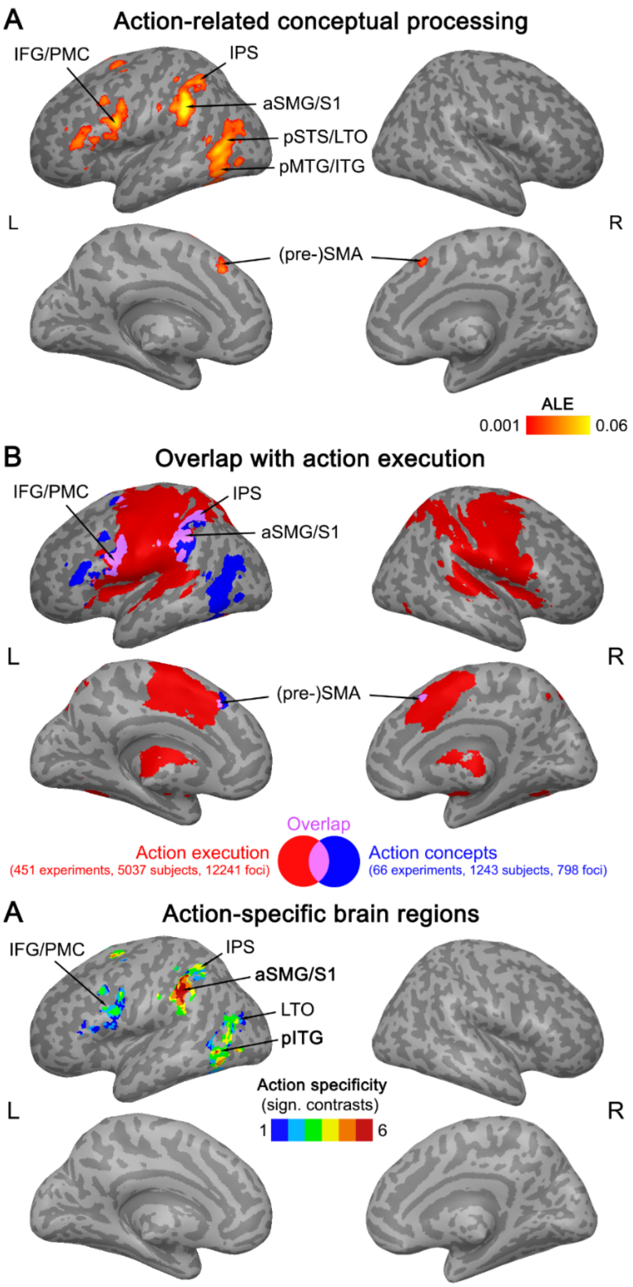

**Figure S1.** (A) Results of a supplementary ALE meta-analysis on action-related conceptual processing that excluded low-level contrasts (thresholded at a voxel-wise $p < 0.001$, cluster-wise $p < 0.05$ FWE-corrected). (B) Overlap (purple) between meta-analytic maps for action-related conceptual processing (blue) and real action execution (red). (C) Regions showing consistent engagement for conceptual processing related to action and no other modality, and higher activation likelihood for action than the other modalities (number of significant contrasts is displayed).

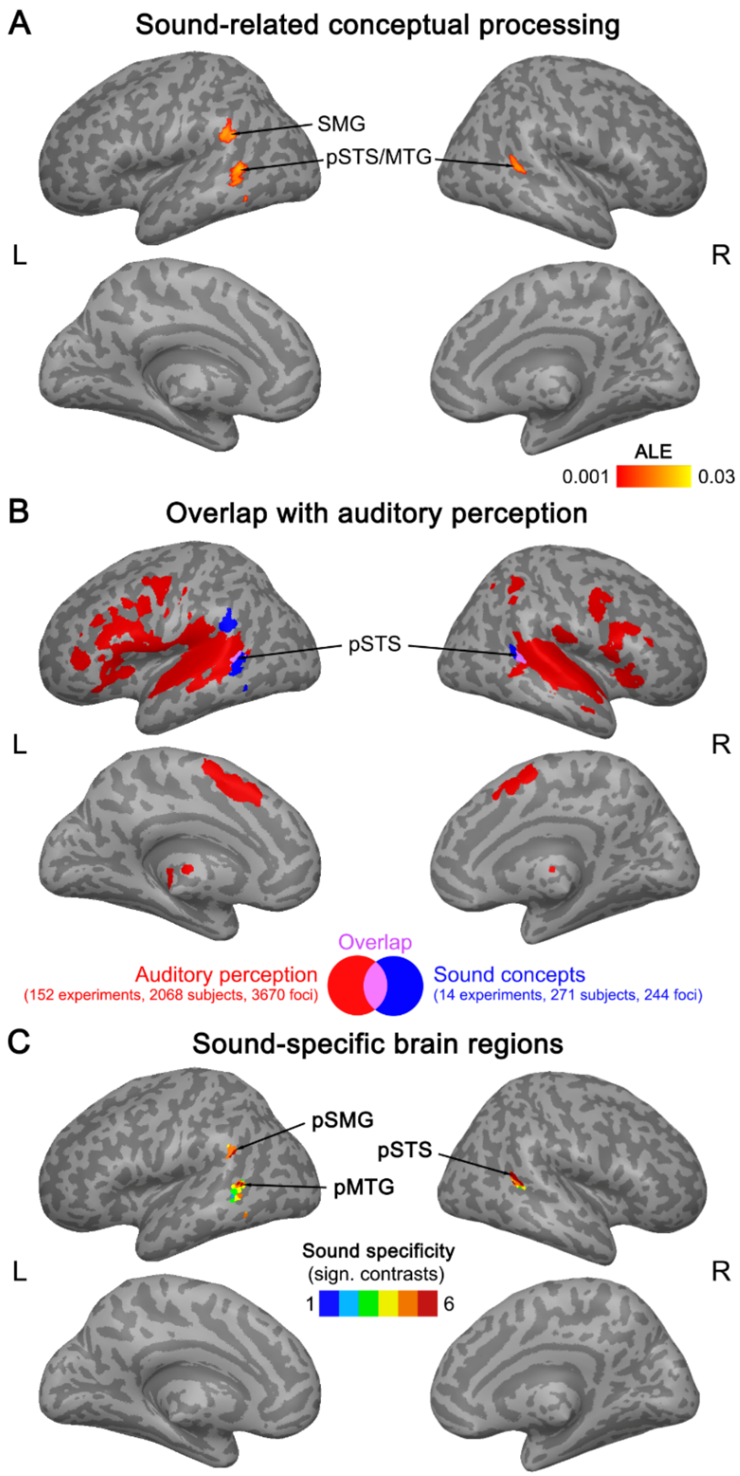

**Figure S2.** (A) Results of a supplementary ALE meta-analysis on sound-related conceptual processing that excluded low-level contrasts (thresholded at a voxel-wise $p < 0.001$, cluster-wise $p < 0.05$ FWE-corrected). (B) Overlap (purple) between meta-analytic maps for sound-related conceptual processing (blue) and real sound perception (red). (C) Regions showing consistent engagement for conceptual processing related to sound and no other modality, and higher activation likelihood for sound than the other modalities (number of significant contrasts is displayed).

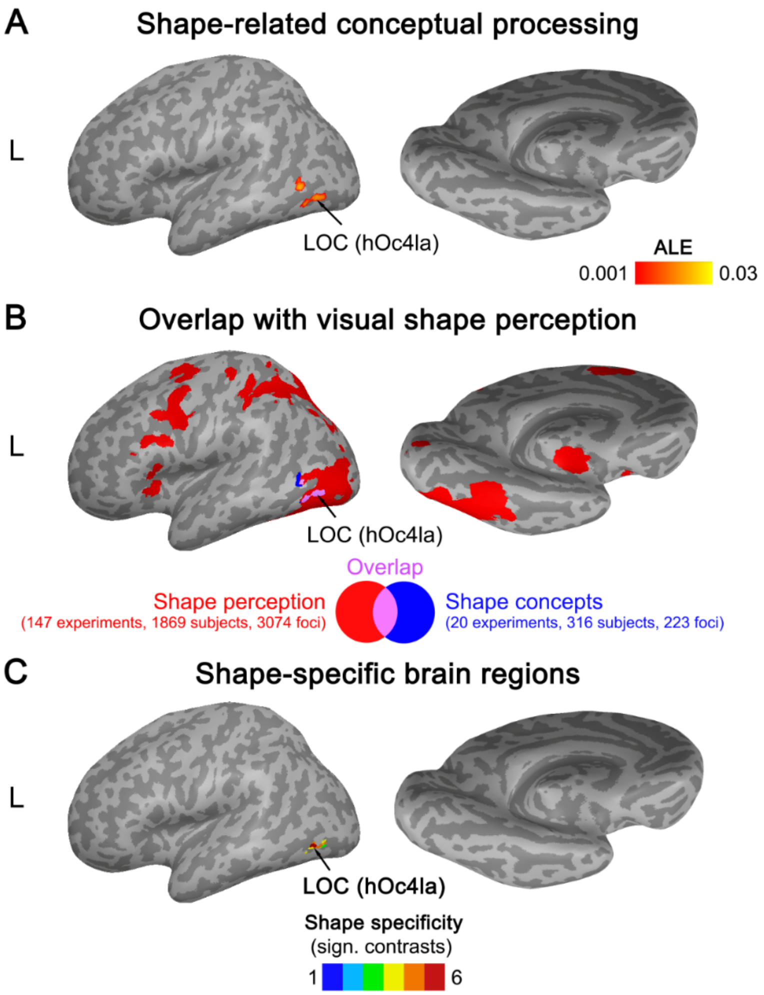

**Figure S3.** (A) Results of a supplementary ALE meta-analysis on shape-related conceptual processing that excluded low-level contrasts (thresholded at a voxel-wise $p < 0.001$, cluster-wise $p < 0.05$ FWE-corrected). (B) Overlap (purple) between meta-analytic maps for shape-related conceptual processing (blue) and real visual shape perception (red). (C) Regions showing consistent engagement for conceptual processing related to shape and no other modality, and higher activation likelihood for shape than the other modalities (number of significant contrasts is displayed).

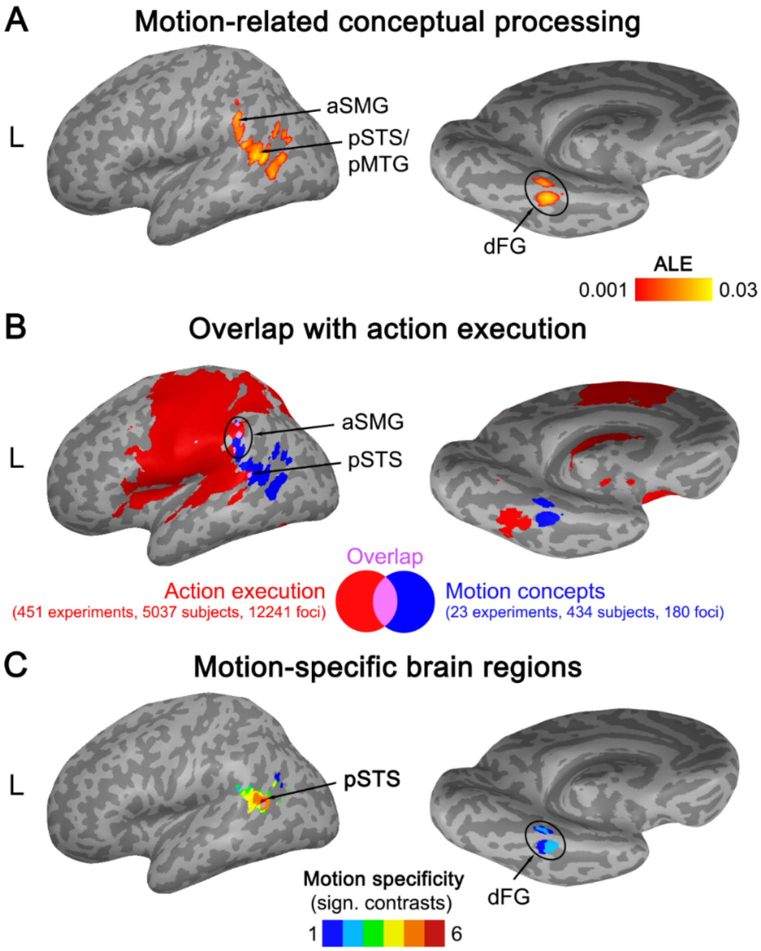

**Figure S4.** (A) Results of a supplementary ALE meta-analysis on motion-related conceptual processing that excluded low-level contrasts (thresholded at a voxel-wise $p < 0.001$, cluster-wise $p < 0.05$ FWE-corrected). (B) Overlap (purple) between meta-analytic maps for motion-related conceptual processing (blue) and real action execution (red). (C) Regions showing consistent engagement for conceptual processing related to motion and no other modality, and higher activation likelihood for motion than the other modalities (number of significant contrasts is displayed).

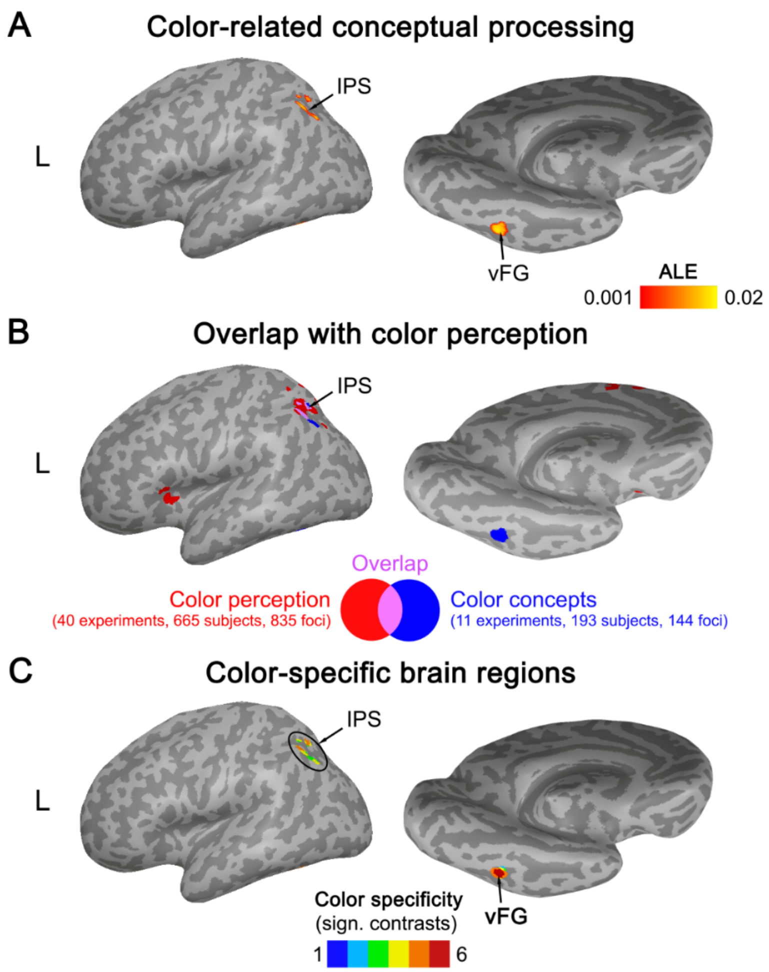

**Figure S5.** (A) Results of a supplementary ALE meta-analysis on color-related conceptual processing that excluded low-level contrasts (thresholded at a voxel-wise $p < 0.001$, cluster-wise $p < 0.05$ FWE-corrected). (B) Overlap (purple) between meta-analytic maps for color-related conceptual processing (blue) and real color perception (red). (C) Regions showing consistent engagement for conceptual processing related to color and no other modality, and higher activation likelihood for color than the other modalities (number of significant contrasts is displayed).

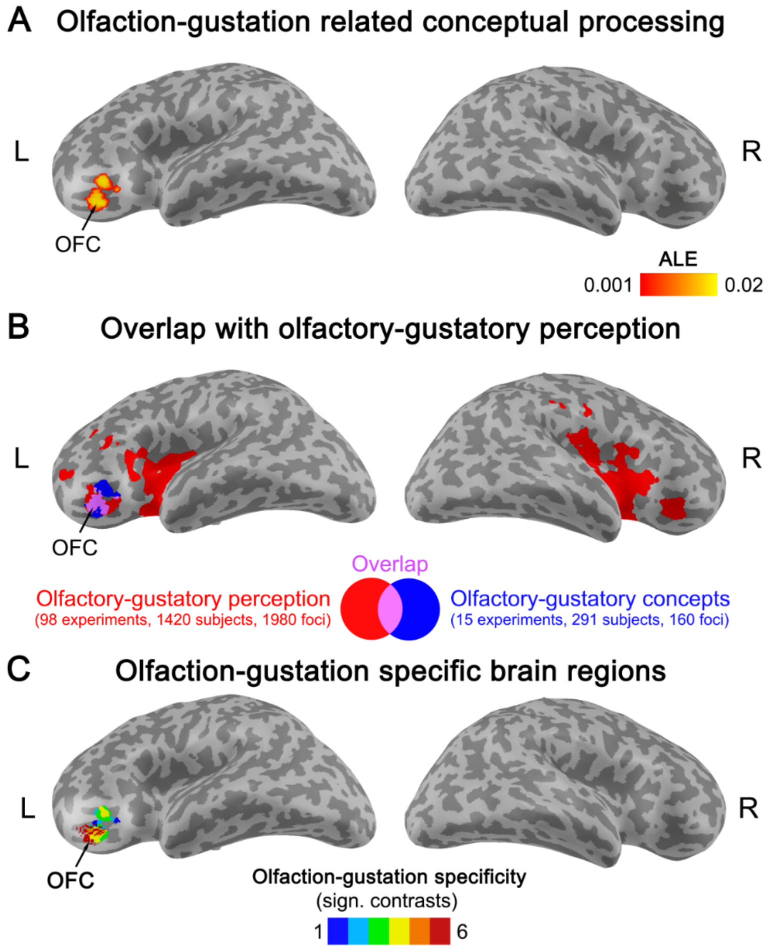

**Figure S6.** (A) Results of a supplementary ALE meta-analysis on olfaction-gustation related conceptual processing that excluded low-level contrasts (thresholded at a voxel-wise $p < 0.001$, cluster-wise $p < 0.05$ FWE-corrected). (B) Overlap (purple) between meta-analytic maps for olfaction-gustation related conceptual processing (blue) and real olfactory-gustatory perception (red). (C) Regions showing consistent engagement for conceptual processing related to olfaction-gustation and no other modality, and higher activation likelihood for olfaction-gustation than the other modalities (number of significant contrasts is displayed).

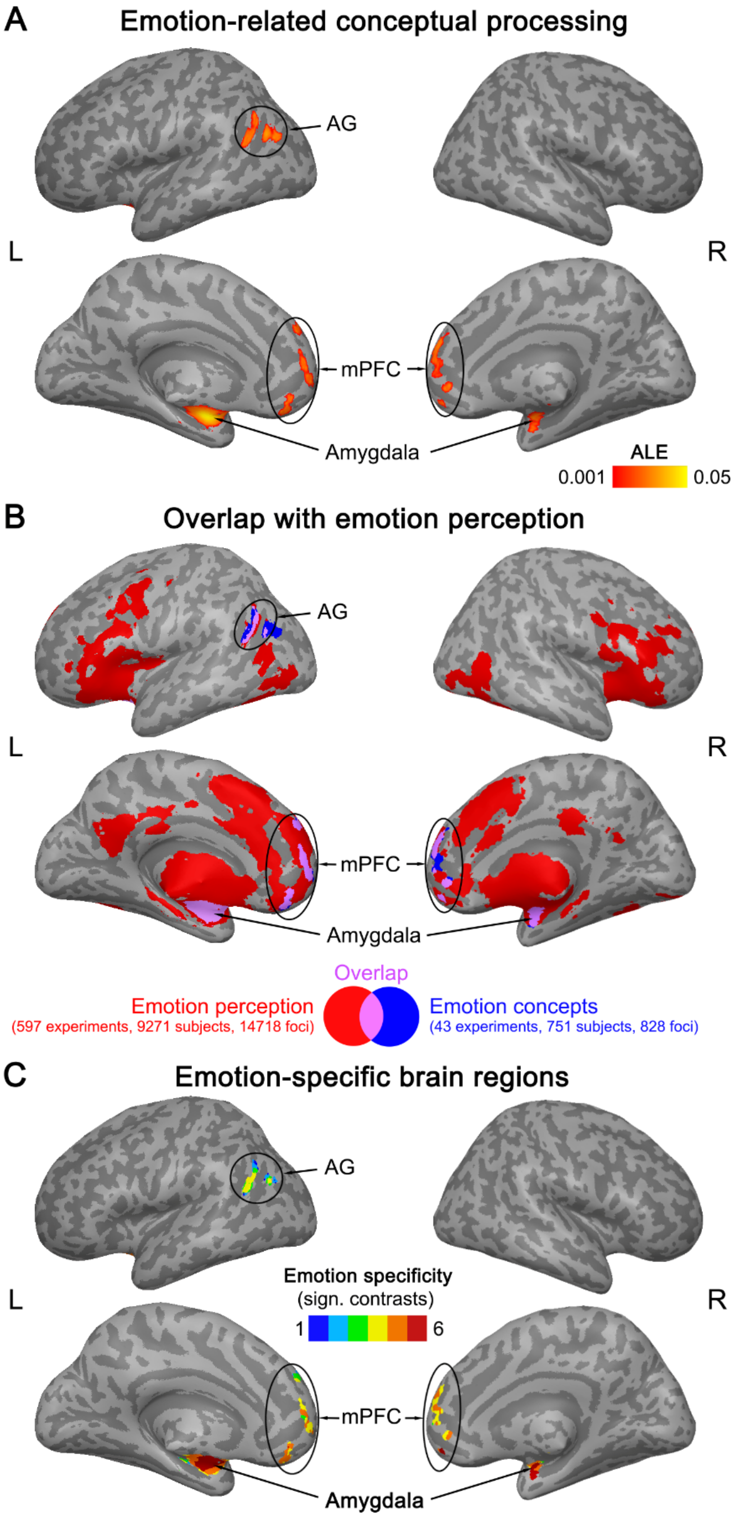

**Figure S7.** (A) Results of a supplementary ALE meta-analysis on emotion-related conceptual processing that excluded low-level contrasts (thresholded at a voxel-wise $p < 0.001$, cluster-wise $p < 0.05$ FWE-corrected). (B) Overlap (purple) between meta-analytic maps for emotion-related conceptual processing (blue) and real emotion perception (red). (C) Regions showing consistent engagement for conceptual processing related to emotion and no other modality, and higher activation likelihood for emotion than the other modalities (number of significant contrasts is displayed).

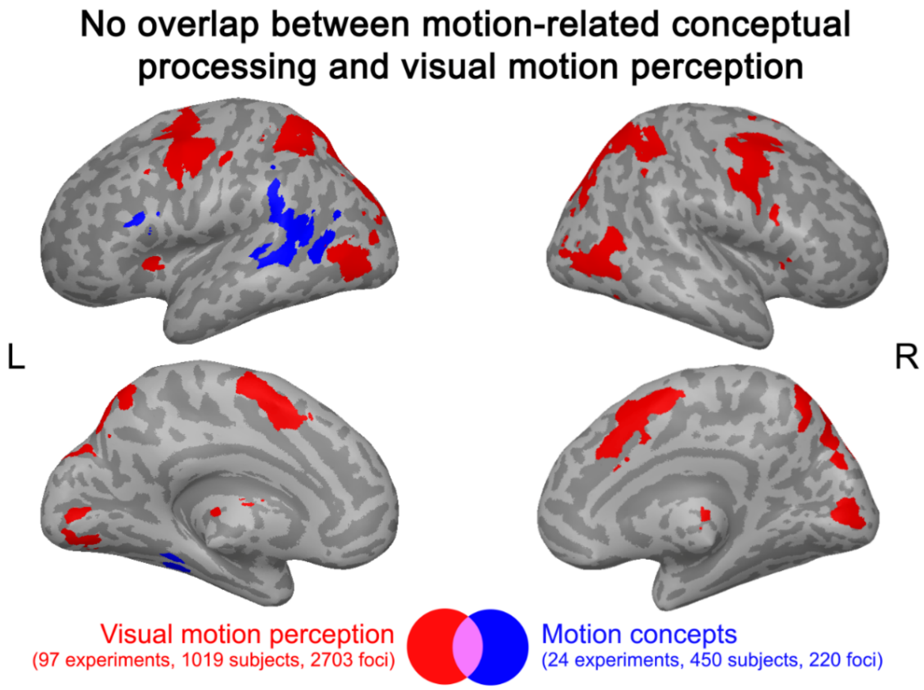

**Figure S8.** Motion-related conceptual processing (blue) did not overlap with real visual motion perception (red).

#### Experiment Tables

The following tables summarize essential information on the included experiments and contrasts for each ALE meta-analysis.

**Table S31. Experiments included in the ALE meta-analysis on conceptual processing related to action.**

| Exp. | Paper | Method | Participants (N) | Task | Stimulus Modality | Contrast | Space | Foci (N) | Contrast Type |
|---|---|---|---|---|---|---|---|---|---|
| 1 | Baumgaertner et al. (2007) | fMRI | 19 | semantic decision | spoken sentences | action sentences > non-action sentences | MNI | 9 | high |
| 1 | Baumgaertner et al. (2007) | fMRI | 19 | semantic decision | videos | action sentences > non-action sentences | MNI | 2 | high |
| 1 | Baumgaertner et al. (2007) | fMRI | 19 | semantic decision | spoken sentences, videos | (action > non-action sentences) ∩ (action > non-action videos) | MNI | 3 | high |
| 2 | Bedny et al. (2008) | fMRI | 12 | semantic decision | spoken words | action verbs > animal nouns | MNI | 1 | high |
| 3 | Bellebaum et al. (2013) | fMRI | 16 | perceptual decision | pictures | post- > pre-training for manipulation > scrambled images | MNI | 16 | high |
| 4 | Bonner et al. (2013) | fMRI | 20 | lexical decision | written words | manipulation words > sound words | MNI | 4 | high |
| 4 | Bonner et al. (2013) | fMRI | 20 | lexical decision | written words | manipulation words > pseudowords | MNI | 8 | high |
| 5 | Borghesani et al. (2019) | fMRI | 13 | semantic decision | pictures | tool pictures > animal pictures | MNI | 2 | high |
| 6 | Boronat et al. (2005) | fMRI | 15 | semantic decision | pictures, written words | manipulation > pseudowords and scrambled pictures | MNI | 7 | high |
| 6 | Boronat et al. (2005) | fMRI | 15 | semantic decision | written words | manipulable objects > pseudowords | MNI | 5 | high |
| 6 | Boronat et al. (2005) | fMRI | 15 | semantic decision | pictures | manipulable objects > scrambled pictures | MNI | 7 | high |
| 7 | Boulenger et al. (2009) | fMRI | 18 | passive reading | written sentences | early analysis: action sentences > hashmarks | MNI | 11 | low |
| 7 | Boulenger et al. (2009) | fMRI | 18 | passive reading | written sentences | late analysis: action sentences > hashmarks | MNI | 10 | low |
| 7 | Boulenger et al. (2009) | fMRI | 18 | passive reading | written sentences | early analysis: literal sentences > hash marks | MNI | 12 | low |
| 7 | Boulenger et al. (2009) | fMRI | 18 | passive reading | written sentences | late analysis: literal sentences > hash marks | MNI | 10 | low |
| 7 | Boulenger et al. (2009) | fMRI | 18 | passive reading | written sentences | early analysis: idiomatic sentences > hash marks | MNI | 14 | low |
| 7 | Boulenger et al. (2009) | fMRI | 18 | passive reading | written sentences | late analysis: idiomatic sentences > hash marks | MNI | 13 | low |

| Exp. | Paper | Method | Participants (N) | Task | Stimulus Modality | Contrast | Space | Foci (N) | Contrast Type |
|---|---|---|---|---|---|---|---|---|---|
| 8 | Bracci & Peelen (2013) | fMRI | 13 | working memory recall | pictures | object effectors > non-effector object control | TAL | 1 | high |
| 9 | Cappa et al. (1998) | PET | 13 | semantic decision | written words | tools > animals | TAL | 4 | high |
| 9 | Cappa et al. (1998) | PET | 13 | semantic decision | written words | associative semantics (tools) > pseudowords | TAL | 3 | high |
| 10 | Chao & Martin (2000) | fMRI | 5 | passive viewing | pictures | tool pictures > animal pictures | TAL | 2 | high |
| 10 | Chao & Martin (2000) | fMRI | 5 | naming | pictures | tool pictures > animal pictures | TAL | 3 | high |
| 11 | Chow et al. (2014) | fMRI | 24 | semantic decision | spoken narratives | action paragraphs > perception and emotion paragraphs | MNI | 2 | high |
| 12 | Damasio et al. (1996) | PET | 9 | picture naming | pictures | tools > unfamiliar faces | TAL | 3 | high |
| 12 | Damasio et al. (1996) | PET | 9 | picture naming | pictures | tools > unfamiliar faces | TAL | 3 | high |
| 12 | Damasio et al. (1996) | PET | 9 | picture naming | pictures | tools > unfamiliar faces | TAL | 3 | high |
| 13 | Damasio et al. (2001) | PET | 20 | naming | pictures | actions performed with a tool > orientation of faces | TAL | 21 | high |
| 13 | Damasio et al. (2001) | PET | 20 | naming | pictures | actions performed with a tool > concrete entities | TAL | 3 | high |
| 14 | De Grauwe et al. (2014) | fMRI | 20 | semantic decision | written words | motor words > non-motor words | MNI | 14 | high |
| 15 | Desai et al. (2010) | fMRI | 33 | semantic decision | spoken sentences | motor sentences > visual sentences | TAL | 2 | high |
| 15 | Desai et al. (2010) | fMRI | 33 | semantic decision | spoken sentences | motor sentences > abstract sentences | TAL | 2 | high |
| 15 | Desai et al. (2010) | fMRI | 33 | semantic decision | spoken sentences | hand associations > arm associations | TAL | 17 | high |
| 16 | Desai et al. (2011) | fMRI | 22 | semantic decision | written sentences | literal action sentences > abstract sentences | TAL | 9 | high |
| 16 | Desai et al. (2011) | fMRI | 22 | semantic decision | written sentences | metaphoric action sentences > abstract sentences | TAL | 8 | high |
| 16 | Desai et al. (2011) | fMRI | 22 | semantic decision | written sentences | literal action sentences > metaphoric action sentences | TAL | 1 | high |
| 17 | Desai et al. (2013) | fMRI | 27 | semantic decision | written sentences | metaphorical action sentences > abstract sentences | TAL | 3 | high |
| 17 | Desai et al. (2013) | fMRI | 27 | semantic decision | written sentences | idiomatic action sentences > abstract sentences | TAL | 4 | high |
| 17 | Desai et al. (2013) | fMRI | 27 | semantic decision | written sentences | literal sentences > abstract sentences | TAL | 12 | high |

| Exp. | Paper | Method | Participants (N) | Task | Stimulus Modality | Contrast | Space | Foci (N) | Contrast Type |
|---|---|---|---|---|---|---|---|---|---|
| 18 | Desai et al. (2016) | fMRI | 40 | passive reading | written sentences | correlation with noun manipulability | TAL | 11 | high |
| 19 | Desai et al. (2016) | fMRI | 31 | passive reading | written narratives | correlation with noun manipulability | TAL | 15 | high |
| 20 | Dreyer et al. (2018) | fMRI | 28 | passive reading | written words | tool words > hash marks | MNI | 7 | low |
| 21 | Fernandino et al. (2016) | fMRI | 44 | semantic decision | written words | manipulation > other attributes | TAL | 1 | high |
| 21 | Fernandino et al. (2016) | fMRI | 44 | semantic decision | written words | manipulation > other attributes except shape | TAL | 14 | high |
| 22 | Gilead et al. (2016) | fMRI | 28 | semantic decision | written sentences | actions: how > why | MNI | 9 | high |
| 23 | Goldberg et al. (2006a) | fMRI | 13 | word generation, semantic decision | written words | functional words > visual words | TAL | 2 | high |
| 24 | Grossman et al. (2002a) | fMRI | 16 | semantic decision | written words | implements > animal words | TAL | 2 | high |
| 24 | Grossman et al. (2002a) | fMRI | 16 | semantic decision | written words | implements > pseudowords | TAL | 2 | high |
| 25 | Hauk et al. (2004) | fMRI | 14 | passive reading | written words | action words > hash marks | MNI | 7 | low |
| 25 | Hauk et al. (2004) | fMRI | 14 | passive reading | written words | face words > hash marks | MNI | 9 | low |
| 25 | Hauk et al. (2004) | fMRI | 14 | passive reading | written words | arm words > hash marks | MNI | 10 | low |
| 25 | Hauk et al. (2004) | fMRI | 14 | passive reading | written words | leg words > hash marks | MNI | 10 | low |
| 26 | Hauk et al. (2008) | fMRI | 21 | passive reading | written words | correlation with action-relatedness | MNI | 5 | high |
| 27 | Hoenig et al. (2008) | fMRI | 20 | semantic decision | written words | artifactual items > natural items | MNI | 6 | high |
| 28 | Hoeren et al. (2013) | fMRI | 30 | semantic decision | pictures, videos | context tool use incorrect > context tool use correct | MNI | 8 | high |
| 28 | Hoeren et al. (2013) | fMRI | 30 | semantic decision | pictures, videos | correct hand postures > incorrect hand postures | MNI | 2 | high |
| 29 | Johnson-Frey et al. (2005) | fMRI | 13 | mental imagery | spoken words | right hand: tool use gestures > non-meaningful gestures | TAL | 26 | high |
| 29 | Johnson-Frey et al. (2005) | fMRI | 13 | mental imagery | spoken words | left hand: tool use gestures > non-meaningful gestures | TAL | 13 | high |

| Exp. | Paper | Method | Participants (N) | Task | Stimulus Modality | Contrast | Space | Foci (N) | Contrast Type |
|---|---|---|---|---|---|---|---|---|---|
| 30 | Just et al. (2010) | fMRI | 11 | mental imagery | written words | manipulation factor | MNI | 4 | low |
| 31 | Kable & Chatterjee (2006) | fMRI | 9 | semantic decision | videos | new person, new action > new person, old action | MNI | 5 | high |
| 32 | Kana et al. (2012) | fMRI | 24 | semantic decision | written sentences | action verbs > mental verbs | MNI | 8 | high |
| 33 | Kellenbach et al. (2003) | PET | 9 | semantic decision | pictures | action: semantic decision on objects > perceptual decision on scrambled pictures | MNI | 9 | high |
| 33 | Kellenbach et al. (2003) | PET | 9 | semantic decision | pictures | manipulable function: semantic decision on object > perceptual decision on scrambled pictures | MNI | 9 | high |
| 34 | Kemmerer et al. (2008) | fMRI | 16 | semantic decision | written words | hitting verbs > meaningless symbols | MNI | 14 | low |
| 34 | Kemmerer et al. (2008) | fMRI | 16 | semantic decision | written words | cutting verbs > meaningless symbols | MNI | 23 | low |
| 35 | Khader et al. (2010) | fMRI | 17 | word generation task, rhyme generation task | written sentences | verb generation > rhyme generation | TAL | 2 | high |
| 35 | Khader et al. (2010) | fMRI | 17 | word generation task, rhyme generation task | written sentences | verb generation > letter search | TAL | 5 | high |
| 36 | Kleineberg et al. (2018) | fMRI | 18 | semantic decision | pictures | manipulation > function and monetary value | MNI | 6 | high |
| 37 | Kuhnke et al. (2020) | fMRI | 40 | semantic decision | written words | high > low action words | MNI | 65 | high |
| 38 | Lauro et al. (2013) | fMRI | 24 | semantic decision | written sentences | action sentences involving upper-limb > cognition sentences | MNI | 8 | high |
| 38 | Lauro et al. (2013) | fMRI | 24 | semantic decision | written sentences | idiomatic action sentences involving upper-limb > cognition sentences | MNI | 3 | high |
| 38 | Lauro et al. (2013) | fMRI | 24 | semantic decision | written sentences | literal action sentences involving upper-limb > cognition sentences | MNI | 4 | high |
| 39 | Macdonald & Culham (2015) | fMRI | 12 | passive viewing | pictures | tool pictures > non-tool pictures | TAL | 16 | high |
| 40 | Martin et al. (1995) | PET | 12 | word generation | pictures | action word generation > color word generation | TAL | 5 | high |
| 40 | Martin et al. (1995) | PET | 12 | word generation | written words | action word generation > color word generation | TAL | 4 | high |
| 40 | Martin et al. (1995) | PET | 12 | word generation | pictures | action word generation > object naming | TAL | 10 | high |
| 41 | Martin et al. (1996) | PET | 16 | passive naming, naming | pictures | silently naming tools > viewing non-sense objects | TAL | 9 | high |

| Exp. | Paper | Method | Participants (N) | Task | Stimulus Modality | Contrast | Space | Foci (N) | Contrast Type |
|---|---|---|---|---|---|---|---|---|---|
| 42 | Moseley et al. (2012) | fMRI | 18 | passive reading | written words | face action words > hash marks | MNI | 17 | low |
| 42 | Moseley et al. (2012) | fMRI | 18 | passive reading | written words | arm action words > hash marks | MNI | 18 | low |
| 43 | Neudorf et al. (2019) | fMRI | 25 | mental imagery, lexical decision | written words | (related > unrelated) for (action > object priming) | MNI | 13 | high |
| 44 | Nijhof & Willems (2015) | fMRI | 18 | passive listening | spoken narratives | action sentences > mentalizing sentences | MNI | 3 | high |
| 45 | Noppeney et al. (2005) | fMRI | 12 | semantic decision | spoken words | action words > auditory, motion, visual words | TAL | 2 | high |
| 45 | Noppeney et al. (2005) | fMRI | 12 | semantic decision | spoken words | hand action words > body movement words | TAL | 1 | high |
| 46 | Noppeney et al. (2005) | fMRI | 15 | semantic decision | written words | action words > auditory, motion, visual words | TAL | 2 | high |
| 46 | Noppeney et al. (2005) | fMRI | 27 | semantic decision | spoken words, written words | action words > auditory, motion, visual words | TAL | 2 | high |
| 47 | Noppeney et al. (2006) | fMRI | 22 | semantic decision | pictures, written words | tools > animals | TAL | 7 | high |
| 47 | Noppeney et al. (2006) | fMRI | 22 | semantic decision | pictures, written words | explicit > implicit task for tools > animals | TAL | 3 | high |
| 48 | Peelen et al. (2013) | fMRI | 16 | semantic decision | spoken words | tool words > animals and nonmanipulable object words | TAL | 1 | high |
| 49 | Phillips et al. (2002) | PET | 12 | semantic decision | pictures, written words | (action > size retrieval) ∩ (action retrieval > screen size control) | MNI | 3 | high |
| 49 | Phillips et al. (2002) | PET | 12 | semantic decision | pictures, written words | tools > fruit | MNI | 2 | high |
| 50 | Popp et al. (2019a) | fMRI | 22 | lexical decision | written words | action verbs > sound verbs | MNI | 4 | high |
| 50 | Popp et al. (2019a) | fMRI | 22 | lexical decision | written words | action ratings > sound ratings | MNI | 4 | high |
| 50 | Popp et al. (2019a) | fMRI | 22 | lexical decision | written words | action verbs > pseudoverbs | MNI | 46 | high |
| 51 | Popp et al. (2019b) | fMRI | 30 | semantic decision | written words | action > sound verbs | MNI | 7 | high |
| 51 | Popp et al. (2019b) | fMRI | 30 | semantic decision | written words | semantically-related action verbs > semantically-related sound verbs | MNI | 12 | high |
| 51 | Popp et al. (2019b) | fMRI | 30 | lexical decision | written words | repetition suppression by action verbs | MNI | 1 | high |
| 51 | Popp et al. (2019b) | fMRI | 30 | lexical decision | written words | repetition enhancement by action verbs | MNI | 5 | high |

| Exp. | Paper | Method | Participants (N) | Task | Stimulus Modality | Contrast | Space | Foci (N) | Contrast Type |
|---|---|---|---|---|---|---|---|---|---|
| 52 | Raposo et al. (2009) | fMRI | 14 | passive listening | spoken words | action words > non-action words | MNI | 1 | high |
| 53 | Raposo et al. (2009) | fMRI | 22 | passive listening | spoken sentences | literal sentences > idiomatic sentences | MNI | 1 | high |
| 54 | Ruby & Decety (2001) | PET | 10 | mental imagery | pictures, spoken sentences | first-person perspective: (auditory action sentences > auditory non-action sentences ) ∩ (imagery of interaction with objects > passively viewing objects) | MNI | 9 | high |
| 54 | Ruby & Decety (2001) | PET | 10 | mental imagery | pictures, spoken sentences | third-person perspective: (auditory action sentences > auditory non-action sentences ) ∩ (imagery of interaction with objects > passively viewing objects) | MNI | 7 | high |
| 55 | Rueschemeyer et al. (2007) | fMRI | 20 | lexical decision | written words | (simple motor > simple abstract verbs) > (complex motor > complex abstract verbs) | TAL | 3 | high |
| 55 | Rueschemeyer et al. (2007) | fMRI | 20 | lexical decision | written words | simple verbs: motor > abstract | TAL | 9 | high |
| 55 | Rueschemeyer et al. (2007) | fMRI | 20 | lexical decision | written words | complex verbs: motor > abstract | TAL | 2 | high |
| 55 | Rueschemeyer et al. (2007) | fMRI | 20 | lexical decision | written words | (task > baseline) ∩ (simple motor > simple abstract verbs) | TAL | 5 | high |
| 56 | Rueschemeyer et al. (2010) | fMRI | 14 | lexical decision | written words | objects: manipulability necessary for function > manipulability not necessary for function | MNI | 5 | high |
| 57 | Speer et al. (2009) | fMRI | 28 | passive reading | written narratives | manipulable-object change > other changes | MNI | 1 | high |
| 57 | Speer et al. (2009) | fMRI | 28 | passive reading | written narratives | manipulable-object change > clause onset | MNI | 10 | high |
| 58 | Spunt & Lieberman (2012) | fMRI | 21 | semantic decision | videos, written sentences | actions: video (how > why) ∩ text (how > why) | MNI | 6 | high |
| 59 | Tettamanti et al. (2005) | fMRI | 17 | passive listening | spoken sentences | action sentences > abstract sentences | MNI | 1 | high |
| 59 | Tettamanti et al. (2005) | fMRI | 17 | passive listening | spoken sentences | mouth sentences > abstract sentences | MNI | 5 | high |
| 59 | Tettamanti et al. (2005) | fMRI | 17 | passive listening | spoken sentences | hand sentences > abstract sentences | MNI | 8 | high |
| 59 | Tettamanti et al. (2005) | fMRI | 17 | passive listening | spoken sentences | leg sentences > abstract sentences | MNI | 5 | high |
| 60 | Tettamanti et al. (2008) | fMRI | 18 | passive listening | spoken sentences | action sentences > abstract sentences | MNI | 10 | high |
| 60 | Tettamanti et al. (2008) | fMRI | 18 | passive listening | spoken sentences | (action-related affirmative > action-related negative sentences) > (abstract affirmative > abstract negative sentences) | MNI | 6 | high |

| Exp. | Paper | Method | Participants (N) | Task | Stimulus Modality | Contrast | Space | Foci (N) | Contrast Type |
|---|---|---|---|---|---|---|---|---|---|
| 61 | Tomasino et al. (2007) | fMRI | 15 | mental imagery, perceptual decision | written sentences | motor sentences > non-motor sentences | MNI | 1 | high |
| 61 | Tomasino et al. (2007) | fMRI | 15 | mental imagery, perceptual decision | written sentences | imagery > letter detection for motor > non-motor sentences | MNI | 1 | high |
| 62 | Tomasino et al. (2014) | fMRI | 24 | mental imagery | written words | (imagery: action > abstract) > (letter detection: action > abstract) | MNI | 15 | high |
| 63 | van Ackeren et al. (2016) | fMRI | 22 | semantic decision | spoken narratives | indirect request to act > indirect reply | MNI | 3 | high |
| 64 | van Dam et al. (2010) | fMRI | 14 | semantic decision | written words | action verbs > abstract verbs | MNI | 5 | high |
| 64 | van Dam et al. (2010) | fMRI | 14 | semantic decision | written words | highly-specific action verbs > basic action verbs | MNI | 4 | high |
| 64 | van Dam et al. (2010) | fMRI | 14 | semantic decision | written words | basic action verbs > highly-specific action verbs | MNI | 1 | high |
| 64 | van Dam et al. (2010) | fMRI | 14 | semantic decision | written words | basic action verbs > fixation | MNI | 8 | low |
| 64 | van Dam et al. (2010) | fMRI | 14 | semantic decision | written words | highly-specific action verbs > fixation | MNI | 8 | low |
| 65 | van Dam et al. (2019) | fMRI | 17 | semantic decision | pictures | action pictures > manner pictures | TAL | 8 | high |
| 65 | van Dam et al. (2019) | fMRI | 17 | semantic decision | pictures | action pictures > scrambled action pictures | TAL | 6 | high |
| 65 | van Dam et al. (2019) | fMRI | 17 | lexical decision, go/no-go | written words | action verbs > pseudoverbs | TAL | 5 | high |
| 66 | Warburton et al. (1996) | PET | 4 | word generation | spoken words | verb generation > passive listening | TAL | 10 | high |
| 66 | Warburton et al. (1996) | PET | 4 | word generation | spoken words | verb generation > rest | TAL | 12 | low |
| 67 | Warburton et al. (1996) | PET | 9 | word generation | spoken words | verb generation > rest | TAL | 9 | low |
| 67 | Warburton et al. (1996) | PET | 9 | word generation | spoken words | verb generation > rest | TAL | 14 | low |
| 68 | Warburton et al. (1996) | PET | 6 | word generation | spoken words | verb generation > silent repetition of pseudowords | TAL | 5 | high |
| 68 | Warburton et al. (1996) | PET | 6 | word generation | spoken words | verb generation > rest | TAL | 11 | low |
| 69 | Warburton et al. (1996) | PET | 9 | word generation | spoken words | verb generation > noun generation | TAL | 6 | high |
| 69 | Warburton et al. (1996) | PET | 9 | word generation | spoken words | verb-noun comparison > rest | TAL | 5 | low |
| 70 | Webster et al. (2017) | fMRI | 17 | semantic decision | sounds | action sound > vocalization sound | TAL | 7 | high |

| Exp. | Paper | Method | Participants (N) | Task | Stimulus Modality | Contrast | Space | Foci (N) | Contrast Type |
|---|---|---|---|---|---|---|---|---|---|
| 71 | Wheatley et al. (2005) | fMRI | 15 | passive reading | written words | tools > animate objects | TAL | 7 | high |
| 72 | Willems et al. (2010) | fMRI | 20 | lexical decision | written words | lexical decision: manual verbs > non-manual verbs | MNI | 3 | high |
| 72 | Willems et al. (2010) | fMRI | 20 | lexical decision | written words | imagery: manual verbs > non-manual verbs | MNI | 8 | high |
| 72 | Willems et al. (2010) | fMRI | 20 | lexical decision | written words | [lexical decision on manual verbs > rest] ∩ [lexical decision on non-manual verbs > rest] | MNI | 25 | low |
| 72 | Willems et al. (2010) | fMRI | 20 | lexical decision | written words | [imagery for manual verbs > rest] ∩ [Imagery for non-manual verbs > rest] | MNI | 16 | low |
| 73 | Wriessnegger et al. (2016) | fMRI | 21 | mental imagery | pictograms | motor imagery joint action > motor imagery single action | MNI | 2 | high |
| 73 | Wriessnegger et al. (2016) | fMRI | 21 | mental imagery | pictograms | motor imagery joint action > fixation | MNI | 2 | low |
| 73 | Wriessnegger et al. (2016) | fMRI | 21 | mental imagery | pictograms | motor imagery single action > fixation | MNI | 3 | low |
| 73 | Wriessnegger et al. (2016) | fMRI | 21 | mental imagery | pictograms | motor imagery > fixation | MNI | 5 | low |
| 74 | Yang & Shu (2014) | fMRI | 20 | mental imagery, passive reading | written words | hand actions: motor imagery > passive reading | TAL | 6 | high |
| 74 | Yang & Shu (2014) | fMRI | 20 | mental imagery, passive reading | written words | hand and tool actions: motor imagery > passive reading | TAL | 6 | high |
| 74 | Yang & Shu (2014) | fMRI | 20 | mental imagery, passive reading | written words | tool actions: motor imagery > passive reading | TAL | 6 | high |
| 74 | Yang & Shu (2014) | fMRI | 20 | mental imagery, passive reading | written words | motor imagery > rest | TAL | 11 | low |

**Table S32. Experiments included in the ALE meta-analysis on conceptual processing related to sound.**

| Exp. | Paper | Method | Participants (N) | Task | Stimulus Modality | Contrast | Space | Foci (N) | Contrast Type |
|---|---|---|---|---|---|---|---|---|---|
| 1 | Bonner et al. (2013) | fMRI | 20 | lexical decision | written words | sound words > sight words | MNI | 1 | high |
| 1 | Bonner et al. (2013) | fMRI | 20 | lexical decision | written words | sound words > pseudowords | MNI | 5 | high |
| 2 | Fernandino et al. (2016) | fMRI | 44 | semantic decision | written words | sound > other attributes | TAL | 7 | high |
| 2 | Fernandino et al. (2016) | fMRI | 44 | semantic decision | written words | sound > other attributes except motion | TAL | 3 | high |
| 3 | Goldberg et al. (2006b) | fMRI | 14 | semantic decision | written words | auditory words > gustatory, tactile, visual words | TAL | 1 | high |
| 4 | Halpern & Zatorre (1999) | PET | 8 | mental imagery | none | auditory imagery > listening | TAL | 16 | high |
| 4 | Halpern & Zatorre (1999) | PET | 8 | mental imagery | none | auditory imagery after cue > imagery of control sequence | TAL | 10 | high |
| 4 | Halpern & Zatorre (1999) | PET | 8 | mental imagery | none | auditory imagery of control sequence > listening | TAL | 9 | high |
| 5 | Halpern et al. (2004) | fMRI | 10 | mental imagery | written words | timbre imagery > visual imagery | TAL | 2 | high |
| 5 | Halpern et al. (2004) | fMRI | 10 | mental imagery | written words | timbre imagery > visual imagery | TAL | 7 | high |
| 5 | Halpern et al. (2004) | fMRI | 10 | mental imagery | written words | timbre imagery ∩ timbre perception | TAL | 3 | high |
| 6 | Hoenig et al. (2011) | fMRI | 40 | semantic decision | pictures, written words | (musicians > non-musicians) for (musical instruments > control objects) | MNI | 12 | low |
| 7 | Kellenbach et al. (2001) | PET | 10 | semantic decision | written words | sound verifications > color verifications | MNI | 2 | high |
| 7 | Kellenbach et al. (2001) | PET | 10 | semantic decision | written words | sound verifications > size verifications | MNI | 5 | high |
| 7 | Kellenbach et al. (2001) | PET | 10 | semantic decision | written words | sound verifications > color and size verifications | MNI | 4 | high |
| 7 | Kellenbach et al. (2001) | PET | 10 | semantic decision | written words | sound verifications > letter-strings | MNI | 12 | high |
| 8 | Kiefer et al. (2008) | fMRI | 16 | lexical decision | written words | (words with > without auditory features) ∩ (object sounds > silence) | MNI | 1 | high |
| 8 | Kiefer et al. (2008) | fMRI | 16 | lexical decision | written words | words with > without auditory features | MNI | 8 | high |
| 8 | Kiefer et al. (2008) | fMRI | 16 | lexical decision | written words | correlation of auditory conceptual feature ratings | MNI | 1 | high |
| 9 | Kuhnke et al. (2020) | fMRI | 40 | semantic decision | written words | high > low sound words | MNI | 21 | high |
| 10 | Noppeney et al. (2002) | PET | 12 | semantic decision | spoken words | auditory words > visual and abstract words | TAL | 1 | high |

| Exp. | Paper | Method | Participants (N) | Task | Stimulus Modality | Contrast | Space | Foci (N) | Contrast Type |
|---|---|---|---|---|---|---|---|---|---|
| 11 | Popp et al. (2019a) | fMRI | 22 | lexical decision | written words | sound verbs > action verbs | MNI | 18 | high |
| 11 | Popp et al. (2019a) | fMRI | 22 | lexical decision | written words | sound verbs > pseudoverbs | MNI | 36 | high |
| 12 | Popp et al. (2019b) | fMRI | 30 | semantic decision | written words | sound verbs > action verbs | MNI | 8 | high |
| 12 | Popp et al. (2019b) | fMRI | 30 | semantic decision | written words | semantically-related sound verbs > semantically-related action verbs | MNI | 2 | high |
| 12 | Popp et al. (2019b) | fMRI | 30 | lexical decision | written words | repetition enhancement by sound verbs | MNI | 9 | high |
| 13 | Wheeler et al. (2000) | fMRI | 18 | episodic memory recall | written words | sound recall > picture recall | TAL | 9 | high |
| 14 | Yoo et al. (2001) | fMRI | 12 | mental imagery | none | auditory imagery > rest | TAL | 19 | low |
| 15 | Zatorre et al. (1996) | PET | 12 | perceptual decision, mental imagery | written words | decision on pitch in perceptual and imagery task > decision on word length | TAL | 21 | high |
| 16 | Zvyagintsev et al. (2013) | fMRI | 15 | mental imagery | none | auditory imagery > visual imagery | TAL | 15 | high |
| 16 | Zvyagintsev et al. (2013) | fMRI | 15 | mental imagery | none | auditory imagery > backwards counting | TAL | 7 | high |

**Table S33. Experiments included in the ALE meta-analysis on conceptual processing related to visual shape.**

| Exp. | Paper | Method | Participants (N) | Task | Stimulus Modality | Contrast | Space | Foci (N) | Contrast Type |
|---|---|---|---|---|---|---|---|---|---|
| 1 | Bonner et al. (2013) | fMRI | 20 | lexical decision | written words | visual words > sound words | MNI | 1 | high |
| 1 | Bonner et al. (2013) | fMRI | 20 | lexical decision | written words | visual words > pseudowords | MNI | 6 | high |
| 2 | Cappa et al. (1998) | PET | 13 | semantic decision | written words | visual > associative | TAL | 12 | high |
| 2 | Cappa et al. (1998) | PET | 13 | semantic decision | written words | visual semantics (animals) > pseudowords | TAL | 8 | high |
| 2 | Cappa et al. (1998) | PET | 13 | semantic decision | written words | visual semantics (tools) > pseudowords | TAL | 6 | high |
| 3 | D'Esposito et al. (1997) | fMRI | 7 | mental imagery | spoken words, sounds | concrete > abstract | TAL | 3 | high |
| 4 | Fernandino et al. (2016) | fMRI | 44 | semantic decision | written words | shape > other attributes | TAL | 24 | high |
| 4 | Fernandino et al. (2016) | fMRI | 44 | semantic decision | written words | shape > other attributes except color | TAL | 17 | high |
| 5 | Gauvin et al. (2019) | fMRI | 19 | picture naming | pictures | visually similar > congruent distractor word | MNI | 2 | high |
| 6 | Goldberg et al. (2006a) | fMRI | 13 | word generation, semantic decision | written words | visual words > functional words | TAL | 2 | high |
| 7 | Grossman et al. (2002a) | fMRI | 16 | semantic decision | written words | animal words > pseudowords | TAL | 1 | high |
| 8 | Hauk et al. (2008) | fMRI | 21 | passive reading | written words | correlation with imageability | MNI | 7 | high |
| 9 | Ishai et al. (2000) | fMRI | 9 | mental imagery | rest | imagery > passively viewing gray square | TAL | 16 | high |
| 10 | Kellenbach et al. (2001) | PET | 10 | semantic decision | written words | size verifications > color verifications | MNI | 1 | high |
| 10 | Kellenbach et al. (2001) | PET | 10 | semantic decision | written words | size verifications > sound verifications | MNI | 5 | high |
| 10 | Kellenbach et al. (2001) | PET | 10 | semantic decision | written words | size verifications > letter-strings | MNI | 5 | low |
| 11 | Khader et al. (2010) | fMRI | 17 | word generation task, rhyme generation task | written sentences | noun generation > rhyme generation | TAL | 8 | high |
| 11 | Khader et al. (2010) | fMRI | 17 | word generation task, rhyme generation task | written sentences | noun generation > letter search | TAL | 4 | high |
| 12 | Kosslyn et al. (1995) | PET | 12 | mental imagery | spoken words | imagery small sized images > passive listening | TAL | 3 | high |
| 12 | Kosslyn et al. (1995) | PET | 12 | mental imagery | spoken words | imagery medium sized images > passive listening | TAL | 1 | high |

| Exp. | Paper | Method | Participants (N) | Task | Stimulus Modality | Contrast | Space | Foci (N) | Contrast Type |
|---|---|---|---|---|---|---|---|---|---|
| 12 | Kosslyn et al. (1995) | PET | 12 | mental imagery | spoken words | imagery large sized images > passive listening | TAL | 3 | high |
| 13 | Mellet et al. (1998) | PET | 8 | mental imagery | spoken sentences | imagery (concrete) > listen (abstract) | TAL | 9 | high |
| 13 | Mellet et al. (1998) | PET | 8 | mental imagery | spoken sentences | imagery > rest | TAL | 15 | low |
| 14 | Nagels et al. (2013) | fMRI | 17 | perceptual decision | videos, spoken sentences | shape-related > space-related speech-gesture pairs | MNI | 10 | high |
| 15 | Neudorf et al. (2019) | fMRI | 25 | mental imagery, lexical decision | written words | (target > pseudohomophone) for (object > action priming) | MNI | 6 | high |
| 16 | Noppeney et al. (2002) | PET | 12 | semantic decision | spoken words | visual words in semantic decision > perceptual decision | TAL | 1 | low |
| 17 | Phillips et al. (2002) | PET | 12 | semantic decision | pictures, written words | (size > action retrieval) ∩ (size retrieval > screen size control) | MNI | 1 | high |
| 18 | Pulvermueller et al. (2006) | fMRI | 14 | passive reading | written words | form words > hash marks | TAL | 12 | low |
| 19 | Thompson-Schill et al. (1999) | fMRI | 5 | semantic decision | spoken sentences | visual semantics (living) > reversed words | TAL | 5 | high |
| 19 | Thompson-Schill et al. (1999) | fMRI | 5 | semantic decision | spoken sentences | visual semantics (non-living) > reversed words | TAL | 4 | high |
| 20 | Wheatley et al. (2005) | fMRI | 15 | passive reading | written words | animate objects > tools | TAL | 26 | high |
| 21 | Wheeler et al. (2000) | fMRI | 18 | episodic memory recall | written words | picture recall > sound recall | TAL | 13 | high |
| 22 | Zvyagintsev et al. (2013) | fMRI | 15 | mental imagery | rest | visual imagery > backwards counting | TAL | 11 | high |
| 22 | Zvyagintsev et al. (2013) | fMRI | 15 | mental imagery | rest | visual imagery > auditory imagery | TAL | 8 | high |

**Table S34. Experiments included in the ALE meta-analysis on conceptual processing related to motion.**

| Exp. | Paper | Method | Participants (N) | Task | Stimulus Modality | Contrast | Space | Foci (N) | Contrast Type |
|---|---|---|---|---|---|---|---|---|---|
| 1 | Bedny et al. (2012) | fMRI | 21 | semantic decision | spoken words | high-motion verbs > low-motion verbs | MNI | 5 | high |
| 1 | Bedny et al. (2012) | fMRI | 21 | semantic decision | spoken words | high-motion verbs > low-motion nouns | MNI | 2 | high |
| 2 | Bedny et al. (2014) | fMRI | 18 | semantic decision | spoken words | event nouns > plant nouns | MNI | 4 | high |
| 2 | Bedny et al. (2014) | fMRI | 18 | semantic decision | spoken words | motion verbs > animal nouns | MNI | 1 | high |
| 3 | Borghesani et al. (2019) | fMRI | 13 | semantic decision | pictures | movement task > place task | MNI | 4 | high |
| 4 | Chen et al. (2008) | fMRI | 14 | semantic decision | written sentences | literal motion sentences > non-motion sentences | TAL | 6 | high |
| 4 | Chen et al. (2008) | fMRI | 14 | semantic decision | written sentences | metaphor motion sentences > non-motion sentences | TAL | 1 | high |
| 5 | Deen & McCarthy (2010) | fMRI | 15 | passive reading | written sentences | motion sentences > non-motion sentences | MNI | 13 | high |
| 6 | Dravida et al. (2013) | fMRI | 18 | semantic decision | written narratives, spoken words | high motion passages > low motion passages | MNI | 19 | high |
| 6 | Dravida et al. (2013) | fMRI | 18 | semantic decision | written narratives, spoken words | high motion nouns > low motion nouns | MNI | 5 | high |
| 6 | Dravida et al. (2013) | fMRI | 18 | semantic decision | written narratives, spoken words | high motion verbs > low motion verbs | MNI | 3 | high |
| 7 | Fernandino et al. (2016) | fMRI | 44 | semantic decision | written words | visual motion > other attributes | TAL | 1 | high |
| 7 | Fernandino et al. (2016) | fMRI | 44 | semantic decision | written words | motion > other attributes except sound | TAL | 4 | high |
| 8 | Grossman et al. (2002b) | fMRI | 16 | semantic decision | written words | motion verbs > cognition verbs | TAL | 3 | high |
| 9 | Humphreys et al. (2013) | fMRI | 10 | semantic decision | pictures, spoken sentences | motion sentences > static sentences | MNI | 1 | high |
| 10 | Kemmerer et al. (2008) | fMRI | 16 | semantic decision | written words | running verbs > meaningless symbols | MNI | 23 | low |
| 10 | Kemmerer et al. (2008) | fMRI | 16 | semantic decision | written words | speaking verbs > meaningless symbols | MNI | 8 | low |
| 10 | Kemmerer et al. (2008) | fMRI | 16 | semantic decision | written words | change of state verbs > meaningless symbols | MNI | 9 | low |
| 11 | Lai & Desai (2016) | fMRI | 22 | semantic decision | written sentences | temporal fictive motion sentences > temporal, static sentences | TAL | 5 | high |

| Exp. | Paper | Method | Participants (N) | Task | Stimulus Modality | Contrast | Space | Foci (N) | Contrast Type |
|---|---|---|---|---|---|---|---|---|---|
| 12 | Lauro et al. (2013) | fMRI | 24 | semantic decision | written sentences | idiomatic motion sentences involving lower-limb > cognition sentences | MNI | 4 | high |
| 12 | Lauro et al. (2013) | fMRI | 24 | semantic decision | written sentences | metaphorical motion sentences involving lower-limb > cognition sentences | MNI | 1 | high |
| 13 | Lin et al. (2015) | fMRI | 17 | semantic decision | written words | social interaction verbs > non-human verbs | TAL | 6 | high |
| 13 | Lin et al. (2015) | fMRI | 17 | semantic decision | written words | motion verbs > non-human verbs | TAL | 5 | high |
| 14 | Nagels et al. (2013) | fMRI | 17 | perceptual decision | videos, spoken sentences | space-related > shape-related speech-gesture pairs | MNI | 8 | high |
| 15 | Noppeney et al. (2005) | fMRI | 12 | semantic decision | spoken words | body movement words > hand action words | TAL | 1 | high |
| 16 | Peelen et al. (2012) | fMRI | 27 | working memory recall | written words | action verbs > object nouns | TAL | 4 | high |
| 17 | Rodriguez-Ferreiro et al. (2011) | fMRI | 14 | passive reading | written words | motion verbs > emotion verbs | MNI | 3 | high |
| 17 | Rodriguez-Ferreiro et al. (2011) | fMRI | 14 | passive reading | written words | transitive motion verbs > emotion verbs | MNI | 2 | high |
| 17 | Rodriguez-Ferreiro et al. (2011) | fMRI | 14 | passive reading | written words | motion verbs > pseudoverbs | MNI | 7 | high |
| 18 | Schuil et al. (2013) | fMRI | 20 | lexical decision | written sentences | literal > non-literal hand- and foot-related sentences | MNI | 2 | high |
| 18 | Schuil et al. (2013) | fMRI | 20 | lexical decision | written sentences | literal hand- and foot-related sentences > non-word sentences | MNI | 6 | high |
| 18 | Schuil et al. (2013) | fMRI | 20 | lexical decision | written sentences | non-literal hand- and foot-related sentences > non-word sentences | MNI | 4 | high |
| 19 | Speer et al. (2009) | fMRI | 28 | passive reading | written narratives | location change > clause onset | MNI | 12 | high |
| 20 | van Dam & Desai (2016) | fMRI | 14 | semantic decision | written sentences | concrete motion sentences > abstract sentences | TAL | 4 | high |
| 20 | van Dam & Desai (2016) | fMRI | 14 | semantic decision | written sentences | concrete caused motion > abstract caused motion | TAL | 4 | high |
| 20 | van Dam & Desai (2016) | fMRI | 14 | semantic decision | written sentences | concrete intransitive motion > abstract intransitive sentences | TAL | 4 | high |
| 21 | van Dam et al. (2019) | fMRI | 17 | semantic decision | pictures | manner pictures > action pictures | TAL | 5 | high |
| 21 | van Dam et al. (2019) | fMRI | 17 | semantic decision | pictures | manner pictures > scrambled manner pictures | TAL | 3 | high |
| 21 | van Dam et al. (2019) | fMRI | 17 | lexical decision, go/no-go | written words | manner verbs > pseudoverbs | TAL | 5 | high |
| 22 | Vigliocco et al. (2006) | PET | 12 | passive listening | spoken words | motion words > sensory words | MNI | 2 | high |

| Exp. | Paper | Method | Participants (N) | Task | Stimulus Modality | Contrast | Space | Foci (N) | Contrast Type |
|---|---|---|---|---|---|---|---|---|---|
| 23 | Wallentin et al. (2005) | fMRI | 15 | semantic decision | spoken sentences | motion sentences > static sentences | MNI | 5 | high |
| 24 | Wallentin et al. (2011) | fMRI | 26 | passive listening | spoken narratives | motion verbs > rest of story | MNI | 2 | high |

**Table S35. Experiments included in the ALE meta-analysis on conceptual processing related to color.**

| Exp. | Paper | Method | Participants (N) | Task | Stimulus Modality | Contrast | Space | Foci (N) | Contrast Type |
|---|---|---|---|---|---|---|---|---|---|
| 1 | Chao & Martin (1999) | PET | 12 | word generation | pictures | color word generation > achromatic object naming | TAL | 16 | high |
| 1 | Chao & Martin (1999) | PET | 12 | word generation | pictures | color word generation > passive viewing | TAL | 20 | high |
| 1 | Chao & Martin (1999) | PET | 12 | word generation | pictures | color word generation > colored object naming | TAL | 6 | high |
| 2 | Fernandino et al. (2016) | fMRI | 44 | semantic decision | written words | color > other attributes | TAL | 6 | high |
| 2 | Fernandino et al. (2016) | fMRI | 44 | semantic decision | written words | color > other attributes except shape | TAL | 25 | high |
| 3 | Goldberg et al. (2006b) | fMRI | 14 | semantic decision | written words | color words > auditory, gustatory, tactile words | TAL | 3 | high |
| 4 | Howard et al. (1998) | fMRI | 7 | mental imagery | spoken sentences | color imagery > non-color visual imagery | TAL | 5 | high |
| 5 | Hsu et al. (2011) | fMRI | 12 | semantic decision | written words | (task > fixation) ∩ (within > between color comparisons) | TAL | 13 | high |
| 6 | Hsu et al. (2011) | fMRI | 12 | semantic decision | spoken words | (task > fixation) ∩ (within > between color comparisons) | MNI | 13 | high |
| 7 | Kellenbach et al. (2001) | PET | 10 | semantic decision | written words | color verifications > size verifications | MNI | 1 | high |
| 7 | Kellenbach et al. (2001) | PET | 10 | semantic decision | written words | color verifications > sound verifications | MNI | 3 | high |
| 7 | Kellenbach et al. (2001) | PET | 10 | semantic decision | written words | color verifications > letter-strings | MNI | 7 | low |
| 8 | Martin et al. (1995) | PET | 12 | word generation | pictures | color word generation > action word generation | TAL | 5 | high |
| 8 | Martin et al. (1995) | PET | 12 | word generation | written words | color word generation > action word generation | TAL | 4 | high |
| 8 | Martin et al. (1995) | PET | 12 | word generation | pictures | color word generation > object naming | TAL | 8 | high |
| 9 | Mummery et al. (1998) | PET | 10 | similarity judgment task | written words | color words > location words | TAL | 2 | high |

| Exp. | Paper | Method | Participants (N) | Task | Stimulus Modality | Contrast | Space | Foci (N) | Contrast Type |
|---|---|---|---|---|---|---|---|---|---|
| 10 | Pulvermueller et al. (2006) | fMRI | 14 | passive reading | written words | color words > hash marks | TAL | 3 | low |
| 11 | Slotnick (2009) | fMRI | 12 | episodic memory recall | pictures | color-hit-hit > color-hit-miss and grey-hit-hit | TAL | 1 | high |
| 12 | Wang et al. (2013) | fMRI | 48 | semantic decision | perceptual decision, semantic decision | correlation with color knowledge scores | MNI | 6 | high |
| 12 | Wang et al. (2013) | fMRI | 48 | semantic decision | perceptual decision, semantic decision | correlation with color knowledge scores (controlling for form, motion and sound scores) | MNI | 7 | high |

#### Table S36. Experiments included in the ALE meta-analysis on conceptual processing related to olfaction-gustation.

| Exp. | Paper | Method | Participants (N) | Task | Stimulus Modality | Contrast | Space | Foci (N) | Contrast Type |
|---|---|---|---|---|---|---|---|---|---|
| 1 | Barros-Loscertales et al. (2011) | fMRI | 59 | passive reading | written words | taste-related words > taste-unrelated words | MNI | 11 | high |
| 2 | Cerf-Ducastel & Murphy (2006) | fMRI | 10 | working memory recall | written words | known > unknown odor names | TAL | 21 | high |
| 2 | Cerf-Ducastel & Murphy (2006) | fMRI | 10 | working memory recall | written words | known > unknown odor names | TAL | 26 | high |
| 3 | Citron & Goldberg (2014) | fMRI | 26 | semantic decision | written sentences | taste-related metaphors > literal, not taste-related counterparts | MNI | 19 | high |
| 4 | Dreyer et al. (2018) | fMRI | 28 | passive reading | written words | food words > hash marks | MNI | 4 | low |
| 5 | Fairhall (2020) | fMRI | 16 | semantic decision | written words | food words > people and place words | MNI | 3 | high |
| 6 | Fournel et al. (2017) | fMRI | 19 | passive smelling, working memory recall | odors | associative learning > perceptual learning | MNI | 5 | high |
| 6 | Fournel et al. (2017) | fMRI | 19 | passive smelling, working memory recall | odors | previously learned odor > unknown odor | MNI | 5 | high |
| 7 | Ghio et al. (2016a) | fMRI | 16 | episodic memory recall | pictures | olfactory > visual | MNI | 1 | high |
| 8 | Goldberg et al. (2006a) | fMRI | 13 | word generation, semantic decision | written words | fruit words > bird, body part, clothing words | TAL | 5 | high |
| 9 | Goldberg et al. (2006b) | fMRI | 14 | semantic decision | written words | gustatory words > auditory, tactile, visual words | TAL | 1 | high |

| Exp. | Paper | Method | Participants (N) | Task | Stimulus Modality | Contrast | Space | Foci (N) | Contrast Type |
|---|---|---|---|---|---|---|---|---|---|
| 10 | Gonzalez et al. (2006) | fMRI | 23 | passive reading | written words | olfactory words > non-olfactory words | MNI | 8 | high |
| 11 | Gottfried & Dolan (2003) | fMRI | 17 | perceptual decision | odors | olfactory-visual > olfaction + vision | MNI | 5 | high |
| 11 | Gottfried & Dolan (2003) | fMRI | 17 | perceptual decision | odors | olfactory-visual congruent > olfaction + vision | MNI | 4 | high |
| 11 | Gottfried & Dolan (2003) | fMRI | 17 | perceptual decision | odors | olfactory-visual congruent > incongruent | MNI | 3 | high |
| 11 | Gottfried & Dolan (2003) | fMRI | 17 | perceptual decision | odors | correlation with olfactory-visual congruency | MNI | 3 | high |
| 12 | Just et al. (2010) | fMRI | 11 | mental imagery | written words | eating factor | MNI | 3 | low |
| 13 | Kobayashi et al. (2004) | fMRI | 25 | mental imagery | written words | gustatory imagery > passive reading | MNI | 6 | high |
| 13 | Kobayashi et al. (2004) | fMRI | 25 | mental imagery | written words | gustatory imagery > visual imagery | MNI | 3 | high |
| 13 | Kobayashi et al. (2004) | fMRI | 25 | mental imagery | written words | gustatory imagery > visual imagery | MNI | 1 | high |
| 13 | Kobayashi et al. (2004) | fMRI | 25 | mental imagery | written words | gustatory imagery > passive observation | MNI | 6 | high |
| 13 | Kobayashi et al. (2004) | fMRI | 25 | mental imagery | written words | gustatory imagery > rest | MNI | 7 | high |
| 14 | Phillips et al. (2002) | PET | 12 | semantic decision | pictures, written words | fruits > tools | MNI | 1 | high |
| 15 | Pomp et al. (2018) | fMRI | 18 | semantic decision | written sentences | olfactory metaphors > literal paraphrases | MNI | 2 | high |
| 15 | Pomp et al. (2018) | fMRI | 18 | semantic decision | written sentences | literal olfactory sentences > literal paraphrases | MNI | 2 | high |
| 15 | Pomp et al. (2018) | fMRI | 18 | semantic decision | written sentences | literal paraphrases > literal olfactory sentences | MNI | 2 | high |
| 15 | Pomp et al. (2018) | fMRI | 18 | semantic decision | written sentences | literal olfactory sentences > olfactory metaphors | MNI | 1 | high |
| 16 | Savic & Berglund (2004) | PET | 14 | passive smelling | odors | familiar odor > unfamiliar odor | TAL | 3 | high |
| 17 | Simmons et al. (2005) | fMRI | 9 | working memory recall | pictures | food pictures > location pictures | MNI | 6 | high |

**Table S37. Experiments included in the ALE meta-analysis on conceptual processing related to emotion.**

| Exp. | Paper | Method | Participants (N) | Task | Stimulus Modality | Contrast | Space | Foci (N) | Contrast Type |
|---|---|---|---|---|---|---|---|---|---|
| 1 | Altenmueller et al. (2014) | fMRI | 18 | episodic memory recall | music | positive valence > negative valence | TAL | 9 | high |
| 2 | Berthoz et al. (2002) | fMRI | 12 | mental imagery | written narratives | (intentional and unintentional embarrassment) > normal story | TAL | 18 | high |
| 2 | Berthoz et al. (2002) | fMRI | 12 | mental imagery | written narratives | unintentional embarrassment > normal story | TAL | 15 | high |
| 3 | Bogert et al. (2016) | fMRI | 56 | semantic judgment | music | main effect of emotion | MNI | 7 | high |
| 3 | Bogert et al. (2016) | fMRI | 56 | semantic judgment | music | explicit task > implicit task | MNI | 4 | high |
| 4 | Chow et al. (2014) | fMRI | 24 | semantic decision | spoken narratives | emotion paragraphs > action and perception paragraphs | MNI | 16 | high |
| 5 | Crosson et al. (1999) | fMRI | 17 | word generation | spoken words | emotionally connotated words > neutral words | TAL | 2 | high |
| 5 | Crosson et al. (1999) | fMRI | 17 | word generation | spoken words | emotionally evocative words > repetition | TAL | 12 | high |
| 5 | Crosson et al. (1999) | fMRI | 17 | word generation | spoken words | emotionally neutral words > repetition | TAL | 7 | high |
| 6 | Cunningham et al. (2003) | fMRI | 12 | semantic decision | written words | evaluative judgments > non-evaluative judgments | MNI | 3 | high |
| 6 | Cunningham et al. (2003) | fMRI | 12 | semantic decision | written words | correlation of ambivalence score for evaluative judgments | MNI | 1 | high |
| 6 | Cunningham et al. (2003) | fMRI | 12 | semantic decision | written words | bad judgments > good judgments | MNI | 2 | high |
| 7 | Cunningham et al. (2004) | fMRI | 20 | semantic decision | written words | (good > bad) > (abstract > concrete) | MNI | 14 | high |
| 7 | Cunningham et al. (2004) | fMRI | 20 | semantic decision | written words | emotional intensity: good > bad ∩ abstract > concrete | MNI | 13 | high |
| 7 | Cunningham et al. (2004) | fMRI | 20 | semantic decision | written words | emotional intensity: (good > bad) > (abstract > concrete) | MNI | 3 | high |
| 7 | Cunningham et al. (2004) | fMRI | 20 | semantic decision | written words | emotional intensity: good > bad | MNI | 7 | high |
| 7 | Cunningham et al. (2004) | fMRI | 20 | semantic decision | written words | emotional valence: good > bad ∩ abstract > concrete | MNI | 3 | high |
| 7 | Cunningham et al. (2004) | fMRI | 20 | semantic decision | written words | emotional valence: good > bad | MNI | 1 | high |
| 8 | Dietrich et al. (2008) | fMRI | 16 | passive listening | spoken words | affective high-lexical interjections > rest | MNI | 2 | low |
| 8 | Dietrich et al. (2008) | fMRI | 16 | passive listening | spoken words | affective low-lexical interjections > rest | MNI | 2 | low |
| 8 | Dietrich et al. (2008) | fMRI | 16 | passive listening | spoken words | affective high and low lexical interjections > rest | MNI | 4 | low |
| 9 | Dreyer et al. (2018) | fMRI | 28 | passive reading | written words | abstract emotional words > hash marks | MNI | 5 | low |
| 10 | Ferstl & von Cramon (2007) | fMRI | 20 | semantic decision | written narratives | emotion stories > time and space stories | TAL | 1 | high |

| Exp. | Paper | Method | Participants (N) | Task | Stimulus Modality | Contrast | Space | Foci (N) | Contrast Type |
|---|---|---|---|---|---|---|---|---|---|
| 11 | Ferstl et al. (2005) | fMRI | 20 | semantic decision | spoken narratives | emotional stories > chronological stories | TAL | 3 | high |
| 11 | Ferstl et al. (2005) | fMRI | 20 | semantic decision | spoken narratives | emotional stories: inconsistent > consistent | TAL | 3 | high |
| 12 | Ghio et al. (2016b) | fMRI | 50 | passive viewing | pictures | emotion localizer task > rest | MNI | 38 | low |
| 13 | Goel & Dolan (2003) | fMRI | 19 | semantic decision | written sentences | emotional reasoning > neutral reasoning | MNI | 2 | high |
| 14 | Hellbernd & Sammler (2018) | fMRI | 22 | semantic decision | sounds | social function: clear > ambiguous | MNI | 24 | high |
| 15 | Herz et al. (2004) | fMRI | 5 | mental imagery | pictures, odors | memory-linked odors > photographs | TAL | 1 | high |
| 15 | Herz et al. (2004) | fMRI | 5 | mental imagery | pictures, odors | memory-linked odors > neutral odors and photographs | TAL | 1 | high |
| 15 | Herz et al. (2004) | fMRI | 5 | mental imagery | pictures, odors | memory-linked odors > control odors | TAL | 6 | high |
| 16 | Hofstetter et al. (2012) | fMRI | 18 | episodic memory recall | pictures, written words | negative association > neutral association | MNI | 10 | high |
| 17 | Isenberg et al. (1999) | PET | 6 | perceptual decision | written words | threat words > neutral words | TAL | 4 | high |
| 18 | Jabbi et al. (2007) | fMRI | 18 | semantic judgment | pictures, beverages | (disgusting taste and disgusted faces) > (neutral taste and neutral faces) | MNI | 5 | high |
| 18 | Jabbi et al. (2007) | fMRI | 18 | semantic judgment | pictures, beverages | (delicious taste and satisfied faces) > (neutral taste and neutral faces) | MNI | 2 | high |
| 19 | Kedia et al. (2008) | fMRI | 29 | mental imagery | written sentences | emotional > neutral | TAL | 5 | high |
| 19 | Kedia et al. (2008) | fMRI | 29 | mental imagery | written sentences | anger for other and compassion > anger for self and guilt | TAL | 3 | high |
| 19 | Kedia et al. (2008) | fMRI | 29 | mental imagery | written sentences | (guilt > anger for other) > (anger for self > compassion) | TAL | 12 | high |
| 20 | Knutson et al. (2006) | fMRI | 30 | semantic decision | pictures | attitude-congruent > attitude-incongruent | TAL | 2 | high |
| 20 | Knutson et al. (2006) | fMRI | 30 | semantic decision | pictures | attitude-congruent > perceptual decision | TAL | 4 | high |
| 21 | Kuchinke et al. (2005) | fMRI | 20 | lexical decision | written words | emotional words > neutral words | MNI | 3 | high |
| 21 | Kuchinke et al. (2005) | fMRI | 20 | lexical decision | written words | negative words > neutral words | MNI | 1 | high |
| 21 | Kuchinke et al. (2005) | fMRI | 20 | lexical decision | written words | positive words > neutral words | MNI | 4 | high |
| 21 | Kuchinke et al. (2005) | fMRI | 20 | lexical decision | written words | positive words > negative words | MNI | 4 | high |

| Exp. | Paper | Method | Participants (N) | Task | Stimulus Modality | Contrast | Space | Foci (N) | Contrast Type |
|---|---|---|---|---|---|---|---|---|---|
| 22 | Luo et al. (2004) | fMRI | 9 | perceptual decision | written words | prime identical to negative target > unrelated prime | TAL | 3 | high |
| 22 | Luo et al. (2004) | fMRI | 9 | perceptual decision | written words | prime identical to positive target > unrelated prime | TAL | 2 | high |
| 23 | Maddock et al. (2003) | fMRI | 8 | semantic decision | spoken words | unpleasant words > neutral words | TAL | 28 | high |
| 23 | Maddock et al. (2003) | fMRI | 8 | semantic decision | spoken words | pleasant words > neutral words | TAL | 16 | high |
| 24 | Maratos et al. (2001) | fMRI | 12 | semantic judgment, working memory recall | written sentences | negative sentences > neutral sentences | TAL | 11 | high |
| 24 | Maratos et al. (2001) | fMRI | 12 | semantic judgment, working memory recall | written sentences | negative sentences > neutral sentences | TAL | 8 | high |
| 24 | Maratos et al. (2001) | fMRI | 12 | semantic judgment, working memory recall | written sentences | positive sentences > neutral sentences | TAL | 10 | high |
| 24 | Maratos et al. (2001) | fMRI | 12 | semantic judgment, working memory recall | written sentences | positive sentences > neutral sentences | TAL | 2 | high |
| 25 | Michl et al. (2014) | fMRI | 14 | mental imagery | written sentences | shame > neutral | TAL | 10 | high |
| 25 | Michl et al. (2014) | fMRI | 14 | mental imagery | written sentences | guilt > neutral | TAL | 19 | high |
| 25 | Michl et al. (2014) | fMRI | 14 | mental imagery | written sentences | shame > guilt | TAL | 7 | high |
| 25 | Michl et al. (2014) | fMRI | 14 | mental imagery | written sentences | guilt > shame | TAL | 3 | high |
| 26 | Moll et al. (2007) | fMRI | 12 | mental imagery | written sentences | guilt > neutral | TAL | 6 | high |
| 26 | Moll et al. (2007) | fMRI | 12 | mental imagery | written sentences | embarrassment > neutral | TAL | 5 | high |
| 26 | Moll et al. (2007) | fMRI | 12 | mental imagery | written sentences | compassion > neutral | TAL | 11 | high |
| 26 | Moll et al. (2007) | fMRI | 12 | mental imagery | written sentences | disgust > neutral | TAL | 9 | high |
| 26 | Moll et al. (2007) | fMRI | 12 | mental imagery | written sentences | indignation for self > neutral | TAL | 4 | high |
| 26 | Moll et al. (2007) | fMRI | 12 | mental imagery | written sentences | indignation for other > neutral | TAL | 5 | high |
| 27 | Moseley et al. (2012) | fMRI | 18 | passive reading | written words | emotion words > hash marks | MNI | 16 | low |
| 27 | Moseley et al. (2012) | fMRI | 18 | passive reading | written words | abstract emotion words > hash marks | MNI | 16 | low |
| 28 | Nakic et al. (2006) | fMRI | 13 | lexical decision | written words | negative words > neutral words | TAL | 7 | high |
| 29 | Oosterwick et al. (2015) | fMRI | 18 | semantic decision | written sentences | emotion sentences with internal focus > emotion sentences with external focus | TAL | 1 | high |
| 29 | Oosterwick et al. (2015) | fMRI | 18 | semantic decision | written sentences | emotion sentences with internal focus > non-emotion sentences with internal focus | TAL | 2 | high |
| 29 | Oosterwick et al. (2015) | fMRI | 18 | semantic decision | written sentences | emotion sentences with internal focus > non-emotion sentences with external focus | TAL | 1 | high |

| Exp. | Paper | Method | Participants (N) | Task | Stimulus Modality | Contrast | Space | Foci (N) | Contrast Type |
|---|---|---|---|---|---|---|---|---|---|
| 30 | Phillips et al. (1998) | fMRI | 6 | perceptual decision | pictures | disgust > neutral | TAL | 18 | high |
| 30 | Phillips et al. (1998) | fMRI | 6 | perceptual decision | pictures | fear > neutral | TAL | 12 | high |
| 30 | Phillips et al. (1998) | fMRI | 6 | perceptual decision | pictures | fear > disgust | TAL | 10 | high |
| 30 | Phillips et al. (1998) | fMRI | 6 | perceptual decision | pictures | disgust > fear | TAL | 9 | high |
| 30 | Phillips et al. (1998) | fMRI | 6 | perceptual decision | sounds | disgust > neutral | TAL | 14 | high |
| 30 | Phillips et al. (1998) | fMRI | 6 | perceptual decision | sounds | fear > neutral | TAL | 14 | high |
| 30 | Phillips et al. (1998) | fMRI | 6 | perceptual decision | sounds | fear > disgust | TAL | 7 | high |
| 30 | Phillips et al. (1998) | fMRI | 6 | perceptual decision | sounds | disgust > fear | TAL | 8 | high |
| 31 | Rodriguez-Ferreiro et al. (2011) | fMRI | 14 | passive reading | written words | transitive emotion verbs > motion verbs | MNI | 5 | high |
| 32 | Ruby & Decety (2004) | PET | 10 | mental imagery | written sentences | emotional sentences > neutral sentences | MNI | 7 | high |
| 32 | Ruby & Decety (2004) | PET | 10 | mental imagery | written sentences | emotional > neutral sentences for 3rd > 1st perspective | MNI | 1 | high |
| 32 | Ruby & Decety (2004) | PET | 10 | mental imagery | written sentences | emotional > neutral sentences for 1st > 3rd perspective | MNI | 1 | high |
| 33 | Saxbe et al. (2013) | fMRI | 28 | semantic judgment | videos | correlation affective word use and BOLD activity for emotion trials > fixation | MNI | 32 | high |
| 34 | Skipper & Olson (2014) | fMRI | 19 | semantic decision, lexical decision | written words | valenced words > neutral words | MNI | 1 | high |
| 34 | Skipper & Olson (2014) | fMRI | 19 | semantic decision, lexical decision | written words | valenced words > pseudowords | MNI | 18 | high |
| 35 | Skipper et al. (2011) | fMRI | 18 | semantic decision | pictures, sounds | social > non-social | TAL | 11 | high |
| 36 | Smith et al. (2004) | fMRI | 15 | working memory recall | pictures | emotional hits > neutral hits | MNI | 11 | high |
| 36 | Smith et al. (2004) | fMRI | 15 | working memory recall | pictures | negative hits > positive hits | MNI | 8 | high |
| 36 | Smith et al. (2004) | fMRI | 15 | working memory recall | pictures | positive hits > negative hits | MNI | 2 | high |
| 36 | Smith et al. (2004) | fMRI | 15 | working memory recall | pictures | negative hits > neutral hits | MNI | 5 | high |
| 36 | Smith et al. (2004) | fMRI | 15 | working memory recall | pictures | positive hits > neutral hits | MNI | 2 | high |
| 36 | Smith et al. (2004) | fMRI | 15 | working memory recall | pictures | encoding objects: emotional context > neutral context | MNI | 12 | high |
| 37 | Steinbeis & Koelsch (2008) | fMRI | 16 | semantic decision | sounds, written words | incongruent > congruent pleasantness of chords and target words | TAL | 2 | high |
| 38 | Strange et al. (2000) | fMRI | 11 | perceptual decision, semantic decision | written words | emotional oddball words > neutral words | TAL | 2 | high |

| Exp. | Paper | Method | Participants (N) | Task | Stimulus Modality | Contrast | Space | Foci (N) | Contrast Type |
|---|---|---|---|---|---|---|---|---|---|
| 39 | Takahashi et al. (2004) | fMRI | 19 | passive reading, semantic decision | written sentences | guilt > neutral | TAL | 5 | high |
| 39 | Takahashi et al. (2004) | fMRI | 19 | passive reading, semantic decision | written sentences | embarrassment > neutral | TAL | 10 | high |
| 39 | Takahashi et al. (2004) | fMRI | 19 | passive reading, semantic decision | written sentences | guilt, embarrassment > neutral | TAL | 5 | high |
| 39 | Takahashi et al. (2004) | fMRI | 19 | passive reading, semantic decision | written sentences | embarrassment > guilt | TAL | 7 | high |
| 39 | Takahashi et al. (2004) | fMRI | 19 | passive reading, semantic decision | written sentences | guilt > embarrassment | TAL | 1 | high |
| 40 | Tavares et al. (2011) | fMRI | 16 | semantic decision | animations | interacting animations > neutral animations | TAL | 15 | high |
| 41 | Tettamanti et al. (2012) | fMRI | 19 | passive viewing | videos | emotional > neutral videos | MNI | 13 | high |
| 41 | Tettamanti et al. (2012) | fMRI | 19 | passive viewing | videos | fear > neutral videos | MNI | 12 | high |
| 41 | Tettamanti et al. (2012) | fMRI | 19 | passive viewing | videos | disgust > neutral videos | MNI | 48 | high |
| 41 | Tettamanti et al. (2012) | fMRI | 19 | passive viewing | videos | happy > neutral videos | MNI | 4 | high |
| 42 | Vigliocco et al. (2014) | fMRI | 20 | lexical decision | written words | correlation with emotional valence for all words | MNI | 9 | high |
| 42 | Vigliocco et al. (2014) | fMRI | 20 | lexical decision | written words | correlation with emotional valence for abstract > concrete words | MNI | 1 | high |
| 42 | Vigliocco et al. (2014) | fMRI | 20 | lexical decision | written words | correlation with emotional valence for concrete words | MNI | 2 | high |
| 43 | Wang et al. (2019) | fMRI | 23 | semantic decision | written words | positive valence > neutral valence | MNI | 4 | high |
| 43 | Wang et al. (2019) | fMRI | 23 | semantic decision | written words | social > non-social for positive > neutral valence | MNI | 10 | high |
| 44 | Wicker et al. (2003) | fMRI | 14 | passive viewing | videos | disgusted > neutral faces | MNI | 17 | high |
| 44 | Wicker et al. (2003) | fMRI | 14 | passive viewing | videos | pleasured > neutral faces | MNI | 6 | high |
| 44 | Wicker et al. (2003) | fMRI | 14 | passive viewing | videos | disgusted faces ∩ disgusting odor | MNI | 4 | high |
| 45 | Wilson-Mendenhall et al. (2011) | fMRI | 20 | mental imagery | spoken sentences | anger > fear, observe, plan | TAL | 2 | high |
| 45 | Wilson-Mendenhall et al. (2011) | fMRI | 20 | mental imagery | spoken sentences | anger, fear, plan > observe | TAL | 7 | high |
| 46 | Zysset et al. (2002) | fMRI | 13 | semantic decision | written sentences | evaluative judgments > semantic judgments | TAL | 17 | high |
| 47 | Zysset et al. (2003) | fMRI | 18 | semantic decision | written sentences | evaluative judgments > semantic judgments | TAL | 10 | high |

#### References

Altenmüller, E., Siggel, S., Mohammadi, B., Samii, A., Münte, T.F., 2014. Play it again, Sam: Brain correlates of emotional music recognition. Front. Psychol. 5, 1–8. https://doi.org/10.3389/fpsyg.2014.00114

Barrós-Loscertales, A., González, J., Pulvermüller, F., Ventura-Campos, N., Bustamante, J.C., Costumero, V., Parcet, M.A., Ávila, C., 2012. Reading salt activates gustatory brain regions: FMRI evidence for semantic grounding in a novel sensory modality. Cereb. Cortex 22, 2554–2563. https://doi.org/10.1093/cercor/bhr324

Baumgaertner, A., Buccino, G., Lange, R., McNamara, A., Binkofski, F., 2007. Polymodal conceptual processing of human biological actions in the left inferior frontal lobe. Eur. J. Neurosci. 25, 881–889. https://doi.org/10.1111/j.1460-9568.2007.05346.x

Bedny, M., Caramazza, A., Grossman, E., Pascual-Leone, A., Saxe, R., 2008. Concepts are more than percepts: the case of action verbs. J Neurosci 28, 11347–11353. https://doi.org/10.1523/JNEUROSCI.3039-08.2008

Bedny, M., Caramazza, A., Pascual-Leone, A., Saxe, R., 2012. Typical neural representations of action verbs develop without vision. Cereb. Cortex 22, 286–293. https://doi.org/10.1093/cercor/bhr081

Bedny, M., Dravida, S., Saxe, R., 2014. Shindigs, brunches, and rodeos: The neural basis of event words. Cogn. Affect. Behav. Neurosci. 14, 891–901. https://doi.org/10.3758/s13415-013-0217-z

Bellebaum, C., Tettamanti, M., Marchetta, E., Della Rosa, P., Rizzo, G., Daum, I., Cappa, S.F., 2013. Neural representations of unfamiliar objects are modulated by sensorimotor experience. Cortex 49, 1110–1125. https://doi.org/10.1016/j.cortex.2012.03.023

Berthoz, S., Armony, J.L., Blair, R.J.R., Dolan, R.J., 2002. An fMRI study of intentional and unintentional (embarrassing) violations of social norms. Brain 125, 1696–1708. https://doi.org/10.1093/brain/awf190

Bogert, B., Numminen-Kontti, T., Gold, B., Sams, M., Numminen, J., Burunat, I., Lampinen, J., Brattico, E., 2016. Hidden sources of joy, fear, and sadness: Explicit versus implicit neural processing of musical emotions. Neuropsychologia 89, 393–402. https://doi.org/10.1016/j.neuropsychologia.2016.07.005

Bonner, M.F., Peelle, J.E., Cook, P.A., Grossman, M., 2013. Heteromodal conceptual processing in the angular gyrus. Neuroimage 71, 175–186. https://doi.org/10.1016/j.neuroimage.2013.01.006

Borghesani, V., Riello, M., Gesierich, B., Brentari, V., Monti, A., Gorno-Tempini, M.L., 2019. The Neural Representations of Movement across Semantic Categories. J. Cogn. Neurosci. 31, 791–807. https://doi.org/10.1162/jocn_a_01390

Boronat, C.B., Buxbaum, L.J., Coslett, H.B., Tang, K., Saffran, E.M., Kimberg, D.Y., Detre, J.A., 2005. Distinctions between manipulation and function knowledge of objects: evidence from functional magnetic resonance imaging. Cogn. Brain Res. 23, 361–373. https://doi.org/10.1016/j.cogbrainres.2004.11.001

Boulenger, V., Hauk, O., Pulvermuller, F., 2009. Grasping Ideas with the Motor System: Semantic Somatotopy in Idiom Comprehension. Cereb. Cortex 19, 1905–1914. https://doi.org/10.1093/cercor/bhn217

Bracci, S., Peelen, M. V., 2013. Body and object effectors: The organization of object representations in high-level visual cortex reflects body-object interactions. J. Neurosci. 33, 18247–18258. https://doi.org/10.1523/JNEUROSCI.1322-13.2013

Cappa, S.F., Perani, D., Schnur, T., Tettamanti, M., Fazio, F., 1998. The effects of semantic category and knowledge type on lexical-semantic access: A PET Study. Neuroimage 8, 350–359.

https://doi.org/10.1006/nimg.1998.0368

Cerf-Ducastel, B., Murphy, C., 2006. Neural substrates of cross-modal olfactory recognition memory: An fMRI study. Neuroimage 31, 386–396. https://doi.org/10.1016/j.neuroimage.2005.11.009

Chao, L.L., Martin, A., 2000. Representation of manipulable man-made objects in the dorsal stream. Neuroimage 12, 478–84. https://doi.org/10.1006/nimg.2000.0635

Chao, L.L., Martin, A., 1999. Cortical regions associated with perceiving, naming, and knowing about colors. J. Cogn. Neurosci. 11, 25–35. https://doi.org/10.1162/089892999563229

Chen, E., Widick, P., Chatterjee, A., 2008. Functional-anatomical organization of predicate metaphor processing. Brain Lang. 107, 194–202. https://doi.org/10.1016/j.bandl.2008.06.007

Chow, H.M., Mar, R.A., Xu, Y., Liu, S., Wagage, S., Braun, A.R., 2014. Embodied Comprehension of Stories: Interactions between Language Regions and Modality-specific Neural Systems. J. Cogn. Neurosci. 26, 279–295. https://doi.org/10.1162/jocn_a_00487

Citron, F.M.M., Goldberg, A.E., 2014. Metaphorical Sentences Are More Emotionally Engaging than Their Literal Counterparts. J. Cogn. Neurosci. 26, 2585–2595. https://doi.org/10.1162/jocn_a_00654

Crosson, B., Radonovich, K., Sadek, J.R., Gökçay, D., Bauer, R.M., Fischler, I.S., Cato, M.A., Maron, L., Auerbach, E.J., Browd, S.R., Briggs, R.W., 1999. Left-hemisphere processing of emotional connotation during word generation. Neuroreport 10, 2449–2455. https://doi.org/10.1097/00001756-199908200-00003

Cunningham, W.A., Johnson, M.K., Gatenby, J.C., Gore, J.C., Banaji, M.R., 2003. Neural Components of Social Evaluation. J. Pers. Soc. Psychol. 85, 639–649. https://doi.org/10.1037/0022-3514.85.4.639

Cunningham, W.A., Raye, C.L., Johnson, M.K., 2004. Implicit and explicit evaluation: fMRI correlates of valence, emotional intensity, and control in the processing of attitudes. J. Cogn. Neurosci. 16, 1717–1729. https://doi.org/10.1162/0898929042947919

D'Esposito, M., Detre, J.A., Aguirre, G.K., Stallcup, M., Alsop, D.C., Tippet, L.J., Farah, M.J., 1997. A functional MRI study of mental image generation. Neuropsychologia 35, 725–730. https://doi.org/10.1016/S0028-3932(96)00121-2

Damasio, H., Grabowski, T.J., Tranel, D., Hichwa, R.D., Damasio, A.R., 1996. A neural basis for lexical retrieval. Nature. https://doi.org/10.1038/380499a0

Damasio, H., Grabowski, T.J., Tranel, D., Ponto, L.L.B., Hichwa, R.D., Damasio, A.R., 2001. Neural correlates of naming actions and of naming spatial relations. Neuroimage 13, 1053–1064. https://doi.org/10.1006/nimg.2001.0775

De Grauwe, S., Willems, R.M., Rueschemeyer, S.-A., Lemhöfer, K., Schriefers, H., 2014. Embodied language in first- and second-language speakers: Neural correlates of processing motor verbs. Neuropsychologia 56, 334–349. https://doi.org/10.1016/j.neuropsychologia.2014.02.003

Deen, B., McCarthy, G., 2010. Reading about the actions of others: Biological motion imagery and action congruency influence brain activity. Neuropsychologia 48, 1607–1615. https://doi.org/10.1016/j.neuropsychologia.2010.01.028

Desai, R.H., Binder, J.R., Conant, L.L., Mano, Q.R., Seidenberg, M.S., 2011. The neural career of sensorimotor metaphors. J. Cogn. Neurosci. 23, 2376–2386. https://doi.org/10.1162/jocn.2010.21596

Desai, R.H., Binder, J.R., Conant, L.L., Seidenberg, M.S., 2010. Activation of Sensory-Motor Areas in Sentence Comprehension. Cereb. Cortex 20, 468–478. https://doi.org/10.1093/cercor/bhp115

Desai, R.H., Choi, W., Lai, V.T., Henderson, J.M., 2016. Toward semantics in the wild: Activation to manipulable nouns in naturalistic reading. J. Neurosci. 36, 4050–4055.

https://doi.org/10.1523/JNEUROSCI.1480-15.2016

Desai, R.H., Conant, L.L., Binder, J.R., Park, H., Seidenberg, M.S., 2013. A piece of the action: Modulation of sensory-motor regions by action idioms and metaphors. Neuroimage 83, 862–869. https://doi.org/10.1016/j.neuroimage.2013.07.044

Dietrich, S., Hertrich, I., Alter, K., Ischebeck, A., Ackermann, H., 2008. Understanding the emotional expression of verbal interjections: A functional MRI study. Neuroreport 19, 1751–1755. https://doi.org/10.1097/WNR.0b013e3283193e9e

Dravida, S., Saxe, R., Bedny, M., 2013. People can understand descriptions of motion without activating visual motion brain regions. Front. Psychol. 4, 1–14. https://doi.org/10.3389/fpsyg.2013.00537

Dreyer, F.R., Pulvermüller, F., 2018a. Abstract semantics in the motor system? – An event-related fMRI study on passive reading of semantic word categories carrying abstract emotional and mental meaning. Cortex 100, 52–70. https://doi.org/10.1016/j.cortex.2017.10.021

Dreyer, F.R., Pulvermüller, F., 2018b. Abstract semantics in the motor system? – An event-related fMRI study on passive reading of semantic word categories carrying abstract emotional and mental meaning. Cortex 100, 52–70. https://doi.org/10.1016/j.cortex.2017.10.021

Fairhall, S.L., 2020. Cross recruitment of domain-selective cortical representations enables flexible semantic knowledge. J. Neurosci. 40, 3096–3103. https://doi.org/10.1523/JNEUROSCI.2224-19.2020

Fernandino, L., Binder, J.R., Desai, R.H., Pendl, S.L., Humphries, C.J., Gross, W.L., Conant, L.L., Seidenberg, M.S., 2016. Concept Representation Reflects Multimodal Abstraction: A Framework for Embodied Semantics. Cereb. Cortex 26, 2018–2034. https://doi.org/10.1093/cercor/bhv020

Ferstl, E.C., Rinck, M., Von Cramon, D.Y., 2005. Emotional and temporal aspects of situation model processing during text comprehension: An event-related fMRI study. J. Cogn. Neurosci. 17, 724–739. https://doi.org/10.1162/0898929053747658

Fournel, A., Sezille, C., Licon, C.C., Sinding, C., Gerber, J., Ferdenzi, C., Hummel, T., Bensafi, M., 2017. Learning to name smells increases activity in heteromodal semantic areas. Hum. Brain Mapp. 38, 5958–5969. https://doi.org/10.1002/hbm.23801

Gauvin, H.S., Jonen, M.K., Choi, J., McMahon, K., de Zubicaray, G.I., 2018. No lexical competition without priming: Evidence from the picture–word interference paradigm. Q. J. Exp. Psychol. 71, 2562–2570. https://doi.org/10.1177/1747021817747266

Gauvin, H.S., McMahon, K.L., Meinzer, M., de Zubicaray, G.I., 2019. The Shape of Things to Come in Speech Production: A Functional Magnetic Resonance Imaging Study of Visual Form Interference during Lexical Access. J. Cogn. Neurosci. 31, 913–921. https://doi.org/10.1162/jocn_a_01382

Ghio, M., Schulze, P., Suchan, B., Bellebaum, C., 2016a. Neural representations of novel objects associated with olfactory experience. Behav. Brain Res. 308, 143–151. https://doi.org/10.1016/j.bbr.2016.04.013

Ghio, M., Vaghi, M.M.S., Perani, D., Tettamanti, M., 2016b. Decoding the neural representation of fine-grained conceptual categories. Neuroimage 132, 93–103. https://doi.org/10.1016/j.neuroimage.2016.02.009

Gilead, M., Liberman, N., Maril, A., 2016. The effects of an action's "age-of-acquisition" on action-sentence processing. Neuroimage 141, 341–349. https://doi.org/10.1016/j.neuroimage.2016.07.034

Goel, V., Dolan, R.J., 2003. Reciprocal neural response within lateral and ventral medial prefrontal cortex during hot and cold reasoning. Neuroimage 20, 2314–2321. https://doi.org/10.1016/j.neuroimage.2003.07.027

Goldberg, R.F., Perfetti, C.A., Schneider, W., 2006a. Distinct and common cortical activations for multimodal semantic categories. Cogn. Affect. Behav. Neurosci. 6, 214–222. https://doi.org/10.3758/CABN.6.3.214

Goldberg, R.F., Perfetti, C.A., Schneider, W., 2006b. Perceptual Knowledge Retrieval Activates Sensory Brain Regions. J. Neurosci. 26, 4917–4921. https://doi.org/10.1523/JNEUROSCI.5389-05.2006

González, J., Barros-Loscertales, A., Pulvermüller, F., Meseguer, V., Sanjuán, A., Belloch, V., Ávila, C., 2006. Reading cinnamon activates olfactory brain regions. Neuroimage 32, 906–912. https://doi.org/10.1016/j.neuroimage.2006.03.037

Gottfried, J.A., Dolan, R.J., 2003. The nose smells what the eye sees: Crossmodal visual facilitation of human olfactory perception. Neuron 39, 375–386. https://doi.org/10.1016/S0896-6273(03)00392-1

Grossman, M., Koenig, P., DeVita, C., Glosser, G., Alsop, D., Detre, J., Gee, J., 2002a. The Neural Basis for Category-Specific Knowledge: An fMRI Study. Neuroimage 15, 936–948. https://doi.org/10.1006/nimg.2001.1028

Grossman, M., Koenig, P., DeVita, C., Glosser, G., Alsop, D., Detre, J., Gee, J., 2002b. Neural representation of verb meaning: An fMRI study. Hum. Brain Mapp. 15, 124–134. https://doi.org/10.1002/hbm.10117

Halpern, A.R., Zatorre, R.J., 1999. When that tune runs through your head: A PET investigation of auditory imagery for familiar melodies. Cereb. Cortex 9, 697–704. https://doi.org/10.1093/cercor/9.7.697

Halpern, A.R., Zatorre, R.J., Bouffard, M., Johnson, J.A., 2004. Behavioral and neural correlates of perceived and imagined musical timbre. Neuropsychologia 42, 1281–1292. https://doi.org/10.1016/j.neuropsychologia.2003.12.017

Hauk, O., Davis, M.H., Kherif, F., Pulvermüller, F., 2008. Imagery or meaning? Evidence for a semantic origin of category-specific brain activity in metabolic imaging. Eur. J. Neurosci. 27, 1856–1866. https://doi.org/10.1111/j.1460-9568.2008.06143.x

Hellbernd, N., Sammler, D., 2018. Neural bases of social communicative intentions in speech. Soc. Cogn. Affect. Neurosci. 13, 604–615. https://doi.org/10.1093/scan/nsy034

Herz, R.S., Eliassen, J., Beland, S., Souza, T., 2004. Neuroimaging evidence for the emotional potency of odor-evoked memory. Neuropsychologia 42, 371–378. https://doi.org/10.1016/j.neuropsychologia.2003.08.009

Hoenig, K., Müller, C., Herrnberger, B., Sim, E.-J., Spitzer, M., Ehret, G., Kiefer, M., 2011. Neuroplasticity of semantic representations for musical instruments in professional musicians. Neuroimage 56, 1714–1725. https://doi.org/10.1016/j.neuroimage.2011.02.065

Hoenig, K., Sim, E.-J., Bochev, V., Herrnberger, B., Kiefer, M., 2008. Conceptual Flexibility in the Human Brain: Dynamic Recruitment of Semantic Maps from Visual, Motor, and Motion-related Areas. J. Cogn. Neurosci. 20, 1799–1814. https://doi.org/10.1162/jocn.2008.20123

Hoeren, M., Kaller, C.P., Glauche, V., Vry, M.S., Rijntjes, M., Hamzei, F., Weiller, C., 2013. Action semantics and movement characteristics engage distinct processing streams during the observation of tool use. Exp. Brain Res. 229, 243–260. https://doi.org/10.1007/s00221-013-3610-5

Hofstetter, C., Achaibou, A., Vuilleumier, P., 2012. Reactivation of visual cortex during memory retrieval: Content specificity and emotional modulation. Neuroimage 60, 1734–1745. https://doi.org/10.1016/j.neuroimage.2012.01.110

Howard, R.J., Ffytche, D.H., Barnes, J., McKeefry, D., Ha, Y., Woodruff, P.W., Bullmore, E.T., Simmons, A., Williams, S.C.R., David, A.S., Brammer, M., 1998. The functional anatomy of imagining and perceiving colour. Neuroreport 9, 1019–1023. https://doi.org/10.1097/00001756-199804200-00012

Hsu, N.S., Kraemer, D.J.M., Oliver, R.T., Schlichting, M.L., Thompson-Schill, S.L., 2011. Color, context, and cognitive style: variations in color knowledge retrieval as a function of task and subject variables. J. Cogn. Neurosci. 23, 2544–57. https://doi.org/10.1162/jocn.2011.21619

Humphreys, G.F., Newling, K., Jennings, C., Gennari, S.P., 2013. Motion and actions in language: Semantic representations in occipito-temporal cortex. Brain Lang. 125, 94–105. https://doi.org/10.1016/j.bandl.2013.01.008

Isenberg, N., Silbersweig, D., Engelien, A., Emmerich, S., Malavade, K., Beattie, B., Leon, A.C., Stern, E., 1999. Linguistic threat activates the human amygdala. Proc. Natl. Acad. Sci. U. S. A. 96, 10456–10459. https://doi.org/10.1073/pnas.96.18.10456

Ishai, A., Ungerleider, L.G., Haxby, J. V., 2000. Distributed neural systems for the generation of visual images. Neuron 28, 979–990. https://doi.org/10.1016/S0896-6273(00)00168-9

Jabbi, M., Swart, M., Keysers, C., 2007. Empathy for positive and negative emotions in the gustatory cortex. Neuroimage 34, 1744–1753. https://doi.org/10.1016/j.neuroimage.2006.10.032

Johnson-Frey, S.H., Newman-Norlund, R., Grafton, S.T., 2005. A distributed left hemisphere network active during planning of everyday tool use skills. Cereb. Cortex 15, 681–695. https://doi.org/10.1093/cercor/bhh169

Just, M.A., Cherkassky, V.L., Aryal, S., Mitchell, T.M., 2010. A neurosemantic theory of concrete noun representation based on the underlying brain codes. PLoS One 5. https://doi.org/10.1371/journal.pone.0008622

Kable, J.W., Chatterjee, A., 2006. Specificity of Action Representations in the Lateral Occipitotemporal Cortex. J. Cogn. Neurosci. 18, 1498–1517. https://doi.org/10.1162/jocn.2006.18.9.1498

Kana, R.K., Blum, E.R., Ladden, S.L., Ver Hoef, L.W., 2012. “How to do things with Words”: Role of motor cortex in semantic representation of action words. Neuropsychologia 50, 3403–3409. https://doi.org/10.1016/j.neuropsychologia.2012.09.006

Kedia, G., Berthoz, S., Wessa, M., Hilton, D., Martinot, J.-L., 2008. An agent harms a victim: an fMRI study on specific moral emotions. J. Cogn. Neurosci. 20, 1788–98.

Kellenbach, M.L., Brett, M., Patterson, K., 2003. Actions Speak Louder Than Functions: The Importance of Manipulability and Action in Tool Representation. J. Cogn. Neurosci. 15, 30–46. https://doi.org/10.1162/089892903321107800

Kellenbach, M.L., Brett, M., Patterson, K., 2001. Large, colorful, or noisy? Attribute- and modality-specific activations during retrieval of perceptual attribute knowledge. Cogn. Affect. Behav. Neurosci. 1, 207–221. https://doi.org/10.3758/CABN.1.3.207

Kemmerer, D., Castillo, J.G., Talavage, T., Patterson, S., Wiley, C., 2008. Neuroanatomical distribution of five semantic components of verbs: Evidence from fMRI. Brain Lang. 107, 16–43. https://doi.org/10.1016/j.bandl.2007.09.003

Khader, P.H., Jost, K., Mertens, M., Bien, S., Rösler, F., 2010. Neural correlates of generating visual nouns and motor verbs in a minimal phrase context. Brain Res. 1318, 122–132. https://doi.org/10.1016/j.brainres.2009.12.082

Kiefer, M., Sim, E.-J., Herrnberger, B., Grothe, J., Hoenig, K., 2008. The Sound of Concepts: Four Markers for a Link between Auditory and Conceptual Brain Systems. J. Neurosci. 28, 12224–12230. https://doi.org/10.1523/JNEUROSCI.3579-08.2008

Kleineberg, N.N., Dovern, A., Binder, E., Grefkes, C., Eickhoff, S.B., Fink, G.R., Weiss, P.H., 2018. Action and semantic tool knowledge – Effective connectivity in the underlying neural networks. Hum. Brain Mapp. 39, 3473–3486. https://doi.org/10.1002/hbm.24188

Knutson, K.M., Wood, J.N., Spampinato, M. V., Grafman, J., 2006. Politics on the brain: An fMRI investigation. Soc. Neurosci. 1, 25–40. https://doi.org/10.1080/17470910600670603

Kobayashi, M., Takeda, M., Hattori, N., Fukunaga, M., Sasabe, T., Inoue, N., Nagai, Y., Sawada, T., Sadato, N., Watanabe, Y., 2004. Functional imaging of gustatory perception and imagery: "Top-down" processing of gustatory signals. Neuroimage 23, 1271–1282. https://doi.org/10.1016/j.neuroimage.2004.08.002

Kosslyn, S.M., Thompson, W.L., Klm, I.J., Alpert, N.M., 1995. Topographical representations of mental images in primary visual cortex. Nature 378, 496–498. https://doi.org/10.1038/378496a0

Kuchinke, L., Jacobs, A.M., Grubich, C., Võ, M.L.H., Conrad, M., Herrmann, M., 2005. Incidental effects of emotional valence in single word processing: An fMRI study. Neuroimage 28, 1022–1032. https://doi.org/10.1016/j.neuroimage.2005.06.050

Kuhnke, P., Kiefer, M., Hartwigsen, G., 2020. Task-Dependent Recruitment of Modality-Specific and Multimodal Regions during Conceptual Processing. Cereb. Cortex 30, 3938–3959. https://doi.org/10.1093/cercor/bhaa010

Lai, V.T., Desai, R.H., 2016. The grounding of temporal metaphors. Cortex 76, 43–50. https://doi.org/10.1016/j.cortex.2015.12.007

Lin, N., Bi, Y., Zhao, Y., Luo, C., Li, X., 2015. The theory-of-mind network in support of action verb comprehension: Evidence from an fMRI study. Brain Lang. 141, 1–10. https://doi.org/10.1016/j.bandl.2014.11.004

Luo, Q., Peng, D., Jin, Z., Xu, D., Xiao, L., Ding, G., 2004. Emotional valence of words modulates the subliminal repetition priming effect in the left fusiform gyrus: An event-related fMRI study. Neuroimage 21, 414–421. https://doi.org/10.1016/j.neuroimage.2003.09.048

Macdonald, S.N., Culham, J.C., 2015. Do human brain areas involved in visuomotor actions show a preference for real tools over visually similar non-tools? Neuropsychologia 77, 35–41. https://doi.org/10.1016/j.neuropsychologia.2015.08.004

Maddock, R.J., Garrett, A.S., Buonocore, M.H., 2003. Posterior cingulate cortex activation by emotional words: fMRI evidence from a valence decision task. Hum. Brain Mapp. 18, 30–41. https://doi.org/10.1002/hbm.10075

Maratos, E.J., Dolan, R.J., Morris, J.S., Henson, R.N.A., Rugg, M.D., 2001. Neural activity associated with episodic memory for emotional context. Neuropsychologia 39, 910–920. https://doi.org/10.1016/S0028-3932(01)00025-2

Martin, A., Haxby, J. V., Lalonde, F.M., Wiggs, C.L., Ungerleider, L.G., 1995. Discrete cortical regions associated with knowledge of color and knowledge of action. Science (80-. ). 270, 102–105. https://doi.org/10.1126/science.270.5233.102

Martin, A., Wiggs, C.L., Ungerleider, L.G., Haxby, J. V, 1996. Neural correlates of category-specific knowledge. Nature 379, 649–652. https://doi.org/10.1038/379649a0

Mellet, E., Tzourio, N., Denis, M., Mazoyer, B., 1998. Cortical anatomy of mental imagery of concrete nouns based on their dictionary definition. Neuroreport 9, 803–808. https://doi.org/10.1016/S1053-8119(18)30958-3

Michl, P., Meindl, T., Meister, F., Born, C., Engel, R.R., Reiser, M., Hennig-Fast, K., 2014. Neurobiological underpinnings of shame and guilt: A pilot fMRI study. Soc. Cogn. Affect. Neurosci. 9, 150–157. https://doi.org/10.1093/scan/nss114

Moll, J., de Oliveira-Souza, R., Garrido, G.J., Bramati, I.E., Caparelli-Daquer, E.M.A., Paiva, M.L.M.F., Zahn, R., Grafman, J., 2007. The self as a moral agent: Linking the neural bases of social agency and moral sensitivity. Soc. Neurosci. 2, 336–352. https://doi.org/10.1080/17470910701392024

Moseley, R., Carota, F., Hauk, O., Mohr, B., Pulvermüller, F., 2012. A role for the motor system in binding abstract emotional meaning. Cereb. Cortex 22, 1634–1647. https://doi.org/10.1093/cercor/bhr238

Mummery, C.J., Patterson, K., Hodges, J.R., Price, C.J., 1998. Functional neuroanatomy of the semantic system: Divisible by what? J. Cogn. Neurosci. 10, 766–777. https://doi.org/10.1162/089892998563059

Nagels, A., Chatterjee, A., Kircher, T., Straube, B., 2013. The role of semantic abstractness and perceptual category in processing speech accompanied by gestures. Front. Behav. Neurosci. 7, 1–12. https://doi.org/10.3389/fnbeh.2013.00181

Neudorf, J., Ekstrand, C., Kress, S., Borowsky, R., 2019. FMRI of shared-stream priming of lexical identification by object semantics along the ventral visual processing stream. Neuropsychologia 133, 107185. https://doi.org/10.1016/j.neuropsychologia.2019.107185

Nijhof, A.D., Willems, R.M., 2015. Simulating fiction: Individual differences in literature comprehension revealed with fMRI. PLoS One 10, 7–11. https://doi.org/10.1371/journal.pone.0116492

Noppeney, U., Josephs, O., Kiebel, S., Friston, K.J., Price, C.J., 2005. Action selectivity in parietal and temporal cortex. Cogn. Brain Res. 25, 641–649. https://doi.org/10.1016/j.cogbrainres.2005.08.017

Noppeney, U., Price, C.J., 2002. Retrieval of visual, auditory, and abstract semantics. Neuroimage 15, 917–926. https://doi.org/10.1006/nimg.2001.1016

Noppeney, U., Price, C.J., Penny, W.D., Friston, K.J., 2006. Two Distinct Neural Mechanisms for Category-selective Responses. Cereb. Cortex 16, 437–445. https://doi.org/10.1093/cercor/bhi123

Oosterwijk, S., Mackey, S., Wilson-Mendenhall, C., Winkielman, P., Paulus, M.P., 2015. Concepts in context: Processing mental state concepts with internal or external focus involves different neural systems. Soc. Neurosci. 10, 294–307. https://doi.org/10.1080/17470919.2014.998840

Peelen, M. V., Bracci, S., Lu, X., He, C., Caramazza, A., Bi, Y., 2013. Tool Selectivity in Left Occipitotemporal Cortex Develops without Vision. J. Cogn. Neurosci. 25, 1225–1234. https://doi.org/10.1162/jocn_a_00411

Peelen, M. V., Romagno, D., Caramazza, A., 2012. Independent Representations of Verbs and Actions in Left Lateral Temporal Cortex. J. Cogn. Neurosci. 24, 2096–2107. https://doi.org/10.1162/jocn_a_00257

Phillips, J.A., Noppeney, U., Humphreys, G.W., Price, C.J., 2002. Can segregation within the semantic system account for category-specific deficits? Brain 125, 2067–2080. https://doi.org/10.1093/brain/awf215

Phillips, M.L., Young, A.W., Scott, S.K., Calder, A.J., Andrew, C., Giampietro, V., Williams, S.C.R., Bullmore, E.T., Brammer, M., Gray, J.A., 1998. Neural responses to facial and vocal expressions of fear and disgust. Proc. R. Soc. B Biol. Sci. 265. https://doi.org/10.1098/rspb.1998.0506

Pomp, J., Bestgen, A.K., Schulze, P., Müller, C.J., Citron, F.M.M., Suchan, B., Kuchinke, L., 2018. Lexical olfaction recruits olfactory orbitofrontal cortex in metaphorical and literal contexts. Brain Lang. 179, 11–21. https://doi.org/10.1016/j.bandl.2018.02.001

Popp, M., Trumpp, N.M., Kiefer, M., 2019a. Processing of action and sound verbs in context: An fMRI study. Transl. Neurosci. 10, 200–222. https://doi.org/10.1515/tnsci-2019-0035

Popp, M., Trumpp, N.M., Sim, E., Kiefer, M., 2019b. Brain Activation During Conceptual Processing of Action and Sound Verbs. Adv. Cogn. Psychol. 15, 236–255. https://doi.org/10.5709/acp-0272-4

Pulvermüller, F., Hauk, O., 2006. Category-specific Conceptual Processing of Color and Form in Left Fronto-temporal Cortex. Cereb. Cortex 16, 1193–1201. https://doi.org/10.1093/cercor/bhj060

Raposo, A., Moss, H.E., Stamatakis, E.A., Tyler, L.K., 2009. Modulation of motor and premotor cortices by actions, action words and action sentences. Neuropsychologia 47, 388–396. https://doi.org/10.1016/j.neuropsychologia.2008.09.017

Rodríguez-Ferreiro, J., Gennari, S.P., Davies, R., Cuetos, F., 2011. Neural correlates of abstract verb processing. J. Cogn. Neurosci. 23, 106–118. https://doi.org/10.1162/jocn.2010.21414

Romero Lauro, L.J., Mattavelli, G., Papagno, C., Tettamanti, M., 2013. She runs, the road runs, my mind runs, bad blood runs between us: Literal and figurative motion verbs: An fMRI study. Neuroimage 83, 361–371. https://doi.org/10.1016/j.neuroimage.2013.06.050

Ruby, P., Decety, J., 2004. How would You feel versus how do you think She would feel? A neuroimaging study of perspective-taking with social emotions. J. Cogn. Neurosci. 16, 988–999. https://doi.org/10.1162/0898929041502661

Ruby, P., Decety, J., 2001. Effect of subjective perspective taking during simulation of action: A PET investigation of agency. Nat. Neurosci. 4, 546–550. https://doi.org/10.1038/87510

Rueschemeyer, S.-A., Brass, M., Friederici, A.D., 2007. Comprehending Prehending: Neural Correlates of Processing Verbs with Motor Stems. J. Cogn. Neurosci. 19, 855–865. https://doi.org/10.1162/jocn.2007.19.5.855

Rueschemeyer, S.-A., van Rooij, D., Lindemann, O., Willems, R.M., Bekkering, H., 2010. The function of words: distinct neural correlates for words denoting differently manipulable objects. J. Cogn. Neurosci. 22, 1844–51. https://doi.org/10.1162/jocn.2009.21310

Savic, I., Berglund, H., 2004. Passive perception of odors and semantic circuits. Hum. Brain Mapp. 21, 271–278. https://doi.org/10.1002/hbm.20009

Saxbe, D.E., Yang, X.F., Borofsky, L.A., Immordino-Yang, M.H., 2013. The embodiment of emotion: Language use during the feeling of social emotions predicts cortical somatosensory activity. Soc. Cogn. Affect. Neurosci. 8, 806–812. https://doi.org/10.1093/scan/nss075

Schuil, K.D.I., Smits, M., Zwaan, R.A., 2013. Sentential Context Modulates the Involvement of the Motor Cortex in Action Language Processing: An fMRI Study. Front. Hum. Neurosci. 7, 1–13. https://doi.org/10.3389/fnhum.2013.00100

Simmons, W.K., Martin, A., Barsalou, L.W., 2005. Pictures of appetizing foods activate gustatory cortices for taste and reward. Cereb. Cortex 15, 1602–1608. https://doi.org/10.1093/cercor/bhi038

Skipper, L.M., Olson, I.R., 2014. Semantic memory: Distinct neural representations for abstractness and valence. Brain Lang. 130, 1–10. https://doi.org/10.1016/j.bandl.2014.01.001

Skipper, L.M., Ross, L.A., Olson, I.R., 2011. Sensory and semantic category subdivisions within the anterior temporal lobes. Neuropsychologia 49, 3419–3429. https://doi.org/10.1016/j.neuropsychologia.2011.07.033

Slotnick, S.D., 2009. Memory for color reactivates color processing region. Neuroreport 20, 1568–1571. https://doi.org/10.1097/WNR.0b013e328332d35e

Smith, A.P.R., Henson, R.N.A., Dolan, R.J., Rugg, M.D., 2004. fMRI correlates of the episodic retrieval of emotional contexts. Neuroimage 22, 868–878. https://doi.org/10.1016/j.neuroimage.2004.01.049

Speer, N.K., Reynolds, J.R., Swallow, K.M., Zacks, J.M., 2009. Reading stories activations neural representations of visual and motor experiences. Psychol. Sci. 20, 289–299.

Spunt, R.P., Lieberman, M.D., 2012. Dissociating Modality-Specific and Supramodal Neural Systems for Action Understanding. J. Neurosci. 32, 3575–3583. https://doi.org/10.1523/JNEUROSCI.5715-11.2012

Steinbeis, N., Koelsch, S., 2008. Comparing the processing of music and language meaning using EEG and fMRI provides evidence for similar and distinct neural representations. PLoS One 3, 1–7. https://doi.org/10.1371/journal.pone.0002226

Strange, B.A., Henson, R.N.A., Friston, K.J., Dolan, R.J., 2000. Brain mechanisms for detecting perceptual, semantic, and emotional deviance. Neuroimage 12, 425–433. https://doi.org/10.1006/nimg.2000.0637

Takahashi, H., Yahata, N., Koeda, M., Matsuda, T., Asai, K., Okubo, Y., 2004. Brain activation associated with evaluative processes of guilt and embarrassment: An fMRI study. Neuroimage 23, 967–974. https://doi.org/10.1016/j.neuroimage.2004.07.054

Tavares, P., Barnard, P.J., Lawrence, A.D., 2011. Emotional complexity and the neural representation of emotion in motion. Soc. Cogn. Affect. Neurosci. 6, 98–108. https://doi.org/10.1093/scan/nsq021

Tettamanti, M., Buccino, G., Saccuman, M.C., Gallese, V., Danna, M., Scifo, P., Fazio, F., Rizzolatti, G., Cappa, S.F., Perani, D., 2005. Listening to Action-related Sentences Activates Fronto-parietal Motor Circuits. J. Cogn. Neurosci. 17, 273–281. https://doi.org/10.1162/0898929053124965

Tettamanti, M., Manenti, R., Della Rosa, P.A., Falini, A., Perani, D., Cappa, S.F., Moro, A., 2008. Negation in the brain: Modulating action representations. Neuroimage 43, 358–367. https://doi.org/10.1016/j.neuroimage.2008.08.004

Tettamanti, M., Rognoni, E., Cafiero, R., Costa, T., Galati, D., Perani, D., 2012. Distinct pathways of neural coupling for different basic emotions. Neuroimage 59, 1804–1817. https://doi.org/10.1016/j.neuroimage.2011.08.018

Thompson-Schill, S.L., Aguirre, G.K., D'Esposito, M., Farah, M.J., 1999. A neural basis for category and modality specificity of semantic knowledge. Neuropsychologia 37, 671–676. https://doi.org/10.1016/S0028-3932(98)00126-2

Tomasino, B., Fabbro, F., Brambilla, P., 2014. How do conceptual representations interact with processing demands: An fMRI study on action- and abstract-related words. Brain Res. 1591, 38–52. https://doi.org/10.1016/j.brainres.2014.10.008

Tomasino, B., Werner, C.J., Weiss, P.H., Fink, G.R., 2007. Stimulus properties matter more than perspective: An fMRI study of mental imagery and silent reading of action phrases. Neuroimage 36, T128–T141. https://doi.org/10.1016/j.neuroimage.2007.03.035

van Ackeren, M.J., Smaragdi, A., Rueschemeyer, S.A., 2016. Neuronal interactions between mentalising and action systems during indirect request processing. Soc. Cogn. Affect. Neurosci. 11, 1402–1410. https://doi.org/10.1093/scan/nsw062

van Dam, W.O., Almor, A., Shinkareva, S. V., Kim, J., Boiteau, T.W., Shay, E.A., Desai, R.H., 2019. Distinct neural mechanisms underlying conceptual knowledge of manner and instrument verbs. Neuropsychologia 133, 107183. https://doi.org/10.1016/j.neuropsychologia.2019.107183

van Dam, W.O., Desai, R.H., 2016. The Semantics of Syntax: The Grounding of Transitive and Intransitive Constructions. J. Cogn. Neurosci. 28, 693–709. https://doi.org/10.1162/jocn_a_00926

van Dam, W.O., Rueschemeyer, S.A., Bekkering, H., 2010. How specifically are action verbs represented in the neural motor system: An fMRI study. Neuroimage 53, 1318–1325. https://doi.org/10.1016/j.neuroimage.2010.06.071

Vigliocco, G., Kousta, S.T., Della Rosa, P.A., Vinson, D.P., Tettamanti, M., Devlin, J.T., Cappa, S.F., 2014. The neural representation of abstract words: The role of emotion. Cereb. Cortex 24, 1767–1777. https://doi.org/10.1093/cercor/bht025

Vigliocco, G., Warren, J., Siri, S., Arciuli, J., Scott, S., Wise, R., 2006. The role of semantics and grammatical class in the neural representation of words. Cereb. Cortex 16, 1790–1796. https://doi.org/10.1093/cercor/bhj115

Wallentin, M., Lund, T.E., Østergaard, S., Oøstergaard, L., Roepstorff, A., 2005. Motion verb sentences activate left posterior middle temporal cortex despite static context. Neuroreport 16, 649–652. https://doi.org/10.1097/00001756-200504250-00027

Wallentin, M., Nielsen, A.H., Vuust, P., Dohn, A., Roepstorff, A., Lund, T.E., 2011. BOLD response to motion verbs in left posterior middle temporal gyrus during story comprehension. Brain Lang. 119, 221–225. https://doi.org/10.1016/j.bandl.2011.04.006

Wang, X., Han, Z., He, Y., Caramazza, A., Song, L., Bi, Y., 2013. Where color rests: Spontaneous brain activity of bilateral fusiform and lingual regions predicts object color knowledge performance. Neuroimage 76, 252–263. https://doi.org/10.1016/j.neuroimage.2013.03.010

Wang, X., Wang, B., Bi, Y., 2019. Close yet independent: Dissociation of social from valence and abstract semantic dimensions in the left anterior temporal lobe. Hum. Brain Mapp. 40, 4759–4776. https://doi.org/10.1002/hbm.24735

Warburton, E., Wise, R.J.S., Price, C.J., Weiller, C., Hadar, U., Ramsay, S., Frackowiak, R.S.J., 1996. Noun and verb retrieval by normal subjects: Studies with PET. Brain 119, 159–179. https://doi.org/10.1093/brain/119.1.159

Webster, P.J., Skipper-Kallal, L.M., Frum, C.A., Still, H.N., Ward, B.D., Lewis, J.W., 2017. Divergent human cortical regions for processing distinct acoustic-semantic categories of natural sounds: Animal action sounds vs. vocalizations. Front. Neurosci. 10, 1–18. https://doi.org/10.3389/fnins.2016.00579

Wheatley, T., Weisberg, J., Beauchamp, M.S., Martin, A., 2005. Automatic Priming of Semantically Related Words Reduces Activity in the Fusiform Gyrus. J. Cogn. Neurosci. 17, 1871–1885. https://doi.org/10.1162/089892905775008689

Wheeler, M.E., Petersen, S.E., Buckner, R.L., 2000. Memory's echo: Vivid remembering reactivates sensory-specific cortex. Proc. Natl. Acad. Sci. 97, 11125–11129. https://doi.org/10.1073/pnas.97.20.11125

Wicker, B., Keysers, C., Plailly, J., Royet, J.P., Gallese, V., Rizzolatti, G., 2003. Both of us disgusted in My insula: The common neural basis of seeing and feeling disgust. Neuron 40, 655–664. https://doi.org/10.1016/S0896-6273(03)00679-2

Willems, R.M., Toni, I., Hagoort, P., Casasanto, D., 2010. Neural dissociations between action verb understanding and motor imagery. J. Cogn. Neurosci. 22, 2387–2400. https://doi.org/10.1162/jocn.2009.21386

Wilson-Mendenhall, C.D., Barrett, L.F., Simmons, W.K., Barsalou, L.W., 2011. Grounding emotion in situated conceptualization. Neuropsychologia 49, 1105–1127. https://doi.org/10.1016/j.neuropsychologia.2010.12.032

Wriessnegger, S.C., Steyrl, D., Koschutnig, K., Müller-Putz, G.R., 2016. Cooperation in mind: Motor imagery of joint and single actions is represented in different brain areas. Brain Cogn. 109, 19–25. https://doi.org/10.1016/j.bandc.2016.08.008

Yang, J., Shu, H., 2014. Passive reading and motor imagery about hand actions and tool-use actions: An fMRI study. Exp. Brain Res. 232, 453–467. https://doi.org/10.1007/s00221-013-3753-4

Yoo, S.S., Lee, C.U., Choi, B.G., 2001. Human brain mapping of auditory imagery: Event-related functional MRI study. Neuroreport 12, 3045–3049. https://doi.org/10.1097/00001756-200110080-00013

Zatorre, R.J., Halpern, A.R., Perry, D.W., Meyer, E., Evans, A.C., 1996. Hearing in the Mind's Ear: A PET Investigation of Musical Imagery and Perception. J. Cogn. Neurosci. 8, 29–46. https://doi.org/10.1162/jocn.1996.8.1.29

Zvyagintsev, M., Clemens, B., Chechko, N., Mathiak, K.A., Sack, A.T., Mathiak, K., 2013. Brain networks underlying mental imagery of auditory and visual information. Eur. J. Neurosci. 37, 1421–1434. https://doi.org/10.1111/ejn.12140

Zysset, S., Huber, O., Ferstl, E., Von Cramon, D.Y., 2002. The anterior frontomedian cortex and evaluative judgment: An fMRI study. Neuroimage 15, 983–991. https://doi.org/10.1006/nimg.2001.1008

Zysset, S., Huber, O., Samson, A., Ferstl, E.C., Von Cramon, D.Y., 2003. Functional specialization within the anterior medial prefrontal cortex: A functional magnetic resonance imaging study with human subjects. Neurosci. Lett. 335, 183–186. https://doi.org/10.1016/S0304-3940(02)01196-5
